# The transcription factor network of *E. coli* steers global responses to shifts in RNAP concentration

**DOI:** 10.1101/2022.03.07.483226

**Authors:** Bilena L B Almeida, Mohamed N M Bahrudeen, Vatsala Chauhan, Suchintak Dash, Vinodh Kandavalli, Antti Häkkinen, Jason Lloyd-Price, Cristina S D Palma, Ines S C Baptista, Abhishekh Gupta, Juha Kesseli, Eric Dufour, Olli-Pekka Smolander, Matti Nykter, Petri Auvinen, Howard T Jacobs, Samuel M D Oliveira, Andre S Ribeiro

## Abstract

The robustness and sensitivity of gene networks to environmental changes is critical for cell survival. How gene networks produce specific, chronologically ordered responses to genome-wide perturbations, while robustly maintaining homeostasis, remains an open question. We analysed if short- and mid-term genome-wide responses to shifts in RNA polymerase (RNAP) concentration are influenced by the *known* topology and logic of the transcription factor network (TFN) of *Escherichia coli*. We found that, at the gene cohort level, the magnitude of the single-gene, mid-term transcriptional responses to changes in RNAP concentration can be explained by the absolute difference between the gene’s numbers of activating and repressing input transcription factors (TFs). Interestingly, this difference is strongly positively correlated with the number of input TFs of the gene. Meanwhile, short-term responses showed only weak influence from the TFN. Our results suggest that the global topological traits of the TFN of *E. coli* shape which gene cohorts respond to genome-wide stresses.

## INTRODUCTION

Gene regulatory networks (GRNs) receive, process, act upon, and send out information, while being robust to random fluctuations. How signals targeting one to a few genes are processed is relatively well understood (1,2). Meanwhile, many cellular environments fluctuate (sometimes unpredictably) in nutrient availability, pH, temperature, salts, community of other cells or species they live with, etc., which may cause genome-wide stresses. We investigate how GRNs produce chronologically ordered responses to genome-wide perturbations, while robustly maintaining homeostasis.

Evidence suggests that genome-wide stresses initially perturb hundreds to thousands of genes (3) but are quickly processed. As a result, after a transient period, only specific gene cohorts of tens to a few hundred genes (4,5) (usually sharing common feature(s)) participate in the responsive short-, mid- and long-term transcriptional programs (6). For example, when *Escherichia coli* suffers a cold shock, a specific cohort exhibits a fast, short-term response (∼70 genes), while another has a longer-term response (∼35 genes), with the rest remaining relatively passive (7,8). Since cells exhibit predictable, temporally ordered, beneficial phenotypic changes, these response programs have likely been positively selected during evolution.

It has been shown that global regulators (9-12), DNA supercoiling (13) and small RNAs (14), among other, can select large cohorts of stress-specific, responsive genes. It was also reported that 60%-90% of *E. coli* genes respond to changing growth conditions following a constant global scaling factor (15). Further, there is evidence that the effects of RNA polymerase (and other global regulators) can be separate from the effects of input TFs during genome-wide responses, using fluorescent reporters and small circuits (16,17).

Nevertheless, establishing whether and how the topology and logic of transcription factor (TF) networks (TFN) affect genome-wide responses remains challenging (18), despite some successes (19-22). These responses are most likely controlled, since cells exhibit predictable, ordered responses to natural genome-wide stresses (e.g., shifts in temperature or in growth-medium composition). The evolved features of GRNs that facilitate such outcomes remain unidentified, but their large-scale topology should be a key player.

To investigate the influence of the topology and logic of transcription factor (TF) networks (TFN) on large transcriptional programs, we studied what occurs following genome-wide perturbations. For this, we considered that, in *E. coli*, the concentration of the key genome-wide regulator, the RNA Polymerase, RNAP, naturally differs with medium composition (23). Also, it is well established how transcription kinetics differ according to RNAP concentration at the single-gene level (24,25), from which it is to be expected that changes in the latter should have genome-wide effects. We thus increasingly diluted media to alter systematically and rapidly (26,27) the amount of RNAP (illustrated in Figure 1A), and measured the genome-wide, short- and mid-term changes in transcript abundances.

**Figure 1.**
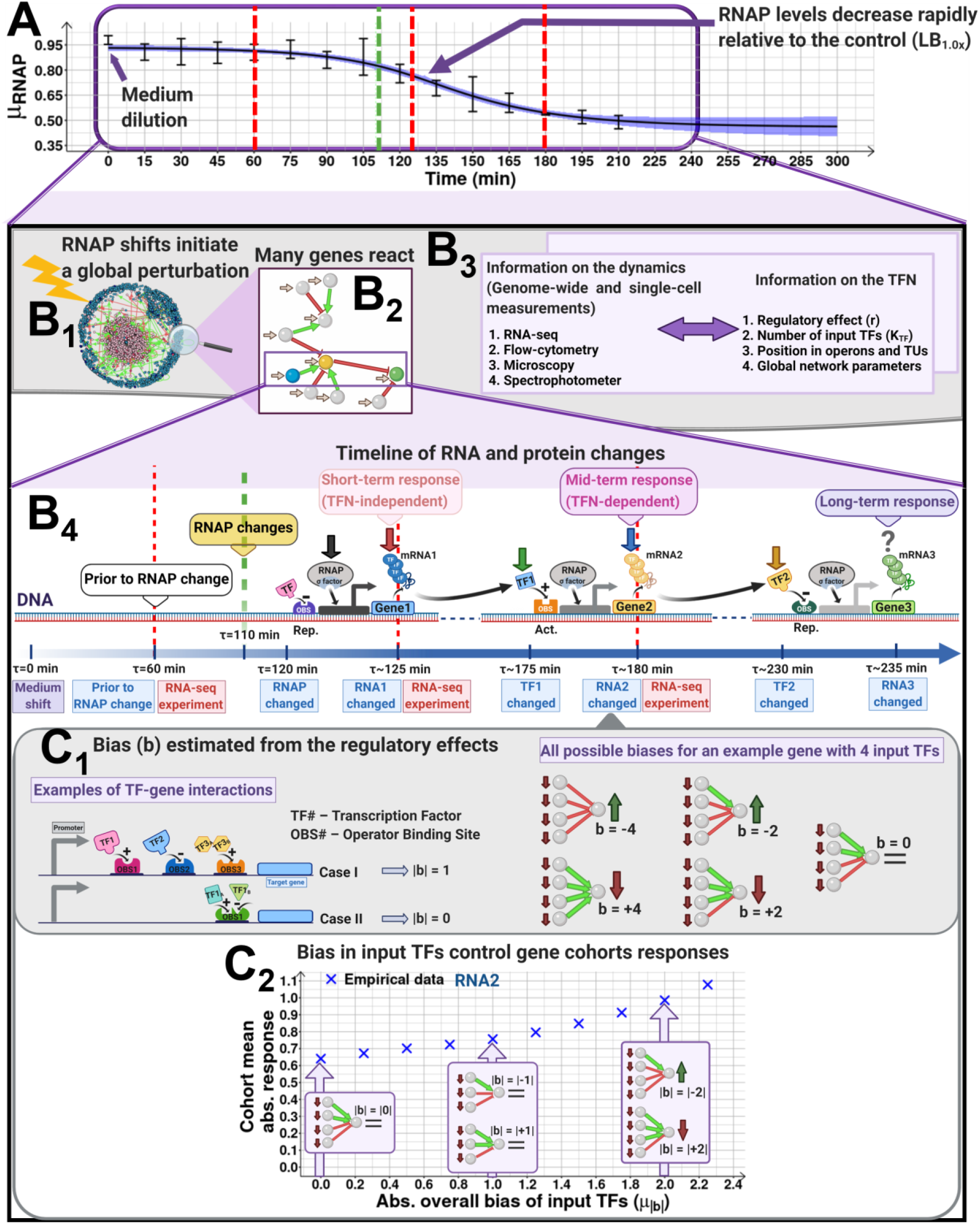
Expected short- and mid-term effects of quick downshifts of the RNAP amounts on the TFN of *E. coli*. **(A)** Example changes in mean RNAP (*μ*_*RNAP*_) and 68% CB (shadow) relative to control (LB_1.0x_) after diluting the medium (LB_0.5x_). Vertical red lines mark when the transcriptome measurements at 60 min, 125 min, and 180 min. Given the RNAP levels and the kinetics of RNA and protein abundances, these moments are named ‘prior to RNAP changes’ and ‘short-’, and ‘mid-term’ changes in RNA abundances. **(B**_**1**_) Known TF-gene interactions (red and green lines, if repressing and activating, respectively) and genes with (pink) and without (blue) input TFs of *E. coli*. **(B**_**2**_) Illustration of the effects of a local topology of activating (green) and repressing (red) input TFs on mid-term responses. Genes (balls) are coloured (blue, yellow, and green) according to the events in B_4_. **(B**_**3**_) Data collected on the genome-wide kinetics as well as data collected on the TFN structure. **(B**_**4**_) Following a medium dilution, intracellular RNAP concentrations (black arrow) decrease after a time lag, and RNA abundances (red arrow) will decrease accordingly. Compared to when at ∼0 min, the RNAP at ∼120 min and corresponding RNAs at ∼125 min should be lower (25,28). Given translation times (∼50 min (29-31)), at ∼175 min, the protein abundances, including input TFs, coded by the perturbed RNAs (green arrow) should differ as well. Fluctuations in these input TFs abundances will then propagate to nearest neighbour ‘output’ genes, further shifting their RNA abundances (blue arrow) depending on whether the input TF is an activator or a repressor. Finally, the yellow arrow represents (not measured) long-term changes (∼230 min or longer). We performed RNA-seq at ∼60 min (prior to RNAP changes), ∼125 min (short-term RNA changes), and ∼180 min (mid-term RNA changes, affected by input TFs). Finally, the green dashed line marks when the RNAP level already differs significantly from the control (see example Figure 1A). **(C**_**1**_) Illustration of biases in sets of input TFs of individual genes. Considering TF-gene interactions as either repressions (regulatory effect of -1) or activations (regulatory effect of +1), the overall effect of a set of input TFs during these stresses should be predictable from the sum of the input TFs regulatory effects, named ‘bias’, (*b*). Regulatory effects obtained from RegulonDB. **(C**_**2**_) Example average response (*μ*_|*LFC*|_ from RNA-seq) at 180 min of gene cohorts with a given *μ*_*b*_. Figures created with BioRender.com.

Decreases in RNAP levels should, in the short-term, cause quick genome-wide decreases in transcription rates, and thus in RNA abundances (Figure 1B_1_, Supplementary Results section *Expected effects of shifting RNA polymerase concentration on a gene’s transcription dynamics*). Such shifts, likely diverse in magnitudes, should then cause downshifts in the corresponding protein abundances. Thus, in the case of input TFs, their ‘output’ genes will, later on, be affected as well (Figure 1B_2_), causing further (here named ‘mid-term’) changes in their RNA abundances (Figure 1B_4_). Meanwhile, the short-term changes mostly likely are only affected by the genes’ individual features, since the protein abundances have not yet changed significantly in the cells.

We focused on the mid-term changes (measured by RNA-seq, Figures 1A and 1B_4_, dashed line). Specifically, we hypothesized that, genes will have their RNAs further decreased or, instead, increased, depending on whether their input TFs are activators or repressors, respectively. In detail, genes with multiple input TFs will have a mid-term response strength correlated to the difference between the numbers of its activator and repressor input TFs (Figures 1C_1_ and C_2_). This difference is here named ‘bias’ in the regulatory effects (activation or repression) of those input TFs of a gene.

At the single-gene level, we expect that the strength of the mid-term change in RNA abundances will be influenced by the strength of the shift in RNAP and input TFs concentrations, as well as by the specifics features of each gene and corresponding input TFs (bindings affinities, initiation kinetics, etc.). However, many of these features are largely unknown. As such, here we only studied empirically if the average responses of cohorts of genes can be explained by the mean bias in the regulatory effects of their input TFs, along with the strength of the shift in RNAP concentration (Figure 1C_2_). In detail, we interpret the data on the genome-wide kinetics based on the information on the TFN structure (logic and topology) (Figure 1B_3_).

*E. coli* was used to validate this hypothesis since its gene expression mechanisms have been largely dissected and the kinetics of transcription, translation, and RNA and protein degradation are well known (26,32,33). Also, its TFN is extensively mapped, with RegulonDB (34) informing on ∼4700 TF interactions between ∼4500 genes (and on their activating or repressing regulatory roles). Consequently, since we know the regulatory network *a priori*, instead of using the data on gene expression for network inference, we use it solely to quantify the genes’ responsiveness with respect to the TFN topology and logic. We then investigate whether the mid-term responsiveness to shifting RNAP concentrations is in accordance with the *presently known* topology and logic of the TFN, as hypothesized (Figures 1B_1_-1B_4_). Supplementary Table S1 has a description of the variables used throughout the manuscript.

## MATERIAL AND METHODS

### Bacterial strains, media, growth conditions and curves

We used wild type MG1655 cells as a base strain to study the transcriptome. In addition, we used an RL1314 strain with RpoC endogenously tagged with GFP (generously provided by Robert Landick) to measure RNAP levels, and 20 YFP fusion strains with genes endogenously tagged with the YFP coding sequence (25) to measure single-cell protein levels (Supplementary Table S4). Further, we used a strain carrying an rpoS::mCherry gene to measure RpoS levels (generously provided by James Locke), shown to track RpoS (35). In addition, we measured the protein expression levels of the spoT gene, which is one of the genes responsible for (p)ppGpp synthesis (3), using the YFP fusion library. Finally, we measured the single-cell levels of the crl gene using a low-copy plasmid fusion library of fluorescent (GFP) reporter strain (36).

From glycerol stocks (at - 80 °C), cells were streaked on lysogeny broth (LB) agar plates with antibiotics and kept at 37 °C overnight. Next, a single colony was picked, inoculated into fresh LB medium and, kept at 30 °C overnight with appropriate antibiotics and aeration at 250 rpm. From overnight cultures (ONC), cells were diluted to 1:1000 in tailored LB media (see below) with antibiotics, incubated at 37 °C with aeration, and allowed to grow until reaching an optical density of ≈ 0.4 at 600 nm (OD_600_).

Using this protocol, to attain cells with different intracellular RNAP concentration, starting from LB, we used tailored media, denoted as ‘LB_1.0x_’, ‘LB_0.75x_’, ‘LB_0.5x_’, ‘LB_0.25x_’, ‘LB_1.5x_’, ‘LB_2.0x_’ and ‘LB_2.5x_’ specifically, as in (26). Their composition for 100 ml (pH of 7.0) are, respectively: (LB_1.0x_) 1 g tryptone, 0.5 g yeast extract and 1 g NaCl; (LB_0.75x_) 0.75 g tryptone, 0.375 g yeast extract and 1 g NaCl; (LB_0.5x_) 0.5 g tryptone, 0.25 g yeast extract and 1 g NaCl; and (LB_0.25x_) 0.25 g tryptone, 0.125 g yeast extract and 1 g NaCl; (LB_1.5x_) 1.5 g tryptone, 0.75 g yeast extract and 1 g NaCl; (LB_2.0x_) 2 g tryptone, 1 g yeast extract and 1 g NaCl; (LB_2.5x_) 2.5 g tryptone, 1.25 g yeast extract and 1 g NaCl.

To measure cell growth curves and rates, ONC of the RL1314 strain were diluted to an initial optical density at 600 nm (OD_600_) of ≈ 0.05 into independent fresh media (LB_1.0x_, LB_0.75x_, LB_0.5x,_ LB_0.25x_, LB_1.5x_, LB_2.0x_ and LB_2.5x_). The cultures were aliquoted in a 24-well flat bottom transparent plate and incubated at 37 °C with continuous shaking in a Biotek Synergy HTX Multi-Mode Reader. Growth was monitored every 10 min for 10 hours.

### Microscopy

To measure single-cell RNAP levels, ONC RL1314 cells were pre-inoculated into LB_1.0x_, LB_0.75x_, LB_0.5x_ and LB_0.25x_ media. Upon reaching mid-exponential growth phase, cells were pelleted by quick centrifugation (10000 rpm for 1 min), and the supernatant was discarded. The pellet was re-suspended in 100 µl of the remaining medium. Next, 3 µl of cells were placed in between 2% agarose gel pad and a coverslip and imaged by confocal microscopy with a 100x objective (example images in Supplementary Figure S1). GFP fluorescence was measured with a 488 nm laser and a 514/30 nm emission filter. Phase-contrast images were simultaneously acquired. MG1655 cells were imaged to measure cell size in LB_1.0x_, LB_0.75x_, LB_0.5x_, LB_0.25x_, LB_1.5x_, LB_2.0x_ and LB_2.5x_ media. Finally, MG1655 cells was also imaged in LB_1.0x_ during stationary growth. Finally, we imaged cells of the YFP strain library to assess if their morphology and physiology were consistent with healthy cells during measurements.

### Flow-cytometry

We performed flow-cytometry of RL1314 cells to measure single-cell RNAP over time. ONC were diluted at 1:1000 into respective fresh media (LB_1.0x_, LB_0.75x_, LB_0.5x_ and LB_0.25x_) and grown as described in Methods section *Bacterial strains, media, and growth conditions and curves*. Flow-cytometry data was recorded every 30 min (3 biological replicates), up to 210 min. Data was also captured in the mid-exponential phase (at 180 min), in the media studied (LB_1.0x_, LB_0.75x_, LB_0.5x,_ LB_0.25x_, LB_1.5x_, LB_2.0x_ and LB_2.5x_), with 3 biological replicates each. We used a similar protocol to perform flow-cytometry of several strains of the YFP library (25) in LB_1.0x_ and LB_0.25x_ (3 biological replicates, Supplementary Table S4), including to measure single-cell SpoT levels in LB_1.0x_, LB_0.75x_, LB_0.5x_, LB_0.25x_, LB_1.5x_, LB_2.0x_ and LB_2.5x_ (3 biological replicates each).

Meanwhile, we measured single-cell levels of the crl gene in LB_0.5x_ at 0 and 180 min, using a strain from the GFP-promoter fusion library. Further, to measure rpoS levels, we performed flow-cytometry of cells of the MGmCherry strain in LB_1.0x_, LB_0.75x_, LB_0.5x_ and LB_0.25x_ during the exponential (180 min) and stationary growth phases (LB_1.0x_,14 hours after pre-inoculation). In these measurements, as well as the measurements above, we recorded FSC-H, SSC-H and Width, to be used as proxies for cell size and density (i.e., composition), as they are positively correlated with these features (37).

In addition, data from measurements of MG1655 cells were used to discount background fluorescence from cells of the MGmCherry and the YFP strains. Similarly, measurements of the W3110 strain were used to discount the background fluorescence from the RL1314 strain.

For performing flow-cytometry, 5 μl of cells were diluted in 1 ml of PBS, and vortexed. In each condition, 50000 events were recorded. Prior to the experiments, QC was performed as recommended by the manufacturer. Measurements were conducted using an ACEA NovoCyte Flow Cytometer (ACEA Biosciences Inc., San Diego, USA) equipped with yellow and blue lasers.

For detecting the GFP and YFP signals, we used the FITC channel (-H parameter) with 488 nm excitation, 530/30 nm emission, and 14 μl/min sample flow rate with a core diameter of 7.7 μm. PMT voltage was set to 550 for FITC and kept the same for all conditions. Similarly, to detect the mCherry sinal, we used PE-Texas Red channel (-H parameter) having an excitation of 561 nm and emission of 615/20 nm and sample flow rate of 14 μl/min, with a core diameter of 7.7 μm. PMT voltage was set to 584 for PE-Texas Red and kept the same for all conditions. To remove background signal from particles smaller than bacteria, the detection threshold was set to 5000. All events were collected by Novo Express software from ACEA Biosciences Inc.

### Protein isolation and western blotting

Western blotting was used to quantify relative RNAP levels of MG1655 cells (Supplementary Figure S2 and Supplementary Table S2). Briefly, cells were diluted from ONC into respective fresh media and incubated at 37 °C with aeration and grown until reaching an OD_600_ ≈ 0.4. Next, cells were harvested by centrifugation (8000 rpm for 5 min) and pellets were lysed with B-PER bacterial protein extraction reagent, added with a protease inhibitor for 10 min at room temperature (RT). Following lysis, centrifugation was done at 14000 rpm for 10 min and the supernatant was collected. Next, the supernatant was diluted in 4X Laemmli buffer with β-mercaptoethanol and samples were boiled at 95 °C for 5 min.

Samples with ∼30 μg of soluble total proteins were loaded on 4%-20% TGX stain-free precast gels (Biorad). These proteins were then separated by electrophoresis and transferred on PVDF membrane using TurboBlot (Biorad). Next, membranes were blocked with 5% non-fat milk at room temperature (RT) for 1 h and probed with primary RpoC (β prime subunit of RNAP) antibodies at 1:2000 dilutions (Biolegend) at 4 °C overnight. HRP-secondary antibody (1:5000) treatment was then done (Sigma Aldrich) for 1 h at RT. Excess antibodies were removed by washing with buffer. The membrane was treated with chemiluminescence reagent (Biorad) for band detection. Images were obtained by the Chemidoc XRS system (Biorad) and band quantification was done using the Image Lab software (v.5.2.1).

### RNA-seq

#### a. Sample preparation

RNA-seq was performed thrice, for decreasing [LB_0.75x_, LB_0.5x_, and LB_0.25x_, at 180 min; LB_0.5x_ at 60 and 125 min] and for increasing (LB_1.5x_, LB_2.0x_ and LB_2.5x_, at 180 min) medium richness relative to a control (LB_1.0x_) (an independent control was used for each three sets of conditions). Cells from 3 independent biological replicates of MG1655 in each modified medium were treated with RNA protect bacteria reagent (Qiagen, Germany), to prevent degradation of RNA, and their total RNA was extracted using RNeasy kit (Qiagen). RNA was treated twice with DNase (Turbo DNA-free kit, Ambion) and quantified using Qubit 2.0 Fluorometer RNA assay (Invitrogen, Carlsbad, CA, USA). Total RNA amounts were determined by gel electrophoresis, using a 1% agarose gel stained with SYBR safe (Invitrogen). RNA was detected using UV with a Chemidoc XRS imager (Biorad).

#### b. Sequencing

##### i) Part 1: For shifts from LB_1.0x_ to LB_0.75x_, LB_0.5x_, and LB_0.25x_, at 180 min

Sequencing was performed by Acobiom (Montpellier, France). The RNA integrity number (RIN) of the samples was obtained with the 2100 Bioanalyzer (Agilent Technologies, Palo Alto, USA) using Eukaryotic Total RNA 6000 Nano Chip (Agilent Technologies). Ribosomal RNA depletion was performed using Ribo-Zero removal kit (Bacteria) from Illumina. RNA-seq libraries were constructed according to the Illumina’s protocol. Samples were sequenced using a single-index, 1×75bp single-end configuration (∼10M reads/library) on an Illumina MiSeq instrument. Sequencing analysis and base calling were performed using the Illumina Pipeline. Sequences were obtained after purity filtering.

##### ii) Part 2: For shifts from LB_1.0x_ to LB_1.5x_, LB_2.0x_, and LB_2.5x_ at 180 min, and from LB_1.0x_ to LB_0.5x_ at 60 and 125 min

Sequencing was performed by GENEWIZ, Inc. (Leipzig, Germany). The RIN of the samples was obtained with the Agilent 4200 TapeStation (Agilent Technologies, Palo Alto, CA, USA). Ribosomal RNA depletion was performed using Ribo-Zero Gold Kit (Bacterial probe) (Illumina, San Diego, CA, USA). RNA-seq libraries were constructed using NEBNext Ultra RNA Library Prep Kit (NEB, Ipswich, MA, USA). Sequencing libraries were multiplexed and clustered on 1 lane of a flow-cell.

For shifts from LB_1.0x_ to LB_1.5x_, LB_2.0x_, and LB_2.5x_ at 180 min, samples were sequenced using a single-index, 2×150bp paired-end (PE) configuration (∼350M raw paired-end reads per lane) on an Illumina HiSeq 4000 instrument. Image analysis and base calling were conducted with HiSeq Control Software (HCS). Raw sequence data (.bcl files) were converted into fastq files and de-multiplexed using Illumina bcl2fastq v.2.20. One mismatch was allowed for index sequence identification.

For shifts from LB_1.0x_ to LB_0.5x_ at 60 and 125 min, samples were sequenced using a single-index, 2×150bp paired-end (PE) configuration (∼10M raw paired-end reads per lane) on an Illumina NovaSeq 6000 instrument. Image analysis and base calling were conducted with NovaSeq Control Software v1.7. Raw sequence data (.bcl files) were converted into fastq files and de-multiplexed using Illumina bcl2fastq v.2.20. One mismatch was allowed for index sequence identification.

#### c. Data analysis

Regarding the RNA-seq data analysis pipeline: i) RNA sequencing reads were trimmed to remove possible adapter sequences and nucleotides with poor quality with Trimmomatic (38) v.0.36 (for data from sequencing part 1) and v.0.39 (for data from sequencing part 2). ii) Trimmed reads were then mapped to the reference genome, *E. coli* MG1655 (NC_000913.3), using the Bowtie2 v.2.3.5.1 aligner, which outputs BAM files (39). iii) Then, *featureCounts* from the Rsubread R package (v.1.34.7) was used to calculate unique gene hit counts (40). Genes with less than 5 counts in more than 3 samples, and genes whose mean counts are less than 10 were removed from further analysis. iv) Unique gene hit counts were then used for the subsequent differential expression analysis. For this, we used the DESeq2 R package (v.1.24.0) (41) to compare gene expression between groups of samples and calculate p-values and log2 of fold changes (LFC) of RNA abundances using Wald tests (function *nbinomWaldTest*). P-values were adjusted for multiple hypotheses testing (Benjamini– Hochberg, BH procedure, (42)) and genes with adjusted p-values (False discovery rate (FDR)) less than 0.05 were selected to be further tested as being differentially expressed (Methods section *RNA-seq d*).

For logistical reasons, the sequencing platform for the RNA-seq data in Methods section *RNA-seq b* differ from one another. Consequently, the data sets used in Figure 3 and in Figure 5 cannot be compared quantitatively nor be used to infer gene-specific conclusions.

Finally, to analyse the data from LB_1.0x_ and LB_0.5x_ at 60 and 125 min and compare its results with the results from the data of Methods section *RNA-seq b Part 1* at 180 min, their raw count matrices were merged and only genes that passed the filtering were studied. The filtering removed genes with less than 5 counts in more than 6 samples, and genes whose mean counts were less than 10.

Moreover, we expect the overall sums of LFCs from each perturbation to equal zero since, in DEseq2, the median-of-ratios normalization calculates the normalizing size factors assuming a symmetric differential expression across conditions (i.e., same number of up- and down-regulated genes) (43). Further, it fits a zero-centered normal distribution to the observed distribution of maximum-likelihood estimates (MLEs) of LFCs over all genes (41). Both steps (perhaps related) force the mean LFC to be 0.

#### d. LFC criteria for differentially expressed genes

From past methods (44-46), we classified genes as statistically significantly differentially expressed (DE) due to the genome-wide perturbations, by setting a maximum FDR threshold for adjusted p-values (Methods section *RNA-seq c*) and also a minimum threshold for the absolute LFC of RNA numbers of individual genes (|LFC|).

From the RNA-seq data of each perturbation, from the *μ*_|*LFC*|_ of genes whose FDR > 0.05, named *μ*_|*LFC*|_(*FDR* > 0.05), we identified DEGs (DE Genes) as those that, in addition to having FDR < 0.05, also have |LFC| > *μ*_|*LFC*|_(*FDR* > 0.05). Specifically, we added the conditions: |LFC| > 0.4248 for LB_0.75x_, > 0.4085 for LB_0.5x_, > 0.4138 for LB_0.25x_, > 0.2488 for LB_1.5x_, > 0.2592 for LB_2.0x_, and > 0.2711 for LB_2.5x_, for accepting a gene as being significantly DE. For the data in LB_0.5x_ at 60 and 125 min we added: |LFC| > 0.2171 for LB_0.5x_ 60 min, and > 0.2977 for LB_0.5x_ 125 min. This allows removing from the data genes whose FDR < 0.05 but that, in fact, have a negligible LFC. Noteworthy, in no condition did we remove, from the set of DEG, more than 5 genes by applying this rule.

#### e. RNA-seq vs Flow-cytometry

RNA and protein abundances are expected to be positively correlated in bacteria, since transcription and translation are mechanically bound (47-49) and most regulation occurs during transcription initiation (50), which is the lengthiest sub-process (24).

To validate that this relationship holds during the genome-wide stresses, we randomly selected a set of genes whose LFC’s, as measured with RNA-seq, cover nearly the entire spectrum of LFCs observed genome-wide. Next, we measured their LFC in protein abundances, using the YFP strain library (25) (Methods section *Bacterial strains, media, and growth conditions and curves*) and flow-cytometry (Methods section *Flow-cytometry*), at 180 min after shifting the medium. The list of selected genes is shown in Supplementary Table S4. In detail, for the fold change levels of 1/8x, 1/4x, 1/2x, 1x, 2x, 4x, and 8x, we selected 3 genes whose LFC in RNA abundances is closest to that value (except for the 8x fold change, since only 2 genes were available). This range of values covers nearly the whole LFC spectrum observed by RNA-seq (Supplementary Figure S9).

### Transcription Factor Network of *Escherichia coli*

We assembled a directed graph of the network of TF interactions between the genes present in our RNA-seq data, based on the data in RegulonDB v10.5 (34), as of 28^th^ of January 2022. We used all reported TF-TF, TF-operon, and TF-TU interactions. These equally contribute to our network of gene-gene directed interactions. In detail, a TF or regulatory protein is a complex protein that activates/represses transcription of a transcription unit (TU) upon binding to specific DNA sites. A TU is one or more genes transcribed from a single promoter. Similarly, an operon are one or more genes and associated regulatory elements, transcribed as a single unit.

The TFN graph was analysed using MATLAB (2021b) and Network Analyzer v.3.7.2 plug-in in cytoscape (51) to extract the following network parameters specified from (51): number of nodes and directed edges, number of connected components, number of isolated nodes and self-loops, and single-gene in- and out-degree, edge-count, clustering coefficient, eccentricity, average minimum path length, betweenness and stress centrality, and neighbourhood connectivity. The statistics considered are shown in Supplementary Tables S5 and S19.

### Statistical tests

#### a. 2-sample T-test, 2-sample KS-test and one-sample Z-test

The 2-sample T-test evaluates the null hypothesis that the two samples come from independent random samples from normal distributions with equal means and unequal and unknown variances. For this, we have established a significance level of 10% significance level (P-value < 0.10) when applying the MATLAB function *ttest2*.

The 2-sample KS-test returns a test decision for the null hypothesis that the data from 2 data sets are from the same continuous distribution, using the two-sample Kolmogorov-Smirnov test. As above, we have set the null hypothesis at 10% significance level (P-value < 0.10).

The one-sample Z-test tests for the null hypothesis that the sample is from a normal distribution with mean *m* and a standard deviation σ. In this case, *m* and *σ* are estimated from the genes with K_TF_ = 0. As above, we have set the null hypothesis at 10% significance level (P-value < 0.10).

#### b. Fisher test

The Fisher test evaluates the null hypothesis that there is no association between the two variables of a contingency table. We reject the null hypothesis at 10% significance level (P-value < 0.1), meaning that the variables are significantly associated.

#### c. Correlations between data sets

The correlation between two data sets with known uncertainties (standard error of the mean (SEM) in each data point) was obtained by performing linear regression fitting using Ordinary Least Squares. The best fitting line along with its 68.2% confidence interval/bounds (CB) and statistics was obtained as described in Supplementary Materials and Methods 1.4 of (52). In short, the uncertainty of each of the N empirical data points was represented by m points, resulting in n = N×m points. Each of these points is obtained by random sampling from a normal distribution whose mean (µ) and standard deviation (σ) equal the mean and error of the empirical data point, respectively. It was set m = 1000, as it was sufficient to represent the error bars of the actual data points. We obtained the coefficient of determination (R^2^) and the root mean square error (RMSE) of the fitted regression line, and the p-values of the regression coefficients. The p-value of x (P-value_1_) was obtained of a T-test under the null hypothesis that the data is best fit by a degenerate model consisting of only a constant term. If P-value_1_ is smaller than 0.1, we reject the null hypothesis that the line is horizontal, i.e., that one variable does not linearly correlate with the other. When there are more than 3 data points, we also calculated regression coefficient of x^2^ (P-value_2_) of a T-test under the null hypothesis that the second order polynomial fit is no better than lower order polynomial fit, i.e., coefficient of x^2^ = 0. If P-value_2_ is smaller than 0.1, we reject the linear model favouring the quadratic.

To obtain the overall best non-linear fit (and its 68.2% CI) for the empirically measured datasets with uncertainties, Monte Carlo simulations (1000 iterations) were performed. In particular, to obtain Figure 2B, on each iteration, we randomly sampled each data point from a normal distribution whose mean and standard deviation are equal to the mean (actual value) and SEM of the corresponding empirical data point, respectively. Then a sigmoid (logistic) curve fitting (R P (2020). sigm_fit (https://www.mathworks.com/matlabcentral/fileexchange/42641-sigm_fit), MATLAB Central File Exchange. Retrieved August 6, 2020) was used to obtain the best fitting curve and its 68.2% CB for each iteration. Finally, the best fitting curve along with their 68.2% CB is obtained by averaging the respective values from the 1000 iterations.

**Figure 2.**
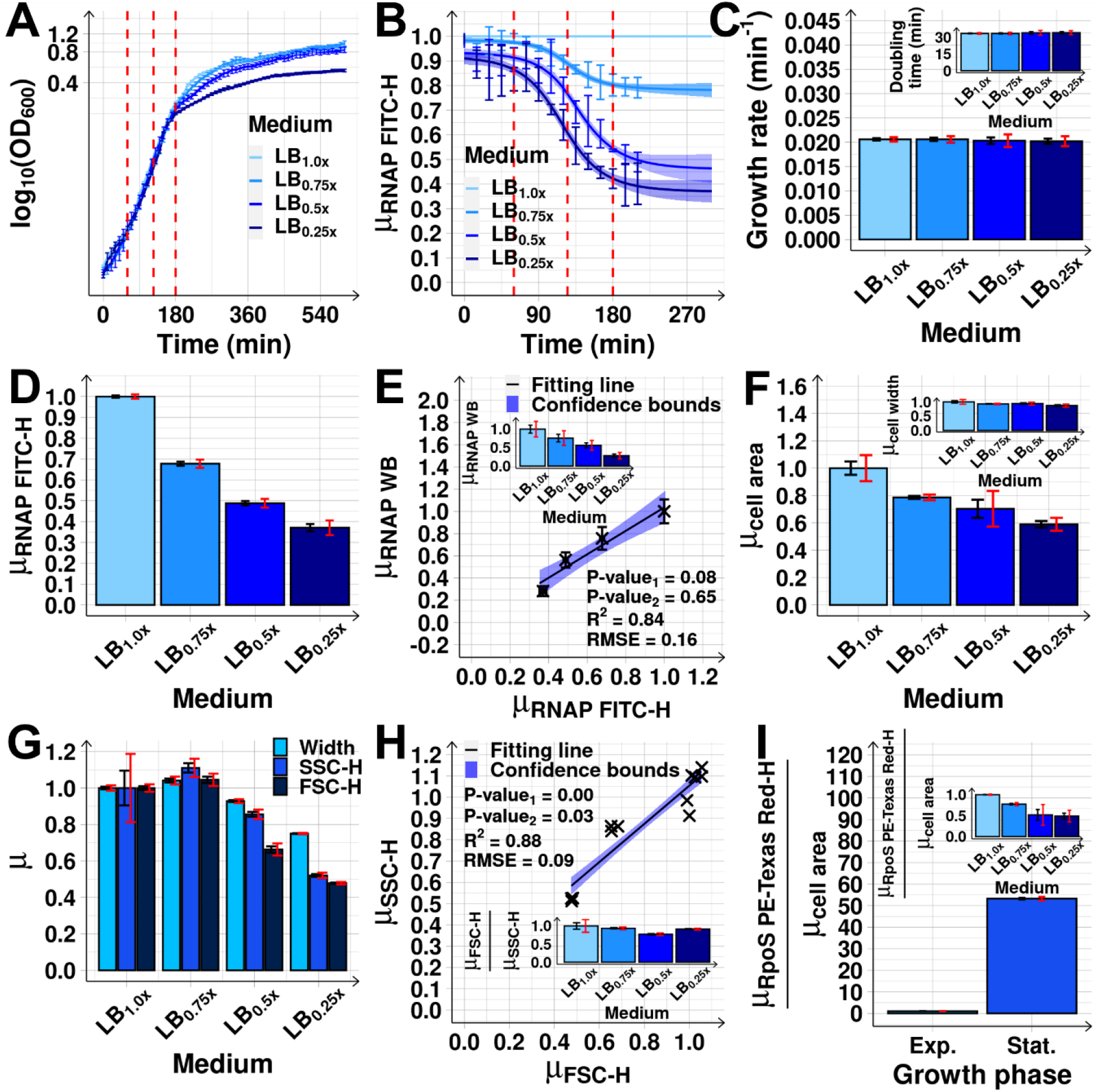
Cell growth and morphology, and RNAP concentration after medium dilutions. **(A)** Growth curves from OD_600_ measured every 10 min (Methods section *Bacterial strains, media, and growth conditions and curves*). The vertical dashed red lines mark when RNA-seq was performed. After ∼180 min, cells subject to different dilutions (LB_mx_) start differing in growth rates. **(B)** Mean single-cell RNAP-GFP fluorescence relative to the control (LB_1.0x_), *μ*_*RNAP FITC*−*H*_, measured every 15 min for 210 min by flow-cytometry (FITC-H channel). The mean cellular background fluorescence in each condition was subtracted (Methods section *Flow-cytometry*). The vertical dashed red lines mark when RNA-seq was performed. **(C)** Growth rates at 180 min after medium dilution. The inset shows the corresponding doubling times. **(D)** Mean single-cell RNAP levels (*μ*_*RNAP FITC*−*H*_) at 180 min relative to the control (Methods section *Flow-cytometry*). **(E)** *μ*_*RNAP FITC* −*H*_ plotted against *μ*_*RNAP WB*_ (RNAP levels measured by Western Blot, Methods section *Protein isolation and western blotting*). The inset shows *μ*_*RNAP WB*_ alone. **(F)** Mean cell area relative to the control, extracted from phase-contrast images (∼2000 cells per condition) (Methods section *Microscopy*). The inset shows the mean cell width relative to the control. **(G)** Mean (relative to the control) Width, FSC-H and SSC-H obtained by flow-cytometry (Methods section *Flow-cytometry*). **(H)** Mean (relative to the control) FSC-H versus SSC-H in each condition, obtained from 3 biological replicates. The inset shows the mean ratio between the relative FSC-H and SSC-H. **(I)** Mean mCherry-tagged RpoS (*μ*_*RpoS PE*−*Texas Red*−*H*_) concentration in the stationary growth phase relative to the exponential growth (set to 1), as measured by mean single-cell fluorescence (PE-Texas Red channel, Methods section *Flow-cytometry*) over mean cell area (*μ*_*cell area*_) (Methods section *Microscopy*), after subtracting mean background fluorescence(s). The inset shows the same, but after each medium dilution. Measurements in (D)-(I) taken 180 min after medium dilution. Data points are from 3 biological replicates (except for (A) and (B), where 6 replicates were used). *μ* stands for mean relative to the control. In (A)-(C) error bars represent the SEM. In (B) and (D)-(I), black error bars are the SEM and red error bars are the 95% confidence bounds (CB) of the SEM. In (C), (E) and (H), the best fitting lines and their 68% CB and statistics (R^2^ and RMSE), and P-values at 10% significance level) were obtained as described in Methods section *Statistical tests c*.

Finally, to create null-models of how variable X affects variable Y, we performed random sampling without replacement of both X and Y datapoints. The number of samplings and the sampling size (number of samples in each sampling) are set to the maximum array size possible to us (∼45980×45980, 15.8 GB). The sampling size is set to 5% of the number of datapoints (size_XY) and the number of samplings (K) is set according to Max_size/(0.05×size_XY) where Max_size = 45980/2. Next, for both X and Y, we combine the sampled datapoints in a vector (sample_X, sample_Y) and calculate the correlation between sample_X and sample_Y by linear regression fitting using Ordinary Least Squares. To correct for over-representation of the original datapoints, we corrected the degrees of freedom to be (size_XY – C), where C is the number of parameters. In detail, for the linear regression fitting, C equals to 2 (intercept and slope of best fitting line).

#### d. ANCOVA test to evaluate if two lines can be distinguished

To evaluate if two lines are statistically different, we performed the analysis of covariance (ANCOVA) test (53). ANCOVA is an extension of the one-way ANOVA to incorporate a covariate. This allows comparing if two lines are statistically distinct in either slope or intercept, by evaluating the significance of the T-test under the null hypothesis that both the slopes and intercepts are equal.

### Figures

Figures were produced in R (v.3.6.0) using the packages ‘ggplot2’ (v.3.2.0), ‘pheatmap’ (v.1.0.12), ‘VennDiagram’ (v.1.6.20) along with ‘grid’ (v.3.6.0), ‘gridExtra’ (v.2.3), ‘gplots’ (v.3.0.1.1), ‘R.matlab’ (v.3.6.2), ‘dplyr’ (v.1.0.2), ‘scales’ (v.1.0.0), ‘Metrics’ (v.0.1.4) and ‘fitdistrplus’ (v.1.0-14).

## RESULTS

### Effects of medium dilution on cell growth, morphology, and RNAP concentration

We first studied how the RNAP concentration changes with medium dilutions. From a control medium (‘LB_1.0×_’), we moved cells to diluted media (LB_0.75×_, LB_0.5×_, or LB_0.25×_, Methods section *Bacterial strains, media, and growth conditions and curves*). RNAP levels start changing ∼75 min later, based on an as yet to be identified mechanism, stabilizing at ∼165 min (Figure 2B). Given this timing of events, measurements to assess the effects on the RNA population should be performed after ∼165 min.

We also considered that at ∼180 min (Figure 2A) the cells are at late mid-log phase. Thus, measuring the effects of changing RNAP should occur *prior* to ∼180 min, since leaving the mid-log phase will involve significant, unrelated genome-wide changes in RNA abundances (54-57). From the point of view of cell divisions, from the moment when the RNAP starts changing up to the moment when we measure the short- and the mid-term changes in RNA abundances, on average, less than one cell cycle and less than two cell cycles should have passed, respectively.

Interestingly, this time moment (∼180 min) matches our predictions of when, on average, RNA abundances have changed due to changes in the abundances of both RNAP as well as direct input TF. In detail, from the timing of the changes in RNAP (Figure 2B) and from known rates of RNA and protein production and degradation in *E. coli* (25,28-31), we expect widespread heterogenous short-term changes in RNA abundances to occur, on average, at ∼120-135 min after shifting the medium (at which moment the RNAP has already changed significantly). Changes in the corresponding protein abundances should then occur tenths of minutes later, i.e., at ∼ 160-175 min (29-31).

Soon after, we expect additional changes in RNA abundances, now due to changes in direct input TF abundances. This second stage of events, here classified as ‘mid-term’, should occur between ∼165-180 min. This is also when cells are in the late mid-log phase (Figure 2A), while cell growth rates do not yet differ between conditions (Figure 2C) and cell sizes only differ slightly (Figures 2F-2H, Supplementary Figures S3 and S4, Methods sections *Microscopy* and *Flow-cytometry*). Such is relevant, since growth rates affect protein concentrations due to dilution in growth and division (58,59). Finally, at 180 min, the σ^38^ concentration is lower than at 0 min (Figure 2I inset and Supplementary Figure S5), in agreement with previous reports (27,35,60,61), suggesting that the cells are not committed to the stationary growth phase. The same is observed for the Crl protein (Supplementary Figure S6), which is a protein that contributes to the expression of genes whose promoter is recognized by σ^38^ and that is known to be at higher abundance during stationary phase (reported in (62) and confirmed here (Supplementary Figure S6)).

Given the above, to capture the average mid-term effects of RNAP shifts, we measured the transcriptome at 180 min (Figure 2A). This timing should allow discerning the average genes’ behaviour, under the influence of their local network of TF interactions, albeit the diversity in RNA and protein production and decay kinetics, etc. RNAP levels at that moment are shown in Figures 2D and 2E, Supplementary Figure S2 and Supplementary Table S2. Similar RNAP downshifts have been observed in natural conditions (63) and described in (23,26,27).

### Genome-wide mid-term responses correlate with shifts in RNAP concentration

Transcription rates are expected to follow the free RNAP concentration in a cell, rather than the total RNAP concentration (which is the sum of the free RNAP with the RNAP engaged with the DNA). We here measured the total RNAP concentration. However, within the range of conditions studied, the fractions of free and DNA-bound RNAP remain rather constant (26). Therefore, the total RNAP is a good proxy for the free RNAP. Specifically, using modified strains and plasmids controlled by lac and tet mutant promoters (64-66), whose regulatory mechanisms have been dissected, it was shown that their transcription rates are linearly correlated with the total RNAP concentration (26). From here on, when mentioning RNAP concentration, we refer to the total RNAP concentration.

The increasing medium dilution and corresponding decreases in RNAP concentration (Figure 3A) cause RNA-seq profiles at 180 min with increasingly broad distributions of single-gene LFCs (Supplementary Figures S7 and S8A-S8C and Supplementary Table S3). Specifically, the mean absolute LFC of the 4045 genes (*μ*_|*LFC*|_) and the number of DEGs increased with medium dilution (Figures 3B and 3C).

**Figure 3.**
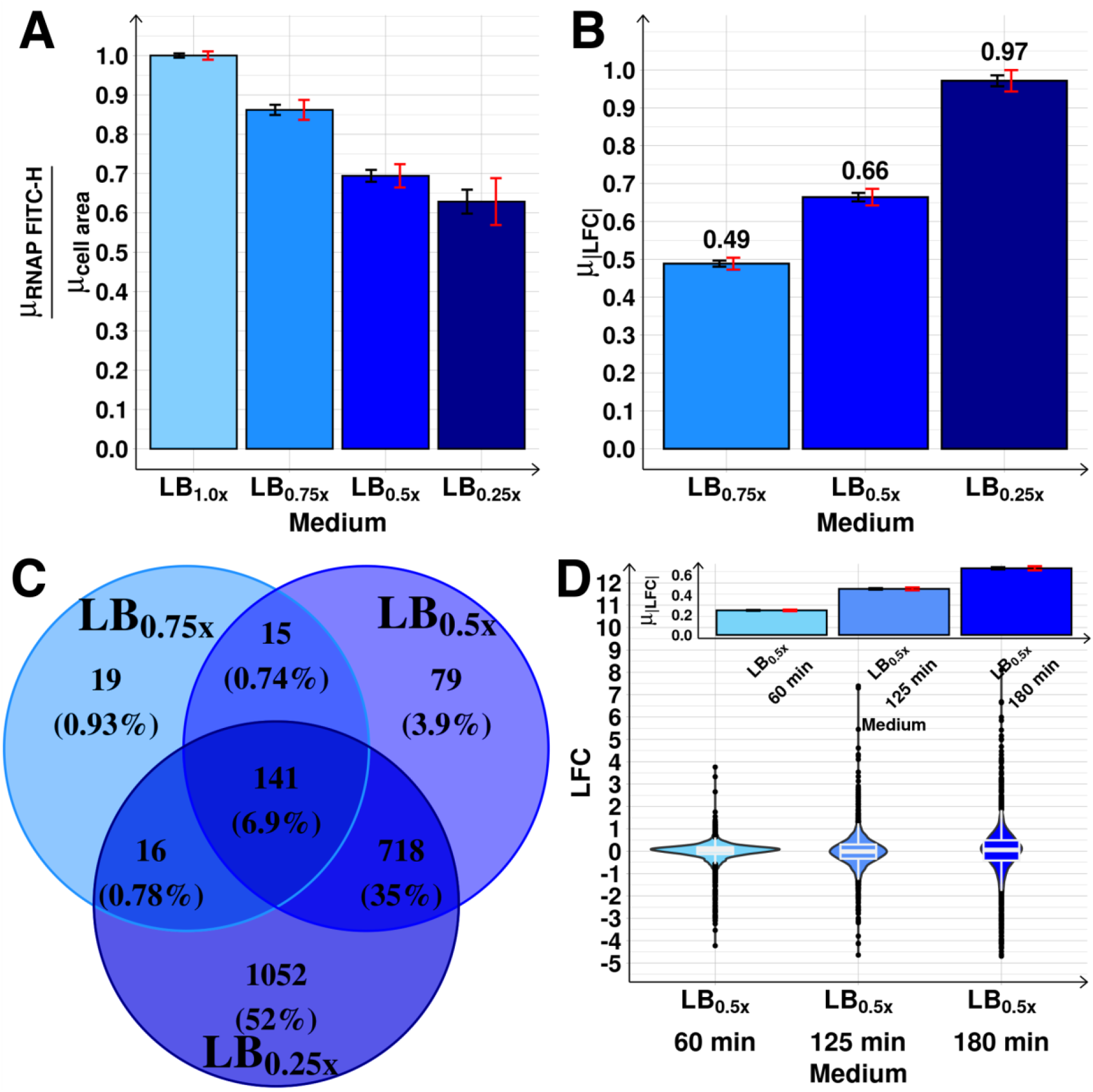
Genome-wide effects on the transcriptome of diluting the medium. **(A)** Ratio between the RNAP measured by FITC-H (*μ*_*RNAP FITC*−*H*_) at 180 min (Methods section *Flow-cytometry*), and the mean cell area (*μ*_*cell area*_) obtained by phase-contrast microscopy (Methods section *Microscopy*). Values relative to the control (LB_1.0x_). **(B)** *μ*_|*LFC*|_ in each medium. **(C)** Venn diagram of the number (and percentage relative to the total number of genes) of DEG. In (A) and (C), black error bars are the SEM, while red error bars are the 95% CB of the SEM. **(D)** Violin plot with the maximum, minimum, median, interquartile ranges, and probability density of the distributions prior to RNAP changes (LB_0.5x_ at 60 min) and the subsequent short-(LB_0.5x_ at 125 min) and mid-term (LB_0.5x_ at 180 min) responses to shifting RNAP. The inset shows *μ*_|*LFC*|_ of the distributions.

These RNA changes correlate with subsequent changes in protein levels (Supplementary Figure S9, Methods sections *Flow-cytometry* and *RNA-seq*). This suggests that no significant translational or post-translational regulation is taking place in between the perturbation and the measurements, that would alter protein abundances significantly.

Interestingly, while both *μ*_|*LFC*|_ and DEGs numbers follow the RNAP concentration (Supplementary Figures S8D and S10B), these relationships are not strictly linear (p-value of 0.29, Supplementary Figure S8D), supporting the notion that, in addition to RNAP, the direct input TFs are also influential (note that the assumption of linearity in the absence of the influence of input TFs, observed and discussed in (26), is only expected to occur within a narrow range of parameter values).

Notably, some of the genes may be also influenced by sources other than RNAP and direct input TFs, such as supercoiling buildup. Also, some input TFs other than the direct input TFs maybe be influential. However, we show evidence below that this does not affect the average results (Figure 4C and Supplementary Figure S18).

**Figure 4.**
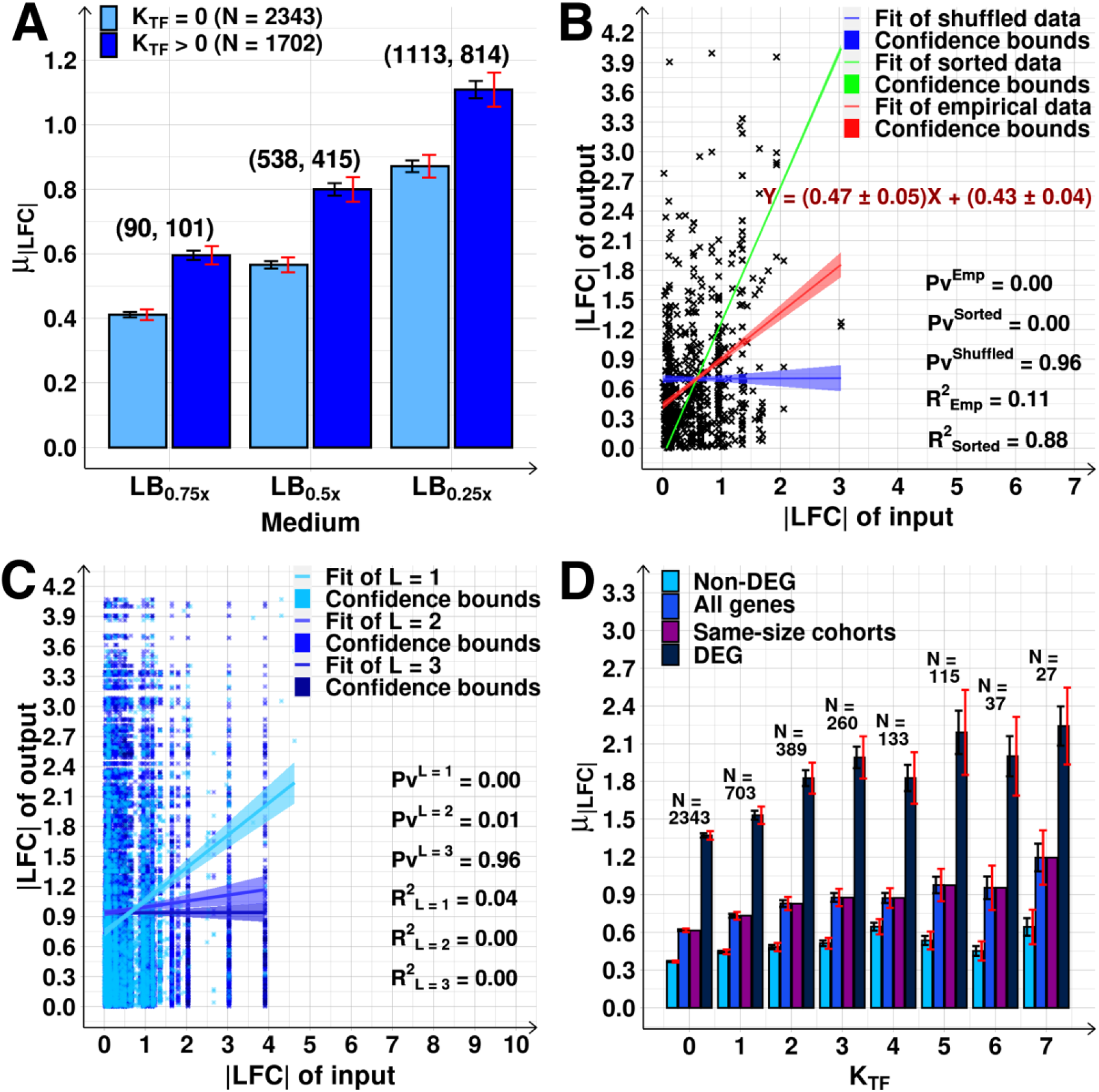
Genome-wide propagation of the effects of shifting RNAP in the TFN. **(A)** *μ*_|*LFC*|_ of *N* genes with and without input TFs (K_TF_ > 0 and = 0, respectively). On the top of each bar is the number of DEGs in each set. **(B)** |LFC| of genes with K_TF_ = 1 versus the |LFC| of genes coding their direct input TFs. Data from the LB_0.5x_ shift. The red line is the best fit. The blue line is the null-model fitting lines and was obtained as described in Methods section *Statistical tests c*. The green line is the best fit after sorting the input-output pair values to maximize the correlation. Shadows are their 68% CB. The equations of the red fitting lines with ‘±’ inform on the standard error of the slope. **(C)** Scatter plots between |LFC| of output and input genes distanced by a minimum path length L of 1, 2, and 3 input TFs (edges) in the TFN, respectively (data from LB_0.5x_). Only for L = 1 do the activities of output and input genes correlate (P-value_1_ > 0 and R^2^ > 0). **(D)** *μ*_|*LFC*|_ of all genes, DEG, non-DEG, and cohorts of randomly selected genes of the same size (‘same sized cohorts’) for K_TF_ = 0 to 7, using merged data from all shifts (LB_0.75x_, LB_0.5x_ and LB_0.25x_). Black error bars are the SEM and red error bars are the 95% CB of the SEM. Best fitting lines and 68% CB obtained using *FITLM* (MATLAB). p-values, obtained using the null hypothesis that the data is best fit by a horizontal line, are not rejected at 10% significance level. (B) and (C) do not include a few data points to facilitate visualization. See Supplementary Figures S14 and S18 for complete data.

We also performed RNA-seq *prior* to when most signals, generated by the shift in RNAP, propagated in the TFN. First, we measured LFCs at 60 mins after diluting the medium (Figure 1). From Figure 2B, at this moment, RNAP abundances have not yet changed relative to the control. In agreement, the genome-wide *μ*_|*LFC*|_ is very weak (Figure 3D). We further performed RNA-seq at 125 min. At this moment, RNAP levels have already reduced significantly (Figure 2B), but we do not expect input TF abundances to have changed significantly given protein production times (Figure 1). In agreement, |LFC|s at 125 min are stronger than at 60 min, but much weaker than at 180 min (Figure 3D). We conclude that the mid-term changes in the TFN have not occurred yet (further evidence is provided below). Given this, from here onwards, we focus on the state of the TFN at 180 min.

### Influences from regulators other than RNAP

We investigated whether other factors influenced the global response of the TFN. We considered global regulators (GR), σ factors, (p)ppGpp, and non-coding sRNAs. We assumed the classification of GR in (67,68) as an input TF that regulates a large number of genes that rarely regulate themselves and participate in metabolic pathways. Meanwhile, we did not account for promoters’ close proximity (e.g., tandem formation), since a recent study (69) showed that, under similar stress, while close proximity causes transcription interference (reducing overall transcription levels), it does not influence if a gene is up- or down-regulated by input TFs.

First, from the RNA-seq, given the large numbers of DEGs (more than 1000 for the two strongest dilutions (Supplementary Figure S10A)), and the linear correlation between these numbers and *μ*_|*LFC*|_ (Supplementary Figure S10C), we argue that the responsive genes are not constrained to a specific cluster, such as genes responding to a global regulator (GR) other than RNAP (the most influential is, arguably, σ^70^ with 1555 genes recognizing it, while other GRs control less than 510 genes each (34)).

Also, from the RNA-seq, we analysed the relative abundances of GRs, σ factors and of their output genes. From Supplementary Figures S26A and S26C, apart from rpoS (an input TF recognized by 321 genes) and flhC (an input TF recognized by 75 genes), GRs and σ factors did not change significantly (Supplementary Figure S26). Further, those two changes (rpoS and flhC) were positively correlated with the RNAP concentration (Figure 2I inset and Supplementary Figure S5), not allowing to separate their effects. Also noteworthy, alternative σ factors did not change significantly relative to σ^70^ (Supplementary Figure S26E), which would have changed the competition for RNAP binding.

Given this, we failed to find evidence that the σ factors and GRs were influential, globally, in the mid-term responses. Supplementary Table S15 lists the conclusion for each specific GR and σ factor and Supplementary Figure S27 shows these results at 125 min.

We then investigated if (p)ppGpp could be influential since, under some nutrient starvation conditions, they affect ∼1000 genes by binding RNAP and altering its affinity for their promoters (3). Reports suggest that the effects are rapid (5 to 10 min (3)). In agreement, genes responsive to (p)ppGpp (3) exhibited abnormal short-term responses (Supplementary Table S20). However, their mid-term responses at 180 min were no longer atypical and, instead, followed the RNAP changes. The expression of spoT, one of the genes responsible for ppGpp synthesis, also followed the RNAP (Supplementary Figure S28). As such, we could not establish a long-lasting global influence from (p)ppGpp in response to growth-medium dilution. Nevertheless, the LFCs of the 14 out of the 22 genes coding for rRNAs listed in RegulonDB did reveal atypical behaviors (Supplementary Table S22).

Next, we searched for unique behaviours in sRNAs by analysing the LFC of the 93 sRNAs reported in RegulonDB. Their behaviour was not atypical, neither at 180 min after the perturbations (Supplementary Table S21), nor at 125 min. Further, we analysed if their output genes followed their behaviour. We found that the LFCs of genes directly regulated by the sRNAs were not correlated with their input TFs, neither at 125 min, nor at 180 min after the medium shifts. Specifically, of the 93 sRNAs, 37 of them have known output genes (in a total of 145 outputs). The RNA-seq data provided information on the LFC of 40 of the 145 outputs. When searching for linear correlations between the pairs of LFCs of sRNAs and their output genes, respectively, in the short-term (125 min in LB_0.5x_) and in the mid-term (180 min in LB_0.5x_), we found an R^2^ of 0.03 (p value = 0.18) at 125 min and an R^2^ of 0.05 (p value = 0.10) at 180 min, respectively. We thus cannot conclude that sRNAs were influential during the short- and mid-term responses to the stresses.

### Input TFs influence the transcriptional response

If the TFN influences the genes’ mid-term response to the shift in RNAP concentration, this should cause genes with and genes without input TFs to behave differently, since the latter should only be affected by the RNAP abundances.

In agreement, genes with input TFs had higher *μ*_|*LFC*|_ than genes without input TFs (Figure 4A, Supplementary Table S6 and Supplementary Figure S13). Also, the |LFC| of output genes and of genes coding for their direct input TFs correlate statistically (Figure 4B, Supplementary Figure S14 and Supplementary Table S7). Therefore, on average, TF-gene interactions affected the single-gene, mid-term responses as hypothesized (Figure 1).

### Input TFs influence all genes within operons

When considering the TFN topology, we have accounted for TF-gene interactions both between the input TF and the first gene of an operon or transcription unit (TU), but we also accounted for the interactions between the same input TF and the other genes of the operon or TU (illustration of TUs and operons in Supplementary Figure S11B, which follows the standard definition of a group of two or more genes transcribed as a polycistronic unit (1)).

If we had not account for all these interactions, we would have failed to correlate the activities of genes interacting with each other. For example, consider an operon consisting of genes X_1_ and X_2_ and assume that gene A represses X_1_ and X_2_, by repressing their common promoter. If X_1_ is an input TF to gene C, while X_2_ is an input TF to gene D, then gene A should indirectly affect both genes C and D. If we had ignored the interaction between A and X_2_, because it is not the first gene in its operon, we would be able to explain why A affects C, but we would fail to explain why A affects D.

Further, many operons contain sets of genes whose RNAs code for subunits of the same protein complex (70,71). However, the opposite is also true and, the fraction of complexes encoded by proteins from different TU’s is higher than those encoded from the same operon (72). This supports the need to track interactions between input TFs and genes in any position in an operon or TU.

To test if the positioning of the genes in the operon influenced their responsiveness to their input TFs, as a case study, we considered operons with 3 genes (which account for ∼21% of all operons with more than 1 gene (34)). We found that the positioning of the genes did not affect significantly how they relate to the input TFs (Supplementary Figure S16). We obtained similar results for TUs (Supplementary Figure S17). The tests of statistical significance are shown in Supplementary Tables S9-S12.

### Genes expressing TFs are correlated with their nearest neighbour output genes

In general, we expect that, after a genome-wide perturbation, signals will propagate between nearest neighbour genes. Depending on the measurement time and given the diversity in the kinetics of different genes and RNA and protein lifetimes, this should result in some signals propagating between genes considerably distanced in the TFN, with the number of such signals decreasing rapidly with the path length between the pairs of genes considered.

Given the interval between the shift in RNAP levels and the sampling for RNA-seq (Figure 1), we hypothesized that, on average, at 180 min (i.e., ∼70 min after the RNAP changed relative to the control), mostly only genes directly linked by input TFs should exhibit correlated responses. Results in Figure 4C support this. Genes distanced by 1 input TF (L = 1, i.e., directly linked) have related |LFC|s, while genes distanced by 2 input TFs in the TFN have much less correlated responses (albeit still statistically significant). Finally, we found no correlations between the |LFC|s, of genes distanced by 3 input TFs (Supplementary Figure S18).

Noteworthy, the lack of correlation between genes separated by L > 1 could also be partially due to interference from the TFs of the ‘intermediary’ genes between the gene pairs. However, this is only a possibility when all input TFs involved can change in abundance in less than 60 min, which is likely uncommon in *E. coli*. This is supported by the RNA-seq data at 125 min after medium dilution (Supplementary Figure S19), where even direct input TFs and output genes are weakly correlated, suggesting lack of time for most signals to have propagated between nearest neighbours (Supplementary Figure S19).

### The number of input TFs of a gene correlates to the magnitude of its transcriptional response

We investigated if the genes mid-term responses are sensitive to their number of input TFs, K_TF_ (Supplementary Figure S20A). When averaging the results from the three perturbations (Figure 4D), we found that the average of the absolute LFCs, *μ*_|*LFC*|_, increases with K_TF_, whether considering all genes or just the DEGs (Figure 4D, Supplementary Figure S21 and Supplementary Tables S13). This holds true even for non-DEGs (Figure 4D), which justifies also considering these genes when studying the genome-wide effects. In agreement, we found no trend in the fraction of DEGs when plotted against K_TF_ (Supplementary Figure S23).

For comparison, neither at 60 min nor 125 min do the genes’ response and their number of input TFs correlate (Supplementary Figures S14 and S15 and Supplementary Tables S7 and S8).

We verified that the relationship between *μ*_|*LFC* |_and K_TF_ at 180 min is not an artifact caused by a decrease in cohort size with K_TF_. We used bootstrapping to obtain cohorts of randomly sampled genes with increasing K_TF_ (10000 cohorts). We imposed a cohort size equal to the number of genes with K_TF_ = 7 (27 genes). The new, estimated *μ*_|*LFC*|_ was always within the SEM of the *μ*_|*LFC*|_ of the cohorts of all genes (Figure 4D). Finally, we again verified that considering only the first gene of each operon does not affect how *μ*_|*LFC*|_ and K_TF_ relate (Supplementary Figure S25).

### The correlation between input and output genes responses decreases with the number of input TFs

Most input TFs discernibly affect the output genes (Supplementary Figure S14), except when K_TF_ > 5 (perhaps due to saturation).

Nevertheless, the correlation between inputs and outputs appears to be decreasing with K_TF_, as the average slopes of the fitted lines between |LFC| of the output and |LFC| of each input (Supplementary Figure S14) decreased with the K_TF_ of the output gene (Supplementary Figure S20B), as did the R^2^ between input-output pairs (Supplementary Table S7).

This could explain why, when plotting |LFC| against the RNAP concentration, there is a weak trend towards increased slope with K_TF_ (Supplementary Figures S20C and S21)

### The variability in single-gene |LFC| increases with K_TF_

We also investigated if the variability in |LFC|s, as quantified by its standard deviation *σ* _|*LFC*|_, relates with K_TF._ There should exist (at least) four sources of this variability: a) RNA-seq measurement noise (73,74); b) intrinsic and c) extrinsic noise in gene expression (75,76), and d) TF and non-TF dependent regulatory mechanisms.

Examples of the variability are shown in Supplementary Figures S24C (genes with null K_TF_), S24F (genes with two global regulators, FNR and ArcA) and S24D and S24E (genes controlled by the global regulators FIS or CRP) (see also Supplementary Table S14). Overall, from a genome-wide perspective, *σ*_|*LFC*|_ increases with K_TF_ (Supplementary Figure S24A) in a similar manner as does *μ*_|*LFC*|_, and the two values are also related (Supplementary Figure S24B).

### Other topological features of the TFN do not influence mid-term responses

Globally, the TFN of *E. coli* has in- and out-degree distributions that are well fit by power laws (Supplementary Figures S12E_1_, S12E_2_, S12F_1_ and S12F_2_) (77,78), which may explain its relatively short mean path length (Supplementary Figure S12G and Supplementary Table S5).

Having established a relationship between the response kinetics and the indegree of the TFN, we next searched for correlations between |LFC| and other single-gene topological traits (Methods section *Transcription Factor Network of Escherichia coli*), namely, the average shortest path length, betweenness, closeness and stress centrality, clustering coefficient, eccentricity, out-degree, neighbourhood connectivity, and edge-count (51). Of these, only the clustering coefficient was statistically correlated with the |LFC| (p-value < 0.1) (Supplementary Table S20). However, it should not be influential, since the corresponding R^2^ is nearly zero (R^2^ = 0.01).

### The numbers of activating and repressing input TFs differ in most genes

In our original hypothesis, the mid-term response (|LFC|) of a gene should follow from the bias in the numbers of activators and repressors in its set of input TFs (Figure 1B_4_ and Supplementary Figure S11A). In detail, we predicted that if the sum of regulatory effects (*r*) of the input TFs (i.e., bias *b* = |∑*r*|) is null (unbiased), then the gene should have weak or zero mid-term LFC. Also, the |LFC| should increase with *b*.

We tested this hypothesis by extracting information on the input TFs and corresponding *r* values for each gene from RegulonDB. We set *r* of an input TF to +1 if it is activating, to -1 if it is repressing, and to 0 if it is either dual or unknown (Supplementary Figure S12B), and we obtained the absolute sum of the regulatory effects of the input TFs for each gene: |*b*|.

From the data in RegulonDB, while the gene-TF interactions that are repressions and activations exist in similar numbers, the numbers of repressor TFs exist in larger numbers (Supplementary Figures S12A-S12C).

Also, of the genes with input TFs, most (∼85%) have a non-zero |*b*| (Supplementary Figure S12D and Supplementary Table S16). This can explain why so many are mid-term responsive (Figure 3C), even though the genome-wide numbers of *activation* and *repression interactions* are similar (Supplementary Figure S12B). This may also explain why genes with K_TF_ ≥ 1 have higher |LFC| than genes with K_TF_ = 0 (Figure 4A).

### The bias in the input TFs follows the number of input TFs

Using information from RegulonDB, we found that the mean bias, *μ*_|*b*|_, increases with K_TF_ (Figure 5A, light blue), except for K_TF_ > 5, which includes only ∼64 out of 4045 genes (Supplementary Table S17). The same is observed if considering only the first gene of each operon (Supplementary Figure S29).

To test if these results were affected by local topological specificities, we employed an ensemble approach (Supplementary Results section *Estimation of the expected* 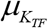 *and *μ*_|*b*|_ using an ensemble approach*) which reduces their influence in the estimations (79). We sampled genes (with replacement) to form cohorts with a given average K_TF_ (from 1 to 5, due to insufficient samples for higher K_TF_). Since this caused the relationship between *μ*_|*b*|_ and K_TF_ to be more stable (Figure 5A), from here onwards, we use the ensemble approach to study the influence of the logical and topological features on the response’s dynamics to the RNAP shifts.

**Figure 5.**
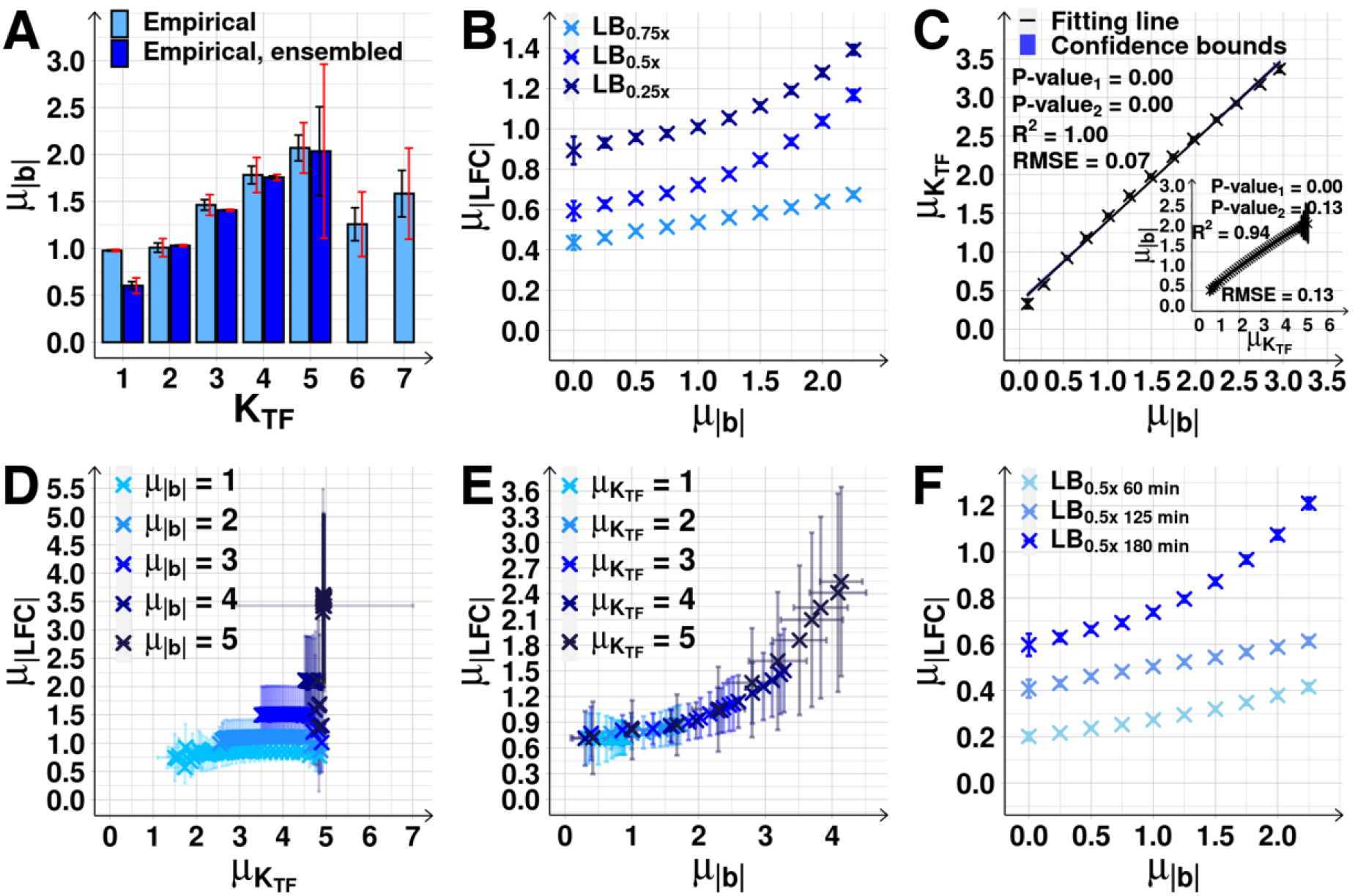
Effect of biases *μ*_|*b*|_ on the magnitude of the response of output genes. **(A)** *μ*_|*b*|_ as a function of K_TF_ (light blue) of gene cohorts with all genes (light blue) and of gene cohorts assembled using the ensemble approach (dark blue). Supplementary Table S17 shows the fractions of genes with equal *b* and K_TF_. Black error bars are the SEM, and red error bars are the 95% CB of the SEM. Dark blue bars not shown for K_TF_ > 5 due to small sample sizes. **(B)** Mid-term *μ*_|*LFC*|_ as a function of *μ*_|*b*|_, obtained using the ensemble approach (Supplementary Results section *Estimation of the expected* 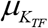 *and μ*_|*b*|_ *using an ensemble approach*, Supplementary Figures S30 and S31 and Supplementary Table S18). **(C)** *μ*_|*b*|_ plotted against the corresponding 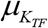, mean of K_TF_ of the cohorts in (B). The inset shows the inverse correlation plot for the cohorts in Supplementary Figure S30, assembled based on 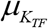 (Supplementary Results section *Estimation of the expected* 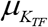 *and μ*_|*b*|_ *using an ensemble approach*). Shown are best fitting lines and 68% CB (shadow areas, barely visible), R^2^, RMSE, and P-value (Methods section *Statistical tests c*). **(D)** *μ*_|*LFC*|_ of gene cohorts with increasing 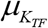, but constant *μ*_|*b*|_ (from 1 to 5) (Supplementary Results section *Estimation of the expected* 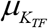 *and μ*_|*b*|_ *using an ensemble approach*). **(E)** *μ*_|*LFC*|_ of gene cohorts with increasing *μ*_|*b*|_, but constant 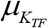 (from 1 to 5) (Supplementary Results section *Estimation of the expected* 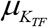 *and μ*_|*b*|_ *using an ensemble approach*). **(F)** *μ*_|*LFC*|_ as a function of *μ*_|*b*|_ prior to RNAP changes (60 min) as well as the short-term (125 min) and the mid-term responses (180 min) to RNAP changes when shifting to LB_0.5x_. In (D) and (E) the data is merged from the 3 conditions corresponding to (B). In all figures the error bars are the SEM. Since the 3 conditions differ slightly in mean values (Figure 5B), the SEM is larger than when observing each condition separately.

### The bias of the sets of input TFs can explain the mid-term responses of individual genes

From the data in RegulonDB and using the ensemble approach (Supplementary Results section *Estimation of the expected* 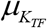 *and μ*_|*b*|_ *using an ensemble approach*), we formed random cohorts of genes with an imposed average |*b*|. Next, from the mid-term RNA-seq data, we calculated the average *μ*_|*LFC*|_ of the set of cohorts with a given *μ*_|*b*|_. We found that *μ*_|*LFC*|_ increases with *μ*_|*b*|_ Figure 5B.

Interestingly, *μ*_|*b*|_ and 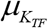 are strongly correlated in the TFN of E. *coli* (Figure 5C). To assert which one controls *μ*_|*LFC*|_, we assembled cohorts differing in 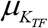, but not in *μ*_|*b*|_. In these, *μ*_|*LFC*|_ does not increase with 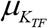 (Figure 5D). We also assembled cohorts differing in *μ*_|*b*|_, but not in 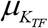. In these, *μ*_|*LFC*|_ increases with *μ*_|*b*|_ (Figure 5E). Thus, the increase of *μ*_|*b*|_ with 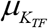 (Figure 5C), is what explains the increase in *μ*_|*LFC*|_ with K_TF_ (Figure 4D).

Finally, for comparison, we also investigated the relationship between *μ*_|*b*|_ and 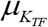 *prior* to the perturbation and in the short-term (at 60 min and at 125 mins after shifting the medium, respectively Figure 1A). From Figure 5F, first, the *μ*_|*LFC*|_ at 125 min is stronger than at 60 min. This agrees with the expectation that shifts in RNAP suffice to shift the |LFC| of many genes. Second, the *μ*_|*LFC*|_ at 180 min is stronger than at 125 min. This agrees with our expectation that, at 125 min, input TFs numbers have not yet changed significantly in order to enhance the |LFC| of their output genes (Figure 1).

### RNA numbers follow the RNAP concentration, not the medium composition

We next increased growth medium richness, instead of diluting it (Methods section *Bacterial strains, media, and growth conditions and curves*). As before, we limited this so as to not alter growth rates significantly in the first 180 min (Figures 6A and 6B), while altering RNAP levels (Figure 6C).

**Figure 6.**
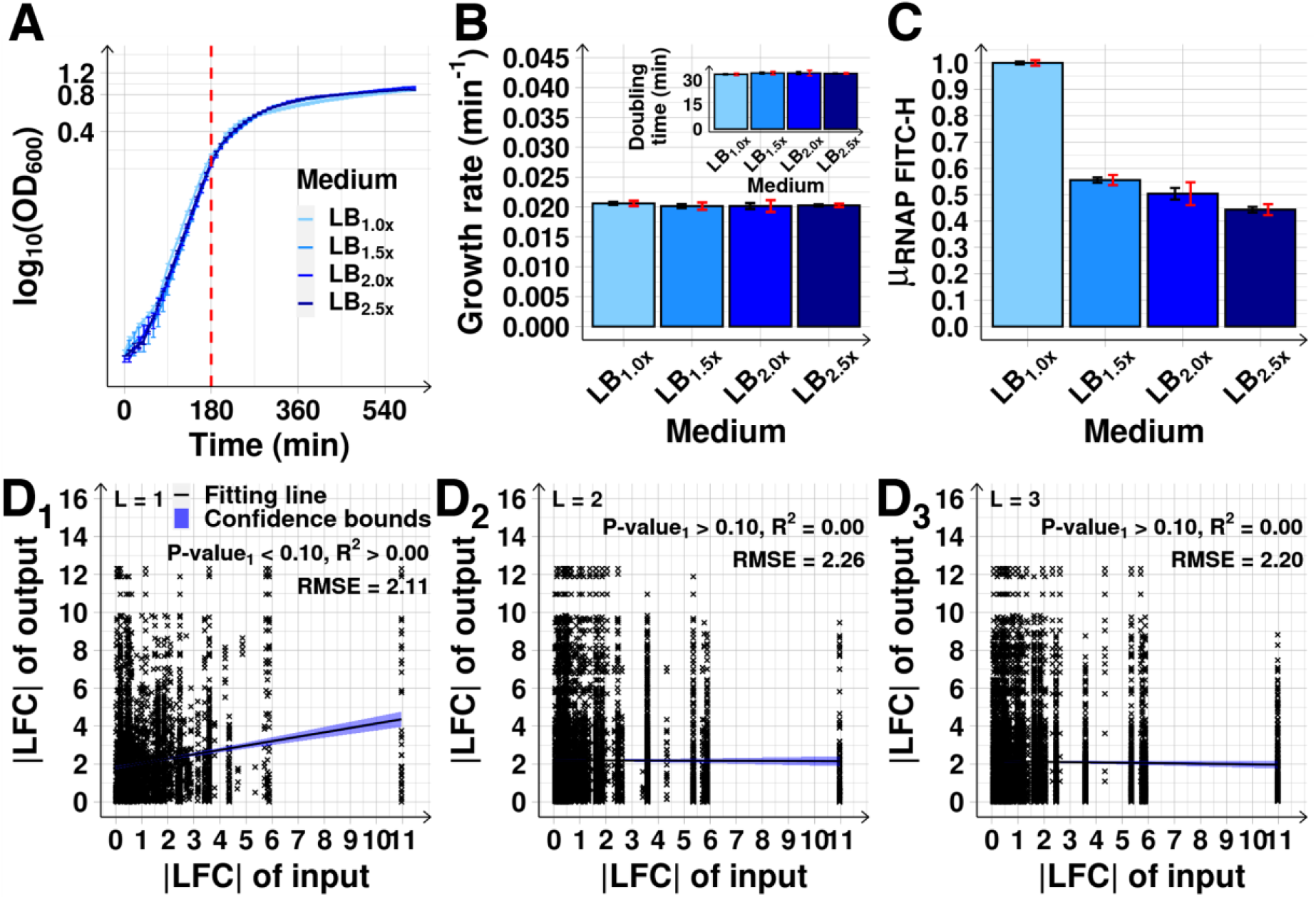
RNAP levels following increasing medium richness and corresponding relationships between |LFC|s of pairs of genes separated by specific path lengths, L. **(A)** Growth curves from OD_600_ assessed every 10 min (Methods section *Bacterial strains, media, and growth conditions and curves*), following each medium shift. **(B)** Growth rates at 180 min after medium enrichment. The inset shows the corresponding doubling times. **(C)** Mean RNAP levels relative to the control estimated from single-cell RNAP-GFP fluorescence intensities (FITC-H) (*μ*_*RNAP FITC*−*H*_). **(D**_**1**_**-D**_**3**_) Scatter plots between absolute LFC (|LFC|) of outputs and corresponding input genes distanced by L (path length) of 1, 2, and 3 transcription factors, respectively. Data from the LB_2.5x_ condition. Shown are the best fitting line and its 68% CB (blue shadow), and the R^2^ and RMSE of the fitted regression line, along with its p-value at 10% significance level under the null hypothesis that this line is horizontal. From (A) to (C) the black error bars are the SEM and red error bars represent the 95% CB of the SEM.

As before (Figure 4C), at mid-term, only genes directly linked by input TFs showed correlation in their |LFC| (Figures 6D_1_-6D_3_ and Supplementary Figure S32), supporting the previous assumption concerning the kinetics of transcription, translation, and signals propagation via shifts in input TFs numbers (Figure 1).

Meanwhile, in contrast to above, shifting cells from LB_1.0x_ to the richer LB_1.5x_ medium was accompanied by a decrease in the RNAP concentration (Figure 7A), followed by substantial alterations in the RNA populations, with a large number of DEGs and high *μ*_|*LFC*|_ (Figures 7B and 7C, respectively). Also, as previously, in the mid-term, genes with input TFs reacted more strongly (Figure 7C).

**Figure 7.**
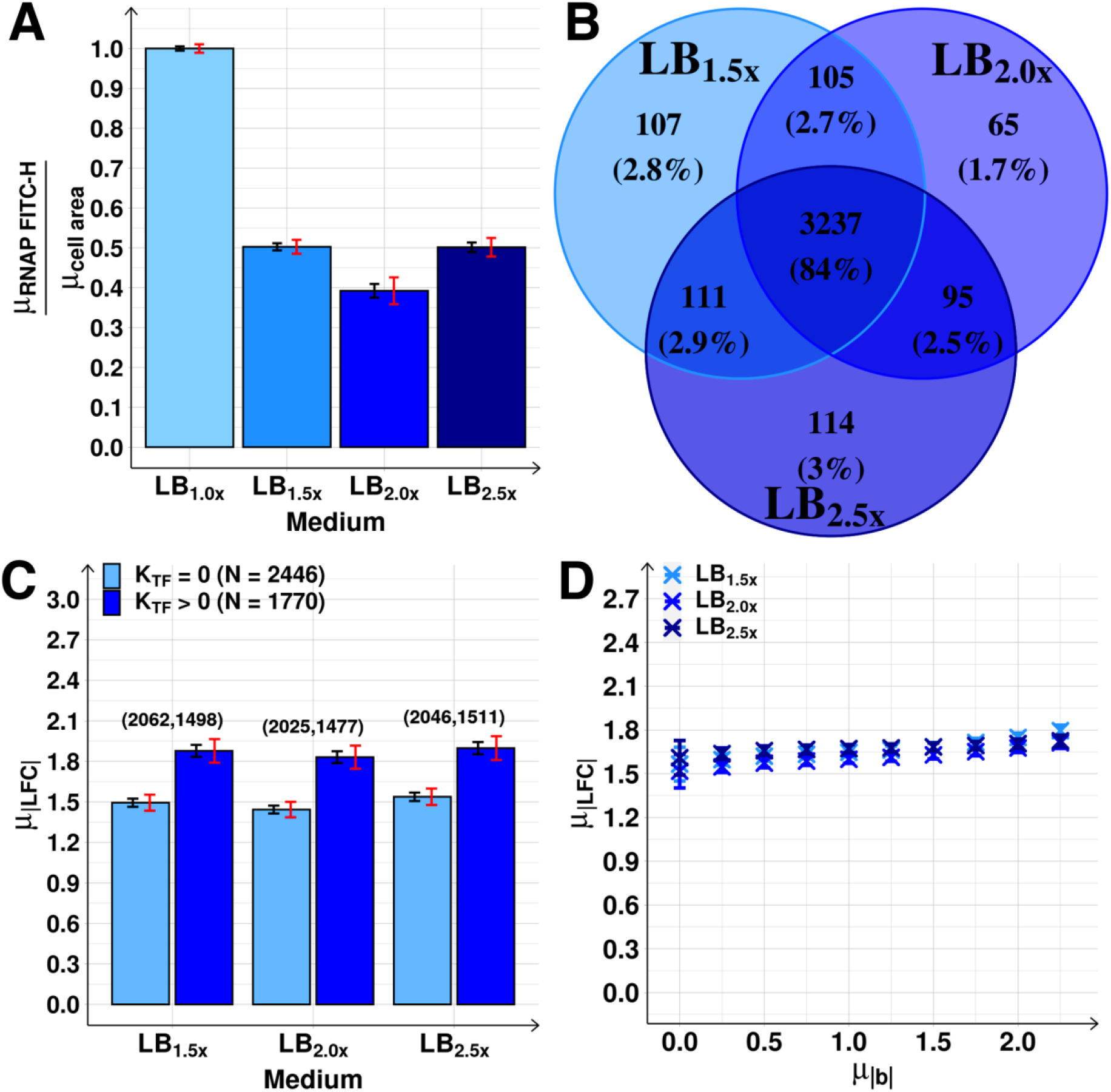
Genome-wide effects of increasing medium richness. **(A)** RNAP concentrations relative to the control, estimated from *μ*_*RNAP FITC* −*H*_ divided by mean cell area (*μ*_*cell area*_). **(B)** Venn diagrams of the DEG. **(C)** *μ*_|*LFC*|_ of *N* genes with K_TF_ equal to and larger than 0, following each medium shift. Above each bar are the number of DEG. **(D)** *μ*_|*LFC*|_ as a function of *μ*_|*b*|_ after the growth-medium shifts. *μ*_|*LFC*|_ obtained using the ensemble approach (Supplementary Results section *Estimation of the expected* 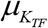 *and μ*_|*b*|_ *using an ensemble approach*, Supplementary Figure S33). Each blue cross is the average outcome from up to 24400 cohorts of 10 genes. In (A) and (C), the black error bars are the SEM and the red error bars are the 95% CB of the SEM. In (D), the small error bars are the SEM (most not visible).

These results support the initial assumption that the changes in RNA abundances follow the RNAP concentration, rather than the medium richness.

### Further increases in medium richness do not decrease RNAP concentration and RNA numbers also do not change

Finally, we further increased growth-medium richness (to LB_2.0x_ and to LB_2.5x_). This caused no significant change in RNAP levels and concentration (Figures 6C and 7A). In agreement with the assumption that the shifts in the RNAP concentration was the cause for the short-term changes in RNA abundances, which then cause the mid-term changes, we observed no significant changes in DEGs or *μ*_|*LFC*|_ at mid-term, when compared with the LB_1.5x_ condition (Figures 7B and 7C, respectively).

Also, as before, 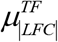 follows *μ*_|*b*|_ (Figures 7D and Supplementary Figure S33) and it does so almost identically in the three perturbations, as expected from the original assumptions (Figure 1).

## DISCUSSION

We investigated if the mid-term responses to genome-wide perturbations of *E. coli*’s TFN are mediated by its topology and logic. We diluted LB medium since this dramatically and reproducibly affects the RNAP concentration (26,27). The increasingly strong nature of the dilutions facilitated the verification of how the RNAP concentration and single-gene, mid-term |LFC|s related. We focused on mid-term transcriptional responses (Figure 1), since short-term responses are unlikely to have been influenced by the TFN due to protein folding and maturation times, etc. Meanwhile, long-term responses were most likely affected by the TFN. However, dissecting them would have been onerous, due to the complicating effects of loss, backpropagation, and coalescence of possibly dozens of signals from origins other than direct input TFs.

Since we lack information on the affinity between each gene and their input TFs, on how the input TFs operate, and on how the *de novo* presence of an input TF alters the binding or activity of other input TFs on the same promoter, we would have failed to predict the behaviour of individual genes with accuracy. As such, we instead predicted the responses of gene cohorts, whose behaviour is less influenced by particular single-gene features (other than the features specific to the cohorts), as these should average out at the cohort level. Further, as in (18), we were only able to correlate *absolute* LFCs of input and output genes (Figure 4B), likely due to limitations in RNA-seq technology and the analysis, and/or missing information on the TFN. Nevertheless, the present information on input TFs and their regulatory effect sufficed to relate the TFN with the genes’ response.

From the RNA-seq data on three time points, we provided evidence that both the TFN and the RNAP affect the results at mid-term (∼180 min), and not before that. In addition, while other factors also influenced genes’ behaviour at mid-term, including single-gene features, they only had minor, local effects. In detail, first, we could not find evidence of GRs (including σ^38^) and (p)ppGpp being material in the global mid-term behaviour (although (p)ppGpp may be significant in the short-term response). Second, we excluded the medium as directly influencing RNA abundances. Third, we excluded global network parameters, other than K_TF_, as being influential as well since none of them correlated to single-gene responses. Fourth, we did not find evidence for significant translational or post-translational regulation, because RNA and protein abundances correlated well, and so did the RNA levels of input TFs and of output genes. Finally, sRNAs did not respond atypically to the RNAP shifts neither in the short-term, nor in in the mid-term.

We have made six key observations on the influence of the logic and topology of the TFN on the mid-term response. First, genes without input TFs were less responsive. Second, the |LFC| of input and output genes correlated positively. Thus, we argue that, on average, input TFs enhanced the |LFC| of individual genes. Third, only nearest neighbour genes in the TFN consistently correlated in |LFC|’s, suggesting that either the effects of the shift in RNAP only reached nearest neighbour genes or they ‘dissipated’ beyond that. However, since the correlations between nearest neighbours were weaker in the short-term than in the mid-term, we expect the first possibility to be more likely. This observation also suggests that there is a degree of genome-wide homogeneity in how long input TF amounts take to change (likely due to physical limitations on the rates constants controlling bacterial gene expression), in agreement with the constraints on timing variability reported in (6). Fourth, the behaviour was orderly (rather than chaotic), with most genes responsive to the weak perturbations also responding to the stronger perturbations, suggesting the existence of features (on genes and/or the TFN) affecting the responsiveness (Supplementary Figure S34). Similarly, there is a good overlap between the sets of genes responsive in the short and in the mid-term, but weak overlap to those responsive prior to the perturbation (Supplementary Figure S35). Fifth, on average, as K_TF_ increased, the correlation between the input and each output gene decreased, which is likely unavoidable and may be a limiting factor in how many input TFs genes can have. Finally, it is *μ*_|*b*|_ that (partially) controls the genes’ responsiveness to the stress, while the apparent relationship between 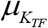 and *μ*_|*LFC*|_ of *μ*_|*b*|_ is only due to the linear correlation between *μ*_|*b*|_ and 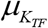. Nevertheless, the possible values *μ*_|*b*|_ are limited by the values of 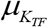

These observations are evidence that the genome-wide responsiveness to this stress depends on the TFN structure, in agreement with past studies (9,20,21,67,80). Expanding this research may thus inform on how to improve the robustness and plasticity of synthetic circuits. Further, as suggested in (20), bacteria subjected to stress, rather than under optimal conditions, may be a better proxy of their state when infecting a host. Thus, imposing stresses may be a valuable strategy to identify new target genes for antibiotics that act by disrupting bacterial adaptability to new conditions.

The use of medium dilution as a genome-wide stress is a good proxy for nutrient imbalance, and we identified ∼900 responsive genes, even for moderate nutritional stress, of which only 58 are essential under optimal conditions. It is plausible that some of the responsive genes, particularly those responsive to all 3 medium dilutions, may be essential to adapt to poorer media, and thus are potential new drug targets. Conversely, it may be possible to tune these genes to assist in the performance of metabolic tasks, without disturbing the basic biology of the cells. As such, they may be appropriate targets for modifications that could improve the yield and sustainability of bio-industrial processes.

Finally, our findings may assist in developing new models of single-gene, mid-term transcriptional responses to genome-wide perturbations, where short-term responses are controlled by single-gene features, while mid-term responses are also influenced by the topology and logic of the TFN. Such large-scale TFN models could be of use in exploring how natural TFNs perform complex transcriptional programs, responsive to large-scale stresses, such as environmental shifts and antibiotics. Further, they may assist in identifying the critical elements of the TFN during stress responses. We hypothesize that the combined regulatory effect of the input TFs of *E. coli* genes (here quantified by *μ*_|*b*|_) is critical in the responses to various different genome-wide stresses. These efforts will be facilitated by ongoing information gathering on single-gene features (34,81-83), including on microorganisms other than *E. coli*.

## SUPPLEMENTARY DATA

**Supplementary Information**. (PDF)

## AUTHOR CONTRIBUTIONS

A.S.R, B.L.B.A, S.M.D.O conceived the study and A.S.R and B.L.B.A directed it. B.L.B.A planned, directed, and executed data analysis, to which C.S.D.P and I.S.C.B contributed to. M.N.M.B, A.S.R, B.L.B.A interpreted and integrated the data. V.C, S.D, V.K performed measurements. A.H, H.T.J and E.D assisted the data analysis. A.H, J.L.P, O.P.S, A.G, M.N, J.K, P.A assisted on RNA-seq. B.L.B.A and A.S.R, assisted by H.T.J, drafted the manuscript, supplementary and figures, which were revised by all co-authors. M.N.M.B, V.C and S.D. contributed equally as second author and have the right to list their name first.

