## Supplementary Information for "The transcription factor network of *E. coli* steers global responses to shifts in RNAP concentration"

### SUPPLEMENTARY RESULTS

#### Expected effects of shifting RNA polymerase concentration on a gene's transcription dynamics

Present knowledge of transcription in *E. coli* suggests that this is a multi-step, highly regulated process, which usually can be well approximated by a two-step model (Reactions S1).

The regulators are usually RNAP, σ factors, and, in many cases, gene-specific input TFs, including global regulators. The two rate constants in (S1) differ with the input TF's concentration. Depending on them, in some conditions, transcription will be blocked, while in others it will be enhanced, compared to a basal rate (1).

According to (S1), an RNAP can find the promoter (*Pro*) of gene $\alpha$, which will be in state $\beta$ at that moment due to a specific set of bound/unbound input TFs, that control the rates of both closed (CC) and subsequent open complex (OC) formations. Supplementary Figure S11A illustrates forms of TF-promoter interactions that affect the propensity of transcription.

Here, when binding to the transcription start site of the promoter region, the RNAP will attempt to form a CC with the DNA (2), via a reversible process. $k_{\alpha,\beta}^{cc,F}$ and $k_{\alpha,\beta}^{cc,B}$ are the forward and backward rate constants of CC formation. Once forming a CC, the RNAP can commit to OC at the rate $k_{\alpha,\beta}^{oc}$, after which initiation is nearly irreversible. It follows elongation (which frees the promoter) and RNA completion, which frees the RNAP (1).

$$Pro_{\alpha}^{\beta(TF_1,...)} + RNAP \underset{k_{\alpha,\beta(TF)}^{cc,B}}{\overset{k_{\alpha,\beta(TF)}^{cc,F}}{\rightleftarrows}} CC_{\alpha}^{\beta(TF_1,...)} \xrightarrow{k_{\alpha,\beta(TF)}^{oc}} Pro_{\alpha}^{\beta(TF_1,...)} + RNAP + RNA_{\alpha} \quad \text{(S1)}$$

The effective rate of transcription, which defines the promoter strength, differs with both $k_{\alpha,\beta}^{cc}$ and $k_{\alpha,\beta}^{oc}$ which then, combined with a reaction for RNA degradation, controls the mean RNA levels. Given the fast degradation rates of RNA (~1-2 min (3)), changes in mean RNA levels are expected to be quick once the transcription rate is altered.

According to (S1), changing RNAP concentration will particularly affect the kinetics of genes with long lasting CC.

Finally, RNA degradation as well as RNA dilution with cell division (reaction S2) is usually modelled as a single-step process since evidence suggests that it can be well fitted by a single exponential function (4). This step is not expected to be subject to significant sequence-specific regulation (3).

$$RNA_{\alpha} \xrightarrow{k_{\alpha}^{\deg}+k_{\alpha}^{dil}} \varnothing \quad \text{(S2)}$$

where $k_{\alpha}^{deg}$ is the RNA degradation rate and $k_{\alpha}^{dil}$ is the RNA dilution rate.

Given the above, shifting RNA polymerase (RNAP) concentration should quickly change the RNA abundances of many genes, particularly those whose step of closed complex formation has a significant time length. The shifts in RNA abundances then propagate to protein abundances, via translation.

Other sources of variability in single-gene response will then emerge from single-TF properties, such as different input TFs binding affinities, and folding and maturation times, etc.

This model is presented here to facilitate the interpretation of the results, and it does not intend to be a strick representation of *all* events occurring during the genome-wide stresses. As an example, this model would fail to capture the influence of ppGpp(p), non-coding RNAs, positive supercoiling buildup, events during transcription, and post-transcription and post-translation regulation, among other.

**Estimation of $\mu_{K_{TF}}$ and $\mu_{|b|}$ from empirical data using an ensemble approach**

The behavior of networks as a function of specific topological features can be studied using an ensemble approach. This approach consists of generating model networks imposing some features (e.g., total number of connected nodes) in order to study their relevance, while other features are randomly generated (5) to decrease the chances that the conclusions generalize poorly. This assists the study of responses to a global perturbation, since it is complex to filter abnormal responses, due to lack of knowledge about the range of possible behaviors. Also, it is not easy to establish if the 'abnormality' is in an arbitrary local topological feature of the genes of interest, or in the response due to an arbitrary internal parameter. Overall, we expect this methodology to decrease the effects of arbitrary features (but not necessarily all effects or for all features).

In our hypothesis, the perturbation strength due to shifting RNAP concentration ($\mu_{|LFC|}$) on a gene is a function of (and, thus, can be predicted from) $\mu_{|b|}$ of the input TFs of that gene. However, since other 'local' factors can affect the $\mu_{|LFC|}$ of a gene (including its original state, its sensitivity to supercoiling build up, etc., we only expect our hypothesis to be true at the level of gene cohorts.

Similarly, for large enough gene cohorts, we hypothesize that $\mu_{|LFC|}$ can be predicted from $\mu_{|b|}$ alone. Thus, we compared the behavior of cohorts of genes differing in $|b|$ using an ensemble approach. For this, we proceeded by comparing genes differing in $|b|$. Thus, we randomly selected genes, from the empirical data, to form cohorts with a given $\mu_{|b|}$. We then compared the average behavior (i.e., $\mu_{|LFC|}$) of these sets of cohorts with a given $\mu_{|b|}$.

This was done in two ways (Supplementary Figure S30 and Figure 5B respectively). In one case, we assembled cohorts of genes with a given $\mu_{K_{TF}}$ (average $K_{TF}$), which obligatorily results in a given $\mu_{|b|}$ (Figure 5C). In the other, we assembled cohorts of genes with a given $\mu_{|b|}$. We found no difference in the results using the two ensemble methods (note the similarities between Supplementary Figure S30 ($LB_{0.75x}$) and Figure 5B ($LB_{0.75x}$), between S30 ($LB_{0.5x}$) and 5B ($LB_{0.5x}$) and, between S30 ($LB_{0.25x}$) and 5B ($LB_{0.25x}$), respectively). In detail, to obtain each of the 46 data points in Supplementary Figure S30, we randomly assembled 100000 cohorts of 10 genes each with specific $\mu_{K_{TF}}$ (and, thus, $\mu_{|b|}$). This was done 100 times, from which we obtained average values and error bars for each of the 46 data points. The same was done for 5B, except that the cohorts, as noted, were assembled directly based on their $\mu_{|b|}$.

The pseudo-algorithm (Algorithm 1) created in MATLAB to generate, using the ensemble approach, the empirical data points in Supplementary Figure S30 was:

1. For each RNAP shift, let **LFC_data** be the **|LFC|** (absolute of log2 of fold change) of each gene and **KTF_data** be the corresponding number of input TFs, i.e., $\mathbf{K_{TF}}$, from 0 to 5.
2. Get the corresponding **S_data**, i.e., $S = |b|$, the absolute of the sum of the regulatory effects of the inputs of each gene.
3. Get the fraction of genes, $\mathbf{p_i}$, with a given $\mathbf{K_{TF}^i}$ ($\mathbf{K_{TF}}$ = i), with i=0:5.
4. Sample $\mathbf{b_i}$ from a Beta PDF (Probability Density Function), for each value of $\mathbf{p_i}$, using the MATLAB function 'BetaPDF' with parameter values: a = b = 0.75.
5. Let **r** (set to 10000) be the number of target genes, with a given $\mathbf{K_{TF}^i}$, to be randomly sampled with replacement.
6. For each $\mathbf{K_{TF}^i}$, define $\mathbf{n_i} = \mathbf{b_i} \times \mathbf{r}$, and create a set of $\mathbf{n_i}$ genes with $\mathbf{K_{TF}^i}$, randomly sampled with replacement from **KTF_data**. Let all these new sets of values be named **new_KTF_data**.
7. Let **TM** be numbers from 0.5 to 5, with an increment of 0.1, and a precision of 0. Let each **TM** value be a 'target mean of $\mathbf{K_{TF}}$'.
8. To find cohorts of genes with a given value of **TM**, set the number of iterations to 100. In each iteration, named **h,** do as follows:
    a. For each $\mathbf{K_{TF}^i}$**,** sample with replacement from **KTF_data** a vector **index_i** with $\mathbf{n_i}$ indexes of genes with $\mathbf{K_{TF}}$ **=** i.
    b. Generate **new_S_data** combining the empirical **S**, from **S_data**, of all genes under the indexes found in all sets of **index_i**. The data is added in ascending order of sorted $\mathbf{K_{TF}}$, in accordance with point 3.

c. Generate, for each RNAP shift, **new_LFC_data** combining the empirical **|LFC|**, from **LFC_data**, of all genes under the indexes found in all sets of **index_i**. The data is added by in ascending order of sorted $K_{TF}$, in accordance with point 3.

d. Set **m**, the target number of genes of cohorts with a given value of **TM**, to 10.

e. Let the number of sampling iterations be 100000. In each iteration **j**, do as follows:

   i. In **new_sampledKTF_data** save **m** values sampled with replacement from **new_KTF_data**. Save in '**I(:, j)'** the indexes of the **m** values sampled.

   ii. In '**M_KTF (1, j)'** save the mean value of **new_sampledKTF_data**.

f. For each value **k** in **TM** located in position **t**:

   i. Save in **idx_mean** the indexes in **M_KTF** of all mean values equal to **k** with a precision of 0.

   ii. Get the indexes **I(:, idx_mean)** of the genes included in all sampling sets whose mean is **k**.

   iii. For each RNAP shift, calculate the mean **M_LFC(t, h)** of **|LFC|** from the data in **new_LFC_data** of the genes in **I(:, idx_mean)**.

   iv. Calculate the mean **M_S(t, h)** of **S** in **new_S_data** of the genes in **I(:, idx_mean)**.

9. For each RNAP shift, get the expected mean **|LFC| M_LFC_final(t)** and **SEM_LFC_final(t)** for each **k** value in position **t** in **TM**. **M_LFC_final(t) and SEM_LFC_final(t)** are calculated, respectively, as the mean and standard deviation of the **M_LFC(t, h)** values over the **h** runs.
10. Get the expected mean **S M_S_final(t)** and **SEM_S_final(t)** for each **k** in position **t** in **TM**. **M_S_final(t)** and **SEM_S_final(t)** are calculated, respectively, as the mean and standard deviation of the **M_S(t, h)** values over the **h** runs.

Meanwhile, for Figure 5B, the pseudo-algorithm (Algorithm 2) used was:

1. For each RNAP shift, let **LFC_data** be the **|LFC|** (absolute of log2 of fold change) of each gene and **KTF_data** be the corresponding number of input TFs, i.e., $K_{TF}$, from 0 to 5.
2. Get the corresponding **S_data**, i.e., $S = |b|$, the absolute of the sum of the regulatory effects of the inputs of each gene.
3. Get the fraction of genes, $p_i$, with a given $S_i$ (**S** = i), for all unique values found **S_data** in ascending order.
4. Sample $b_i$ from a Beta PDF, for each value of $p_i$, using the MATLAB function 'BetaPDF' with parameter values: a = b = 1.5.

5. Let **r** (set to 10000) be the number of target genes, with a given $\mathbf{S_i}$, to be randomly sampled with replacement.
6. For each $\mathbf{S_i}$, define $\mathbf{n_i} = \mathbf{b_i} \times \mathbf{r}$, and create a set of $\mathbf{n_i}$ genes with $\mathbf{S_i}$, randomly sampled with replacement from **S_data**. Let all these new sets of values be named **new_S_data**.
7. Let **TM** be numbers from 0 to 3 with an increment of 0.25, and a precision of ± 0. Let each **TM** value be a 'target mean of **S**'.
8. To find cohorts of genes with a given value of **TM**, set the number of iterations to 100. In each iteration, named **h,** do as follows:
    a. For each value of $\mathbf{S_i}$**,** sample with replacement from **S_data** a vector **index_i** with $\mathbf{n_i}$ indexes of genes with **S =** i.
    b. Generate, for each RNAP shift, **new_LFC_data** combining the empirical **|LFC|**, from **LFC_data**, of all genes under the indexes found in all sets of **index_i**. The data is added in ascending order of sorted **S**, in accordance with point 3.
    c. Set **m**, the target number of genes of cohorts with a given value of **TM**, to 10.
    d. Let the number of sampling iterations be 100000. In each iteration **j**, do as follows:
        i. In **new_sampledS_data** save **m** values sampled with replacement from **new_S_data**. Save in **I(:, j)** the indexes of the **m** values sampled.
        ii. In **M_S (1, j)** save the mean value of **new_sampledS_data**.
    e. For each value **k** in **TM** located in position **t**:
        i. Save in **idx_mean** the indexes in **M_S** of all mean values equal to **k** with a precision of ± 0.1.
        ii. Get the indexes **I(:, idx_mean)** of the genes including in all the sampling sets whose mean is **k.**
        iii. For each RNAP shift, calculate the mean **M_LFC(t, h)** of **|LFC|** from **new_LFC_data** of the genes found in **I(:, idx_mean)**.
9. For each RNAP shift, get the expected mean **|LFC| M_LFC_final(t)** and **SEM_LFC_final(t)** for each **k** value in position **t** in **TM**. **M_LFC_final(t) and SEM_LFC_final(t)** are calculated, respectively, as the mean and standard deviation of the **M_LFC(t, h)** values over the **h** runs.

We also created tailored cohorts with imposed values of $\mu_{K_{TF}}$ and $\mu_{|b|}$, to evaluate how $K_{TF}$ and $|b|$ independently affect |LFC|. For instance, for Figure 5E, we obtained cohorts with a given $\mu_{K_{TF}}$ and different $\mu_{|b|}$. For this, we used the following pseudo-algorithm (Algorithm 3, obtained by "inserting" algorithm 2 into algorithm 1):

1. For each RNAP shift, let **LFC_data** be the **|LFC|** (absolute of log2 of fold change) of each gene and let **KTF_data** be the corresponding number of input TFs, i.e., $\mathbf{K_{TF}}$, from 0 to 5.
2. Get the corresponding **S_data**, i.e., $S = |b|$, the absolute of the sum of the regulatory effects of the inputs of each gene.
3. Get the fraction of genes, $\mathbf{p_i}$, with a given $\mathbf{K_{TF}^i}$ ($\mathbf{K_{TF}}$ = i), with i=0:5.
4. Sample $\mathbf{b_i}$ from a Beta probability density function, for each value of $\mathbf{p_i}$, using the MATLAB function 'BetaPDF' with parameter values: a = b = 0.75.
5. Let **r** (set to 10000) be the number of target genes, with a given $\mathbf{K_{TF}^i}$, to be randomly sampled, with replacement.
6. For each $\mathbf{K_{TF}^i}$, define $\mathbf{n_i} = \mathbf{b_i}$ x **r**, and create a set of $\mathbf{n_i}$ genes with $\mathbf{K_{TF}^i}$, randomly sampled with replacement from **KTF_data**. Let these new sets of values be named **new_KTF_data**.
7. Let **TM** be numbers from 0.5 to 5, with an increment of 0.1, and a precision of 0. Let each **TM** value be a 'target mean of $\mathbf{K_{TF}}$'.
8. To find and analyze cohorts of genes with a given value of **TM**, set the number of iterations to 100. In each iteration, named **h,** do as follows:
   a. For each value $\mathbf{K_{TF}^i}$ **,** sample with replacement from **KTF_data** a vector **index_i** with $\mathbf{n_i}$ indexes of genes with $\mathbf{K_{TF}}$ **=** i.
   b. Generate **new_S_data{1,h}** combining the empirical **S**, from **S_data**, of all genes under the indexes found in all sets of **index_i**. The data is added in ascending order of sorted $\mathbf{K_{TF}}$, in accordance with point 3.
   c. Generate, for each RNAP shift, **new_LFC_data{1, h}** combining the empirical **|LFC|**, from **LFC_data**, of all genes under the indexes found in all sets of **index_i**. The data is added by in ascending order of sorted $\mathbf{K_{TF}}$, in accordance with point 3.
   d. Set **m**, the target number of genes of cohorts with a given value of **TM**, to 10.
   e. Let the number of sampling iterations be 100000. In each iteration **j**, do as follows:
      i. In **new_sampledKTF_data** save **m** values sampled with replacement from **new_KTF_data**. Save in **I(:, j)** the indexes of the **m** values sampled.
      ii. In **M_KTF (1, j)** save the mean value of **new_sampledKTF_data**.
   f. For each value **k** in **TM** located in position **t**:
      i. Save in **idx_mean** the indexes in **M_KTF** of all mean values equal to **k** with a precision of 0.
      ii. In **genes_sampled_KTF_k{1, h}** save the indexes **I(:, idx_mean)** of the genes included in all sampling sets whose mean is **k.**

**iii.** Save in **M_KTF_all_data{t, h}** all values from **new_KTF_data(genes_sampled_KTF_k{1, h})**, i.e., all the $\mathbf{K_{TF}}$ values contained in **new_KTF_data** of the genes under the indexes saved in **genes_sampled_KTF_k{1, h}**.

**iv.** In **M_S_all_data{t, h}** save all the **S** values contained in **new_S_data{1, h}(genes_sampled_KTF_k{1, h})**.

v. For each RNAP shift, save in **M_LFC_all_data{t, h}** all the **|LFC|** values from the respective **new_LFC_data{1, h}(genes_sampled_KTF_k{1, h})**.

vi. Let **Unique_S** be the unique values of **S** found in **M_S_all_data{t, h}**.

vii. In **S_data_S** save all the values in **M_S_all_data{t, h}** in ascending order of sorted **S**.

viii. For each RNAP shift and each value $\mathbf{Unique\_S^i}$ (**Unique_S**=i), obtain $\mathbf{M\_LFC\_all\_data\{t,h\}(M\_S\_all\_data\{t, h\}=Unique\_S^i)}$. Let all these sets of **|LFC|** be named **LFC_data_S** for each shift.

ix. Get the corresponding **KTF_data_S** for each value $\mathbf{Unique\_S^i}$ from $\mathbf{M\_KTF\_all\_data\{t, h\}(M\_S\_all\_data\{t, h\}=Unique\_S^i)}$.

x. Get the fraction of genes, $\mathbf{p_i\_S}$, with a given $\mathbf{Unique\_S^i}$, for all unique values found **S_data_S** in ascending order.

xi. Sample $\mathbf{b_i\_S}$ from a Beta PDF (MATLAB function ‘BetaPDF’ with a = b = 0.75), for each value of $\mathbf{p_i\_S}$.

xii. Let **r_S** (set to 10000) be the number of target genes, with a given $\mathbf{Unique\_S^i}$, to be randomly sampled with replacement.

xiii. For each $\mathbf{Unique\_S^i}$, define $\mathbf{n_i\_S} = \mathbf{b_i\_S} \times \mathbf{r\_S}$, and create a set of $\mathbf{n_i\_S}$ genes with $\mathbf{Unique\_S^i}$, randomly sampled with replacement from **S_data_S**. Let all these new sets of values be named **new_S_data_S**.

xiv. Let **TM_S** be numbers from 1 to the maximum value of **Unique_S**, with a precision of ± 0. Let each **TM_S** value be a ‘target mean of **S**’ given the values in **Unique_S**.

xv. To find cohorts of genes with a given value of **TM_S**, set the number of iterations to 10. In each iteration, named **h_S,** do as follows:

1. For each value $\mathbf{Unique\_S^i}$, sample with replacement from **S_data_S** a vector **index_i_S** with $\mathbf{n_i\_S}$ indexes of genes with $\mathbf{Unique\_S^i}$.
2. Generate, for each RNAP shift, **new_LFC_data_S** combining the empirical **|LFC|**, from **LFC_data_S**, of all genes under the indexes found in all sets of **index_i_S**. The data is added in

ascending order of sorted **Unique_S**, in accordance with point f.x.

3. Get the corresponding **new_KTF_data_S**, from **KTF_data_S**, of all genes under the indexes found in all sets of **index_i_S**. The data is added in ascending order of sorted **Unique_S**, in accordance with point f.x.
4. Set **m_S**, the target number of genes of cohorts with a given value of **TM_S**, to 10.
5. Let the number of sampling iterations be 100000 and, in each iteration **j_S**, do as follows:
   a. In **new_sampledS_data_S** save **m_S** values sampled with replacement from **new_S_data_S**. Save in **I_S(:, j_S)** the indexes of the **m_S** values sampled.
   b. In **M_UNIQUE_S(1, j_S)** save the mean value of **new_sampledS_data_S**.
6. For each value **k_S** in **TM_S** located in position **t_S**:
   a. Save in **idx_mean_S** the indexes in **M_UNIQUE_S** of all mean values equal to **k_S** with a precision of ± 0.1.
   b. Get the indexes **I_S(:, idx_mean_S)** of the genes including in all the sampling sets whose mean is **k_S**.
   c. In **M_S_S(t_S, h_S)** save the mean of **S** from **new_S_data_S** of the genes found in **I_S(:, idx_mean_S)**.
   d. For each RNAP shift, calculate the mean **M_LFC_S(t_S, h_S)** of **|LFC|** from **new_LFC_data_S** of the genes found in **I_S(:, idx_mean_S)**.
   e. Calculate the mean **M_KTF_S(t_S, h_S)** of **$K_{TF}$** from **new_KTF_data_S** of the genes in **I_S(:, idx_mean_S)**.

xvi. For each RNAP shift, save under **M_LFC_final(t,t_S,h)** the expected mean and SEM of **|LFC|** for each **k_S** value in position **t_S** in **TM_S**. **M_LFC_final(t,t_S,h)** is calculated as the mean of the **M_LFC_S(t_S, h_S)** values over the **h_S** runs.

xvii. Save under **M_S_final(t,t_S,h)** the expected mean **S** for each **k_S** in position **t_S** in **TM_S**. **M_S_final(t,t_S,h)** is calculated as the mean of the **M_S_S(t_S, h_S)** values over the **h_S** runs.

xviii. Save under **M_KTF_final(t,t_S,h)** the expected mean $\mathbf{K_{TF}}$ for each **k_S** in position **t_S** in **TM_S**. **M_KTF_final(t,t_S,h)** iscalculated as the mean of the **M_KTF_S(t_S, h_S)** values over the **h_S** runs.

9. For each position **t_S** corresponding to a given **k_S** value, do as follows:
   a. For each RNAP shift, save under **M_LFC_ks** and **SEM_LFC_ks**, respectively, the expected mean and SEM of **|LFC|** for each **k_S** value in position **t_S**. **M_LFC_ks** and **SEM_LFC_ks** are calculated, respectively, as the mean and standard deviation of the **M_LFC_final(t,t_S,h)** values over the **h** runs.
   b. Save under **M_S_ks** and **SEM_S_ks**, respectively, the expected mean and SEM of **S** for each **k_S** in position **t_S**. **M_S_ks** and **SEM_S_ks** are calculated, respectively, as the mean and standard deviation of the **M_S_final(t,t_S,h)** values over the **h** runs.
   c. Save under **M_KTF_ks** and **SEM_KTF_ks**, respectively, the expected mean and SEM of $\mathbf{K_{TF}}$ for each **k_S** in position **t_S**. **M_KTF_ks** and **SEM_KTF_ks** are calculated, respectively, as the mean and standard deviation of the **M_KTF_final(t,t_S,h)** values over the **h** runs.

Finally, from the above, one can produce the results in Figure 5B. In detail, we assembled gene cohorts (Supplementary Results section *Estimation of the expected* $\mu_{K_{TF}}$ *and* $\mu_{|b|}$ *using an ensemble approach*) with a given $\mu_{|b|}$ (up to 24500 cohorts of 10 genes per $\mu_{|b|}$ value). In Figure 5B, each blue cross is the $\mu_{|LFC|}$ of a set of cohorts. Visibly, $\mu_{|LFC|}$ increases with $\mu_{|b|}$.

### SUPPLEMENTARY FIGURES

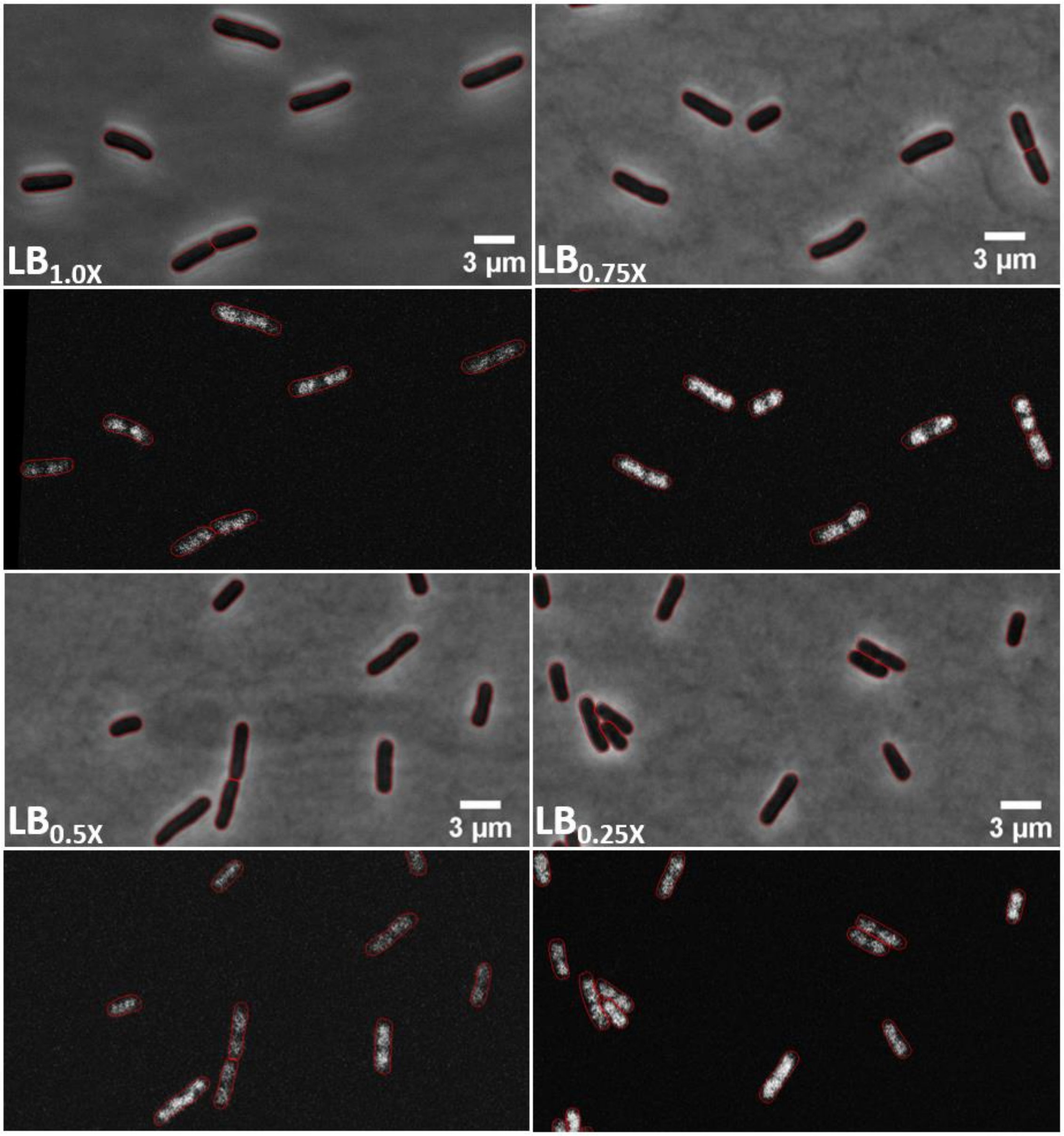

**Figure S1. Example phase-contrast images (first and third rows) and corresponding confocal images of *E. coli* cells (second and forth rows, respectively).** RL1314 cells grown in various media (Methods sections *Bacterial strains, media, growth conditions and curves* and *Microscopy*). Cells were segmented in phase-contrast images using CellAging (6).

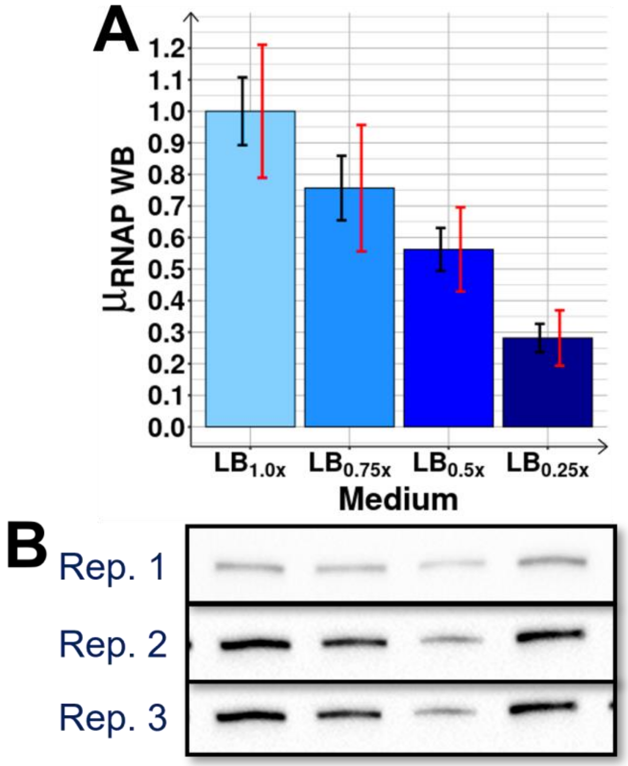

**Figure S2. (Related to Figure 2E) RNAP protein levels by Western Blot following medium dilutions from $LB_{1.0x}$ to $LB_{0.75x}$, $LB_{0.5x}$ and $LB_{0.25x}$.** For each diluted medium, the RNAP protein levels were measured by Western Blot (Methods section *Protein isolation and western blotting*). The absolute values were extracted from the blot images by the 'Image Lab' software (version 5.2.1), from which the relative values were calculated. Supplementary Table S2 shows the normalized intensity volumes for RNAP levels. (A) Relative values. (B) Blot images of 3 biological replicates (Rep). All lines are located at the 155 kDa level, corresponding to the β' unit ((7), first reported in (8)).

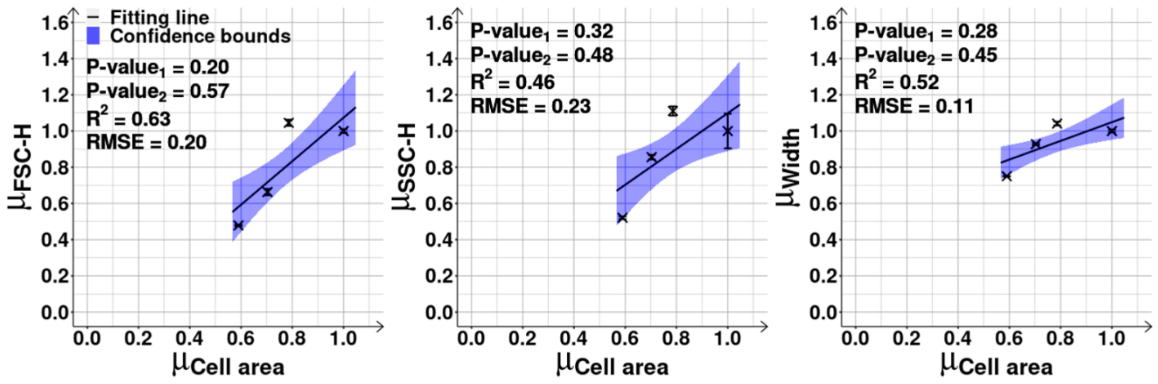

**Figure S3. Related to Figures 2F and 2G) Flow-cytometry and microscopy data on cell size and composition.** Correlation plot between cell areas ($\mu_{cell\ area}$, Methods section *Microscopy*) and each of the three flow-cytometry parameters positively correlated to cell size and composition ($\mu_{FSC-H}$, $\mu_{SSC-H}$, $\mu_{Width}$ from the FCS-H, SSC-H and Width parameters, respectively, Methods section *Flow-cytometry*), in each condition, 180 min after shifting the medium. All data points were obtained from 3 independent biological replicates and are shown relative to $LB_{1.0x}$ (control). Mean cell areas were obtained from phase-contrast images of MG1655 cells grown in the same conditions (Methods sections *Bacterial strains, media, growth conditions and curves* and *Microscopy*). The best fitting lines (solid black), their 68% confidence bounds (blue shadow areas) and statistics (coefficient of determination ($R^2$), root mean square error (RMSE), and P-values at 0.1 significance level) were obtained as described in Methods section *Statistical tests c*.

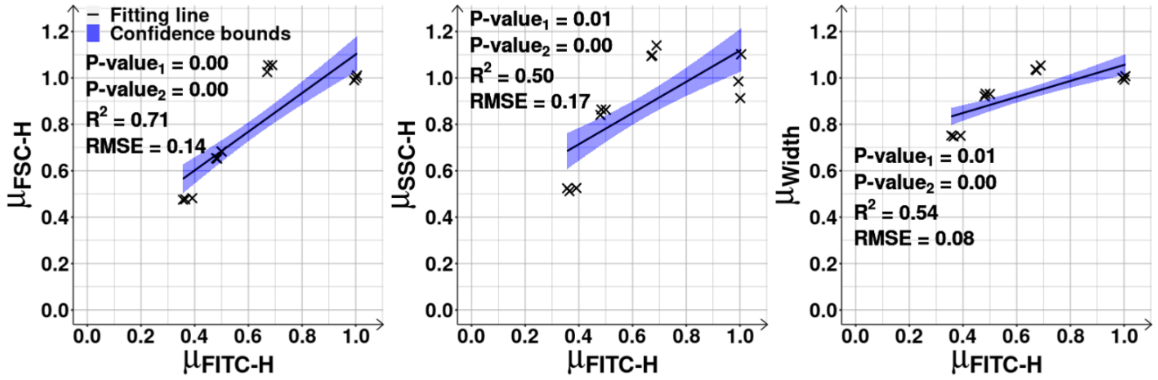

**Figure S4. (Related to Figures 2G and 2D) Flow-cytometry data relating cell size and composition with the expression levels of the RpoC sub-unit of RNAP.** Correlation plots between FITC-H (maximum peak 'Height', -H, of the FITC signal), which is linearly correlated to RNAP-GFP levels ($\mu_{FITC-H}$), and each of the three flow-cytometry parameters positively correlated to cell size and composition ($\mu_{FSC-H}$, $\mu_{SSC-H}$, $\mu_{Width}$ from the FCS-H, SSC-H and Width parameters), in $LB_{1.0x}$, $LB_{0.75x}$, $LB_{0.5x}$ and $LB_{0.25x}$, at 180 min. All data is relative to $LB_{1.0x}$ (control). Mean background fluorescence levels were removed from the FITC-H signals (Methods section *Flow-cytometry*). The best fitting lines (solid black) along with their 68% confidence bounds (blue shadow areas) and statistics (coefficient of determination ($R^2$), root mean square error (RMSE), and P-values at 0.1 significance level) were obtained as described in Methods section *Statistical tests c*.

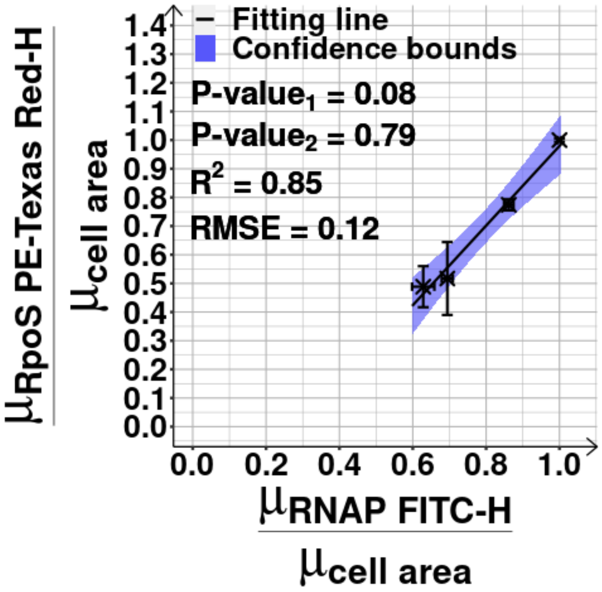

**Figure S5. (Related to Figures 2I and 2D) Mean concentration of RpoS plotted against the mean concentration of RNAP.** The mean concentration of of RpoS ($\sigma^{38}$) tagged with mCherry in $LB_{1.0x}$, $LB_{0.75x}$, $LB_{0.5x}$ and $LB_{0.25x}$ at 180 min, was measured by mean single-cell fluorescence by flow-cytometry (PE-Texas Red-H parameter), over mean cell area ($\mu_{cell\ area}$) (Methods section *Microscopy*). The mean RNAP concentration of RL1314 cells was obtained by mean single-cell fluorescence (FITC-H parameter, Methods section *Flow-cytometry*) over mean cell area ($\mu_{cell\ area}$). Mean background fluorescence levels were removed from both FITC-H and PE-Texas Red-H signals (Methods section *Flow-cytometry*). All data is relative to $LB_{1.0x}$ (control). The best fitting line (solid black) along with its 68% confidence bounds (blue shadow area) and statistics (coefficient of determination ($R^2$), root mean square error (RMSE), and P-values at 0.1 significance level) were obtained as described in Methods section *Statistical tests c*.

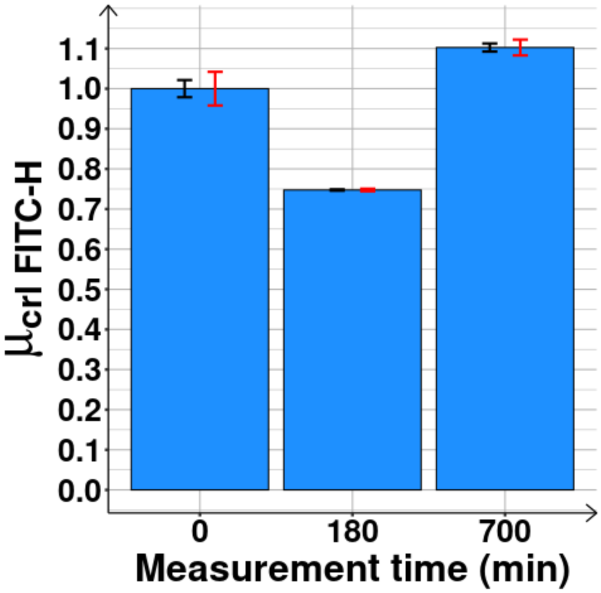

**Figure S6. (Related to Figures 2I) crl gene expression levels in LB$_{0.5x}$.** Mean protein expression levels ( $\mu_{crl\ FITC-H}$ ) of the crl gene, first, at 0 and then at 180 (mid-log phase) and 700 (stationary growth phase) min in the LB$_{0.5x}$ medium, following dilution. The mean values are shown relative to 0 min.

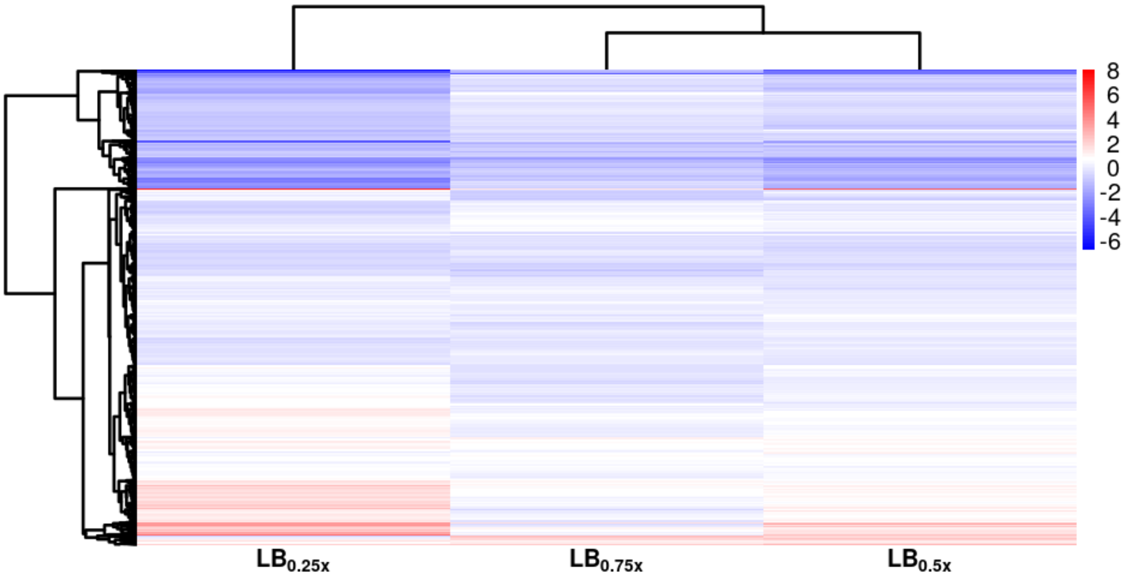

**Figure S7. (Related to Figure 3B) Heatmap of shifts in RNA levels**. Log2 of fold changes (LFC) from RNA-seq following medium dilutions from $LB_{1.0x}$ to $LB_{0.75x}$, $LB_{0.5x}$ and $LB_{0.25x}$. The compressed data includes all genes. The perturbation inflicted by $LB_{0.25x}$ was significantly higher than by $LB_{0.75x}$ and $LB_{0.5x,}$ regarding the number of genes perturbed and their response strengths.

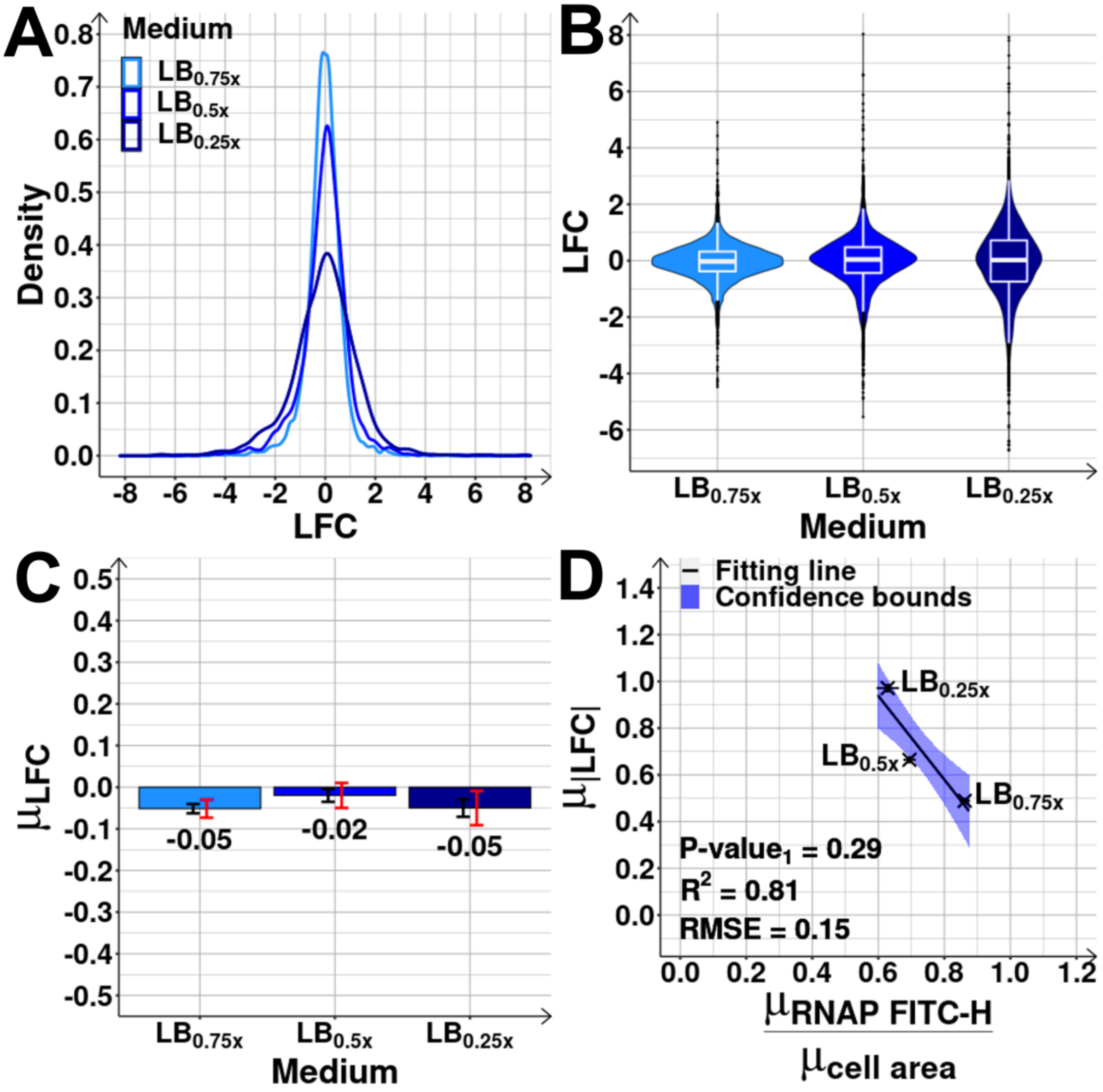

**Figure S8. (Related to Figure 3B) Shifts in RNA levels**.

**(A)** Kernel density estimates (Probability Distribution Function, PDF) of the distribution of genes with a given LFC (log2 of fold change) of RNA abundances, following the shifts from $LB_{1.0x}$ to $LB_{0.75x}$, $LB_{0.5x}$ and $LB_{0.25x}$, respectively. Non-parametric estimates of the PDF were obtained from data on all genes.

**(B)** The violin plot shows the maximum, minimum, median, interquartile ranges and probability density of the distributions in (A).

**(C)** Mean shifts in RNA levels, $\mu_{LFC}$ , following each shift. Black error bars represent the standard error of the mean (SEM), while red error bars represent the 95% confidence bounds of the SEM. From these, the mean cannot be distinguished between the conditions. We also performed 2-

sample T-tests of statistical significance between the distributions. Results in Supplementary Table S3 show that the $LB_{0.75x}$ and $LB_{0.5x}$ cannot be distinguished in a statistical sense.

**(D)** Correlation plot of $\mu_{|LFC|}$ and the respective RNAP concentration shift. All values are relative to the control condition. The best fitting line (solid black) along with its 68% confidence bounds (blue shadow area) and statistics (coefficient of determination ($R^2$), root mean square error (RMSE), and P-values at 0.1 significance level) were obtained as described in Methods section *Statistical tests c*.

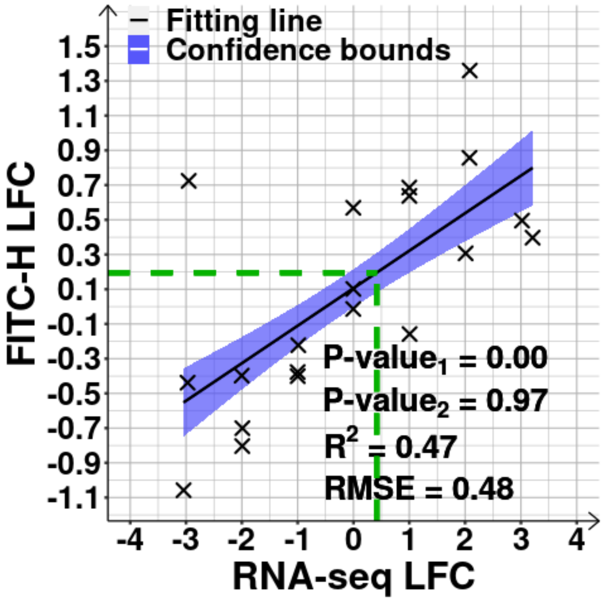

**Figure S9. (Related to Figure 3B) Log$_2$ fold changes (LFCs) in RNA abundances of 20 randomly selected genes covering the whole spectrum of fold changes observed, plotted against their corresponding LFCs in protein abundances**. Data from the shift from $LB_{1.0x}$ to $LB_{0.25x}$. We selected trios of genes whose LFC by RNA-seq was closest to -3, -2, -1, 0, +1, +2, and +3, respectively, to represent the entire spectrum (Methods section *RNA-seq e*). Events considered to be outliers by Tukey's fences (9) were removed. The LFCs in protein abundances were obtained by flow-cytometry (YFP strain library). Mean background fluorescence was removed (Methods section *Flow-cytometry*). The dashed green lines at positions (x, y) = (0.42, 0.20) indicate the average |LFC| of genes whose False Discovery Rate > 0.05, along with the corresponding estimated |LFC| by flow-cytometry (based on the fitting line). The best fitting line (solid black) along with its 68% confidence bounds (blue shadow area) and statistics (coefficient of determination ($R^2$), root mean square error (RMSE), and P-values at 0.1 significance level) were obtained as described in Methods section *Statistical tests c*. Supplementary Table S4 shows the list of strains measured by flow-cytometry.

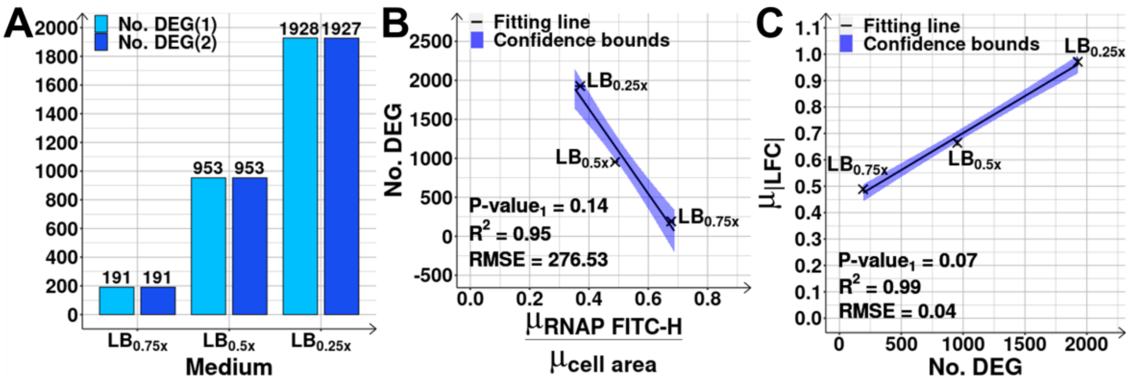

**Figure S10. (Related to Figures 3B and 3C) Number of differentially expressed genes and its correlation with the shift in RNAP concentration.**

**(A)** DEG(1) stands for Differentially Expressed Genes (4029 evaluated) (i.e., genes with False Discovery Rate (FDR) < 0.05), after diluting the medium to $LB_{0.75x}$, $LB_{0.5x}$ and $LB_{0.25x}$ (Methods section *RNA-seq c*). DEG(2) stands for genes whose FDR < 0.05 and absolute log2 of fold change (|LFC|) > 0.4248 ($LB_{0.75x}$), > 0.4085 ($LB_{0.5x}$) or > 0.4138 ($LB_{0.25x}$) (Methods section *RNA-seq d*). As the results using DEG(1) and DEG(2) slightly differ, from here onwards, DEG are selected using the more stringent criteria DEG(2).

**(B)** Number of DEG for each shift plotted against the respective RNAP concentration shift, estimated from the ratio between mean RNAP levels measured by FITC-H, $\mu_{RNAP\ FITC-H}$ using RL1314 cells (Methods section *Flow-cytometry*), and the mean cell area ($\mu_{cell\ area}$) estimated from phase-contrast images of MG1655 cells (Methods section *Microscopy*). All values are relative to the control condition ($LB_{1.0x}$).

**(C)** Correlation plot of the mean of |LFC| (absolute log2 fold change), $\mu_{|LFC|}$, and the respective number of DEG in each shift. The best fitting lines (solid black) along with their 68% confidence bounds (blue shadow area) and statistics (coefficient of determination ($R^2$), root mean square error (RMSE), and P-values at 0.1 significance level) were obtained as described in Methods section *Statistical tests c*.

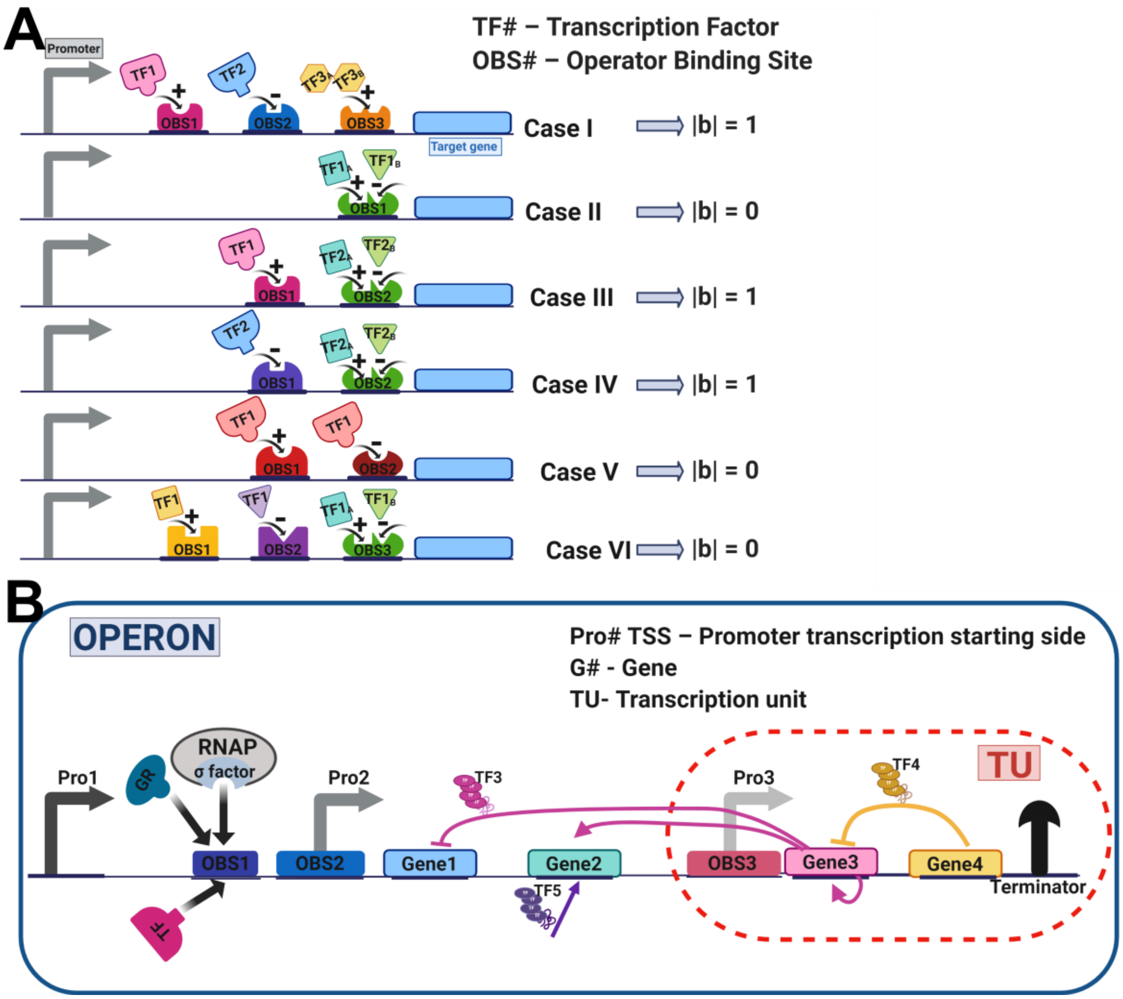

**Figure S11. An example operon and examples of interactions of input transcription factors (TFs) with a total bias of 0 or 1.**

**(A)** Illustrations of 6 forms of transcription regulation by sets of input TFs of an output protein. The input TFs regulation can be to activate (+1), repress (-1), or have a dual outcome (+1 or -1, not considered here due to their rarity). In the latter, the dual functionality is made possible by input TFs with binding affinity to different operator binding sites (OBS) and/or with dual functionality on the same OBS (i.e., can activate or repress a gene depending on the conditions, e.g., TF conformation, concentration and binding affinity, binding of other input TFs, OBS position, etc.). The right side of the figure shows the expected bias, i.e., overall regulatory effect (‘$r$’) of the set input TFs ($|b|$), set to equal the sum of the individual regulatory effect of each input TF, ‘$r$’ (shown by the black sign ‘+’ or ‘-‘, above each input TF on the left side). E.g., in case I, the red OBS has a +1 effect, the blue has a -1 effect, and the orange has a +1 effect. Thus, $|b|$ = |+1-1+1| = +1.

**(B)** Illustration of an operon expressing 4 genes (G1, G2, G3 and G4) under the control of 2 promoters (Pro1 and Pro2). The colored arrows show the different regulations inside this operon. Also illustrated is a transcription unit (TU) inside the operon, controlled by one promoter, Pro3. The vertical black arrow at the end is a terminator while TF5 is produced by a gene not in the operon. This example assumes the definitions of operon and TU in RegulonDB. Figures (A) and (B) created with BioRender.com.

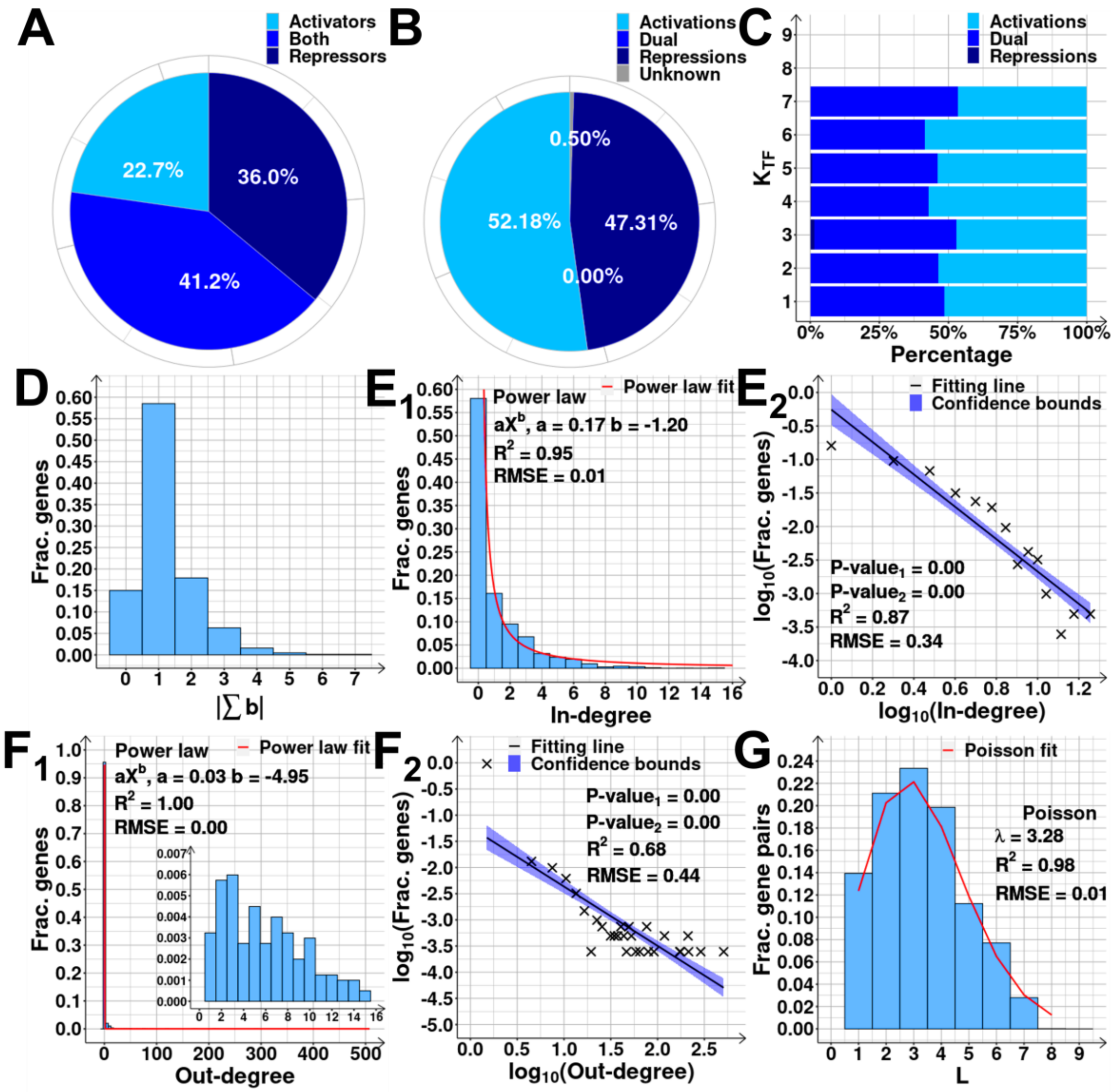

**Figure S12. Topology and logic of the transcription factor (TF) network, TFN, of *E. coli*.**

**(A)** Pie chart of the percentage of input TFs classified as 'Activators' (activate all genes they regulate) and 'Repressors' (repress all genes they regulate). 'Both' are the rare input TFs with dual effects (i.e., activate some genes and repress other, or have opposite strengths on the same promoter, depending on the conditions).

**(B)** Pie chart of the percentage of input TFs binding sites (interactions) classified as activations, repressions, unknown, or dual (rare cases where the input TF can have a positive or negative effect on the same promoter, depending on other factors).

**(C)** Same as (B), but for each cohort of genes defined by the genes' $K_{TF}$ (number of input TFs).

**(D)** Distribution of the fraction of genes with a given absolute sum of the regulatory effects, $|b|$, of the input TFs. Each input TF binding site is set to have a regulatory effect, '$r$', equal to +1 (activation), -1 (repression), or 0 (dual or unknown effect).

**($E_1$)** In-degree (number of incoming edges) distribution.

**($E_2$)** Scatter plot between the $\log_{10}$ of the In-degree and the $\log_{10}$ of the fraction of genes with corresponding In-degree.

**($F_1$)** Out-degree (number of outgoing edges) distributions.

**($F_2$)** Scatter plot between the $\log_{10}$ of the Out-degree and the $\log_{10}$ of the fraction of genes with corresponding out-degree.

**(G)** Path-length (L) distribution. The path length is the number of edges/input TFs needed to reach one node from another. A Poisson fitting (solid red) is shown with its fitting parameters, coefficient of variation ($R^2$) and root mean square error (RMSE).

For ($E_1$) and ($F_1$), we show power-law fittings (solid red lines) and their fitting parameters, $R^2$ and RMSE. In ($E_2$) and ($F_2$), the best fitting line (solid black), its 68% confidence bounds (blue shadow area), statistics ($R^2$, RMSE, and P-values at 0.1 significance level) were obtained as described in Methods section *Statistical tests c*.

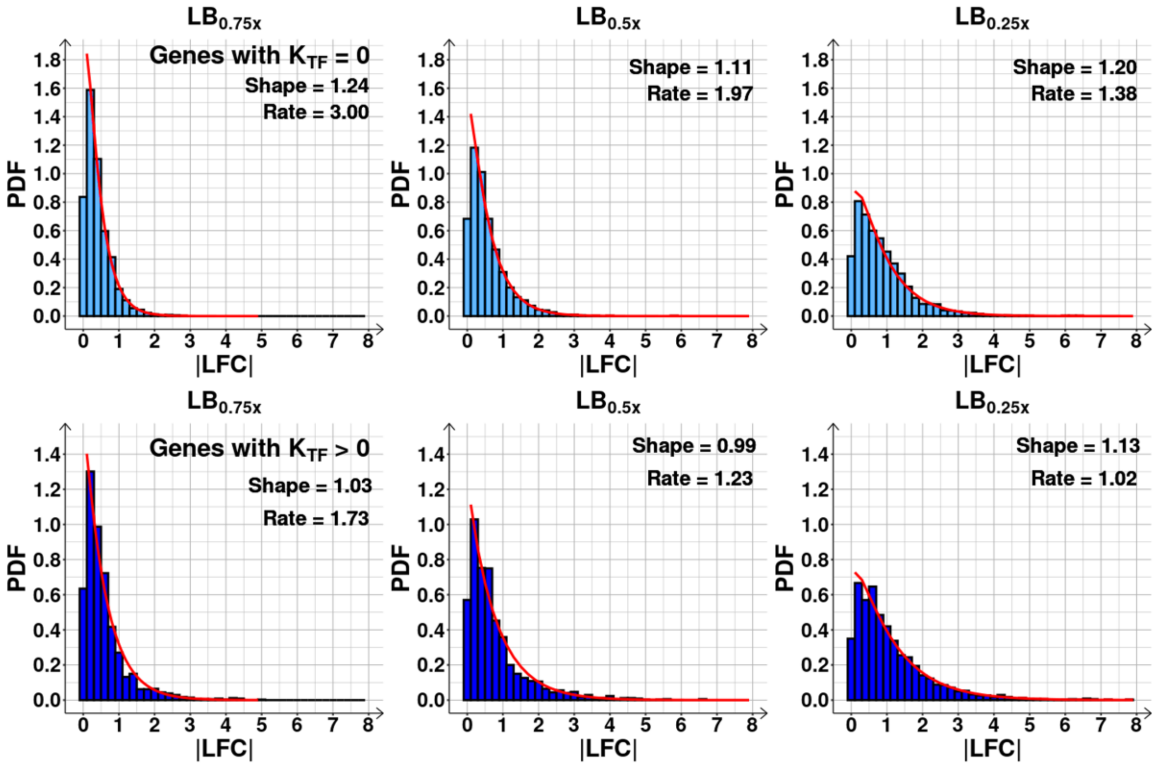

**Figure S13. (Related to Figures 4A) Probability Density Function (PDF) of the number of genes with a given |LFC| (absolute log2 fold change) for each shift**. PDFs obtained from genes without (light blue), and with input transcription factors (TFs) (medium blue). $K_{TF}$ stands for number of input TFs. For each distribution, we fitted a gamma ($\Gamma$) function using *GAMFIT* of MATLAB which tunes the 'Shape' and 'Rate'. We also performed 2-sample T-tests and 2-sample K-tests of statistical significance between the distributions with and without input TFs. For both tests, all p-values were < 0.1.

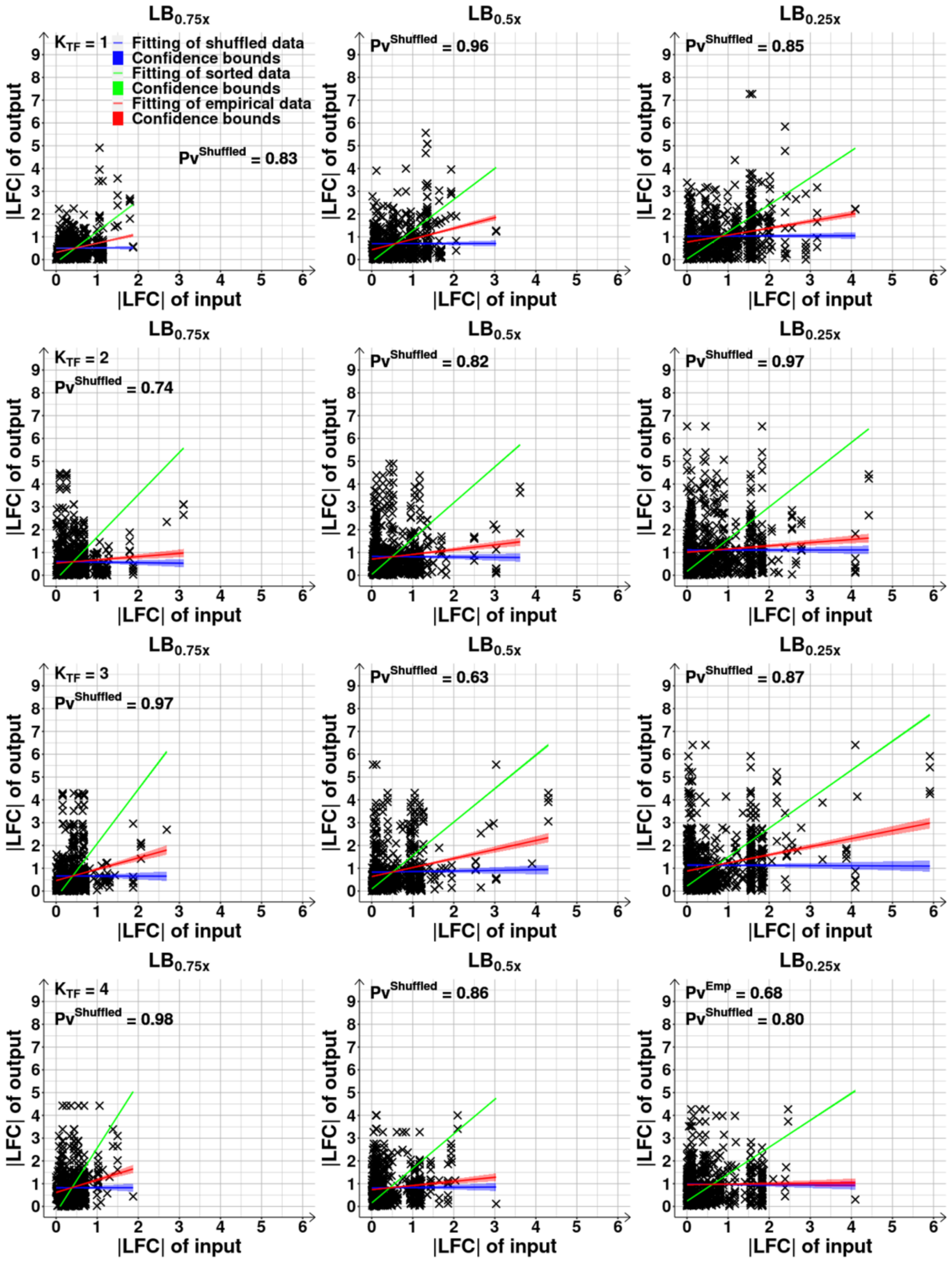

LB0.75x
LB0.5x
LB0.25x
KTF = 1
Fitting of shuffled data
Confidence bounds
Fitting of sorted data
Confidence bounds
Fitting of empirical data
Confidence bounds
PvShuffled = 0.83
PvShuffled = 0.96
PvShuffled = 0.85
KTF = 2
PvShuffled = 0.74
PvShuffled = 0.82
PvShuffled = 0.97
KTF = 3
PvShuffled = 0.97
PvShuffled = 0.63
PvShuffled = 0.87
KTF = 4
PvShuffled = 0.98
PvShuffled = 0.86
PvEmp = 0.68
PvShuffled = 0.80
|LFC| of output
|LFC| of input

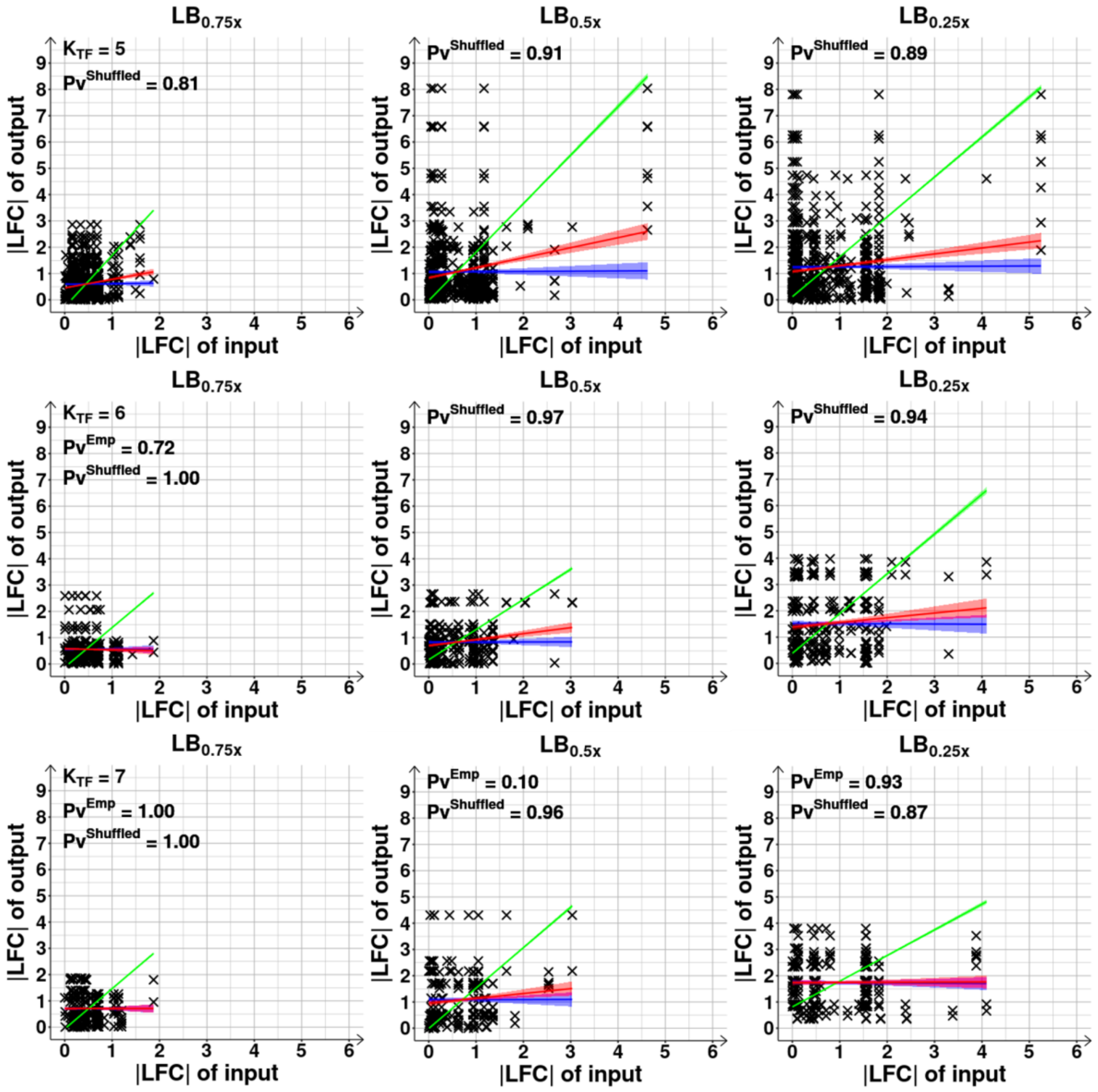

**Figure S14. (Related to Figure 4B and Supplementary Figure S20B) Changes in RNA abundances of input genes plotted against those of the output genes, as a function of $K_{TF}$, the number of input transcription factors (TFs) of the output gene**. Scatter plots of |LFC| (absolute log2 of fold change) of each output gene with the |LFC| of each gene expressing their direct input TFs (i.e., input genes), following the shifts from the control ($LB_{1.0x}$) to $LB_{0.75x}$, $LB_{0.5x}$ and $LB_{0.25x}$, for each class of genes defined by their $K_{TF}$ (from 1 to 7). All genes are included, regardless of being differentially expressed. The red line is the best fitting one. The blue line is a null-model fitting line and was obtained as described in Methods section *Statistical tests c*. The green line is the best fitting one after sorting the values of the pairs input-output in ascending order, to estimate the maximum correlation possible between the two variables. Best fitting lines were obtained by linear least-squares regression fit using the MATLAB function *FITLM*. The colored shadow areas represent the 68% confidence bounds of the best fitted lines. We obtained the p-values of statistical significance for the red ($Pv^{Emp}$), blue ($Pv^{Shuffled}$) and green ($Pv^{Sorted}$) lines under the null hypothesis that the data is best fit by a horizontal line. We show the p-values when the null hypothesis was not rejected at 0.1 significance level. p-values and coefficients of determination ($R^2$) for the red ($R^2_{Emp}$) and green ($R^2_{Sorted}$) fitted lines are shown in Supplementary Table S7.

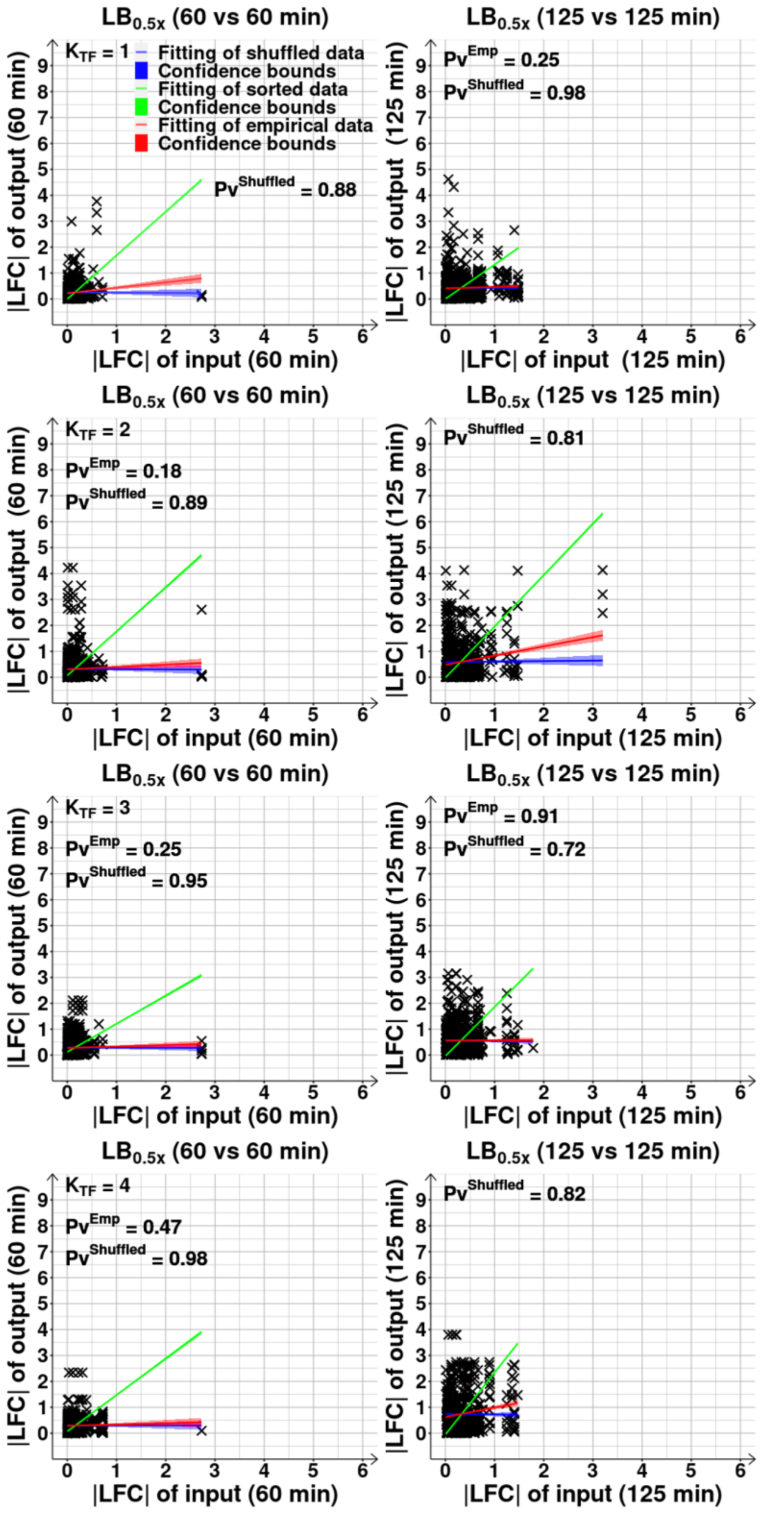

LB0.5x (60 vs 60 min)
LB0.5x (125 vs 125 min)
KTF = 1
Fitting of shuffled data
Confidence bounds
Fitting of sorted data
Confidence bounds
Fitting of empirical data
Confidence bounds
PvShuffled = 0.88
PvEmp = 0.25
PvShuffled = 0.98
KTF = 2
PvEmp = 0.18
PvShuffled = 0.89
PvShuffled = 0.81
KTF = 3
PvEmp = 0.25
PvShuffled = 0.95
PvEmp = 0.91
PvShuffled = 0.72
KTF = 4
PvEmp = 0.47
PvShuffled = 0.98
PvShuffled = 0.82
|LFC| of output (60 min)
|LFC| of output (125 min)
|LFC| of input (60 min)
|LFC| of input (125 min)

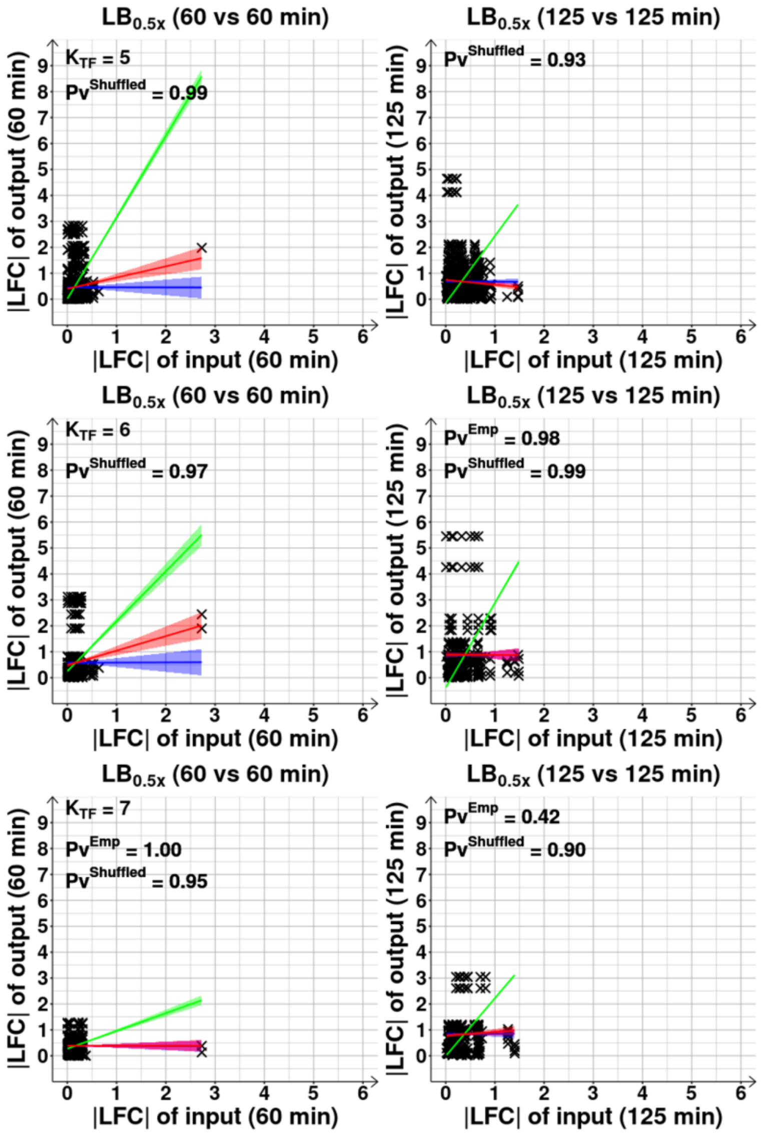

**Figure S15. (Related to Figure 4B) Changes in RNA abundances (prior to the RNAP changes and during the short-term) of input genes plotted against their output genes, as a function of $K_{TF}$, the number of input transcription factors (TFs) of the output gene**. Scatter plots of |LFC| (absolute log2 of fold change) of each output gene with the |LFC| of each gene expressing their direct input TFs (i.e., input genes), following the medium dilution to $LB_{0.5x}$, for each class of genes defined by their $K_{TF}$ (from 1 to 7). The data regards the short-term responses (125 min) and the responses prior to the changes in RNAP concentration (60 min). All genes are included, regardless of being differentially expressed. The red line is the best fitting one. The blue line is the null-model fitting line and was obtained as described in Methods section *Statistical tests c*. The green line is the best fitting one after sorting the pairs input-output in ascending order, to estimate the maximum correlation possible between the two. Best fitting lines were obtained by linear least-squares regression fit using the MATLAB function *FITLM*. The colored shadow areas represent the 68% confidence bounds of the best fitted lines. We obtained the p-values of statistical significance for the red ($Pv^{Emp}$), blue ($Pv^{Shuffled}$) and green ($Pv^{Sorted}$) lines under the null hypothesis that the data is best fit by a horizontal line. We show the p-values when the null hypothesis was not rejected at 0.1 significance level. p-values and coefficients of determination ($R^2$) for the red ($R^2_{Emp}$) and green ($R^2_{Sorted}$) fitted lines are shown in Supplementary Table S8.

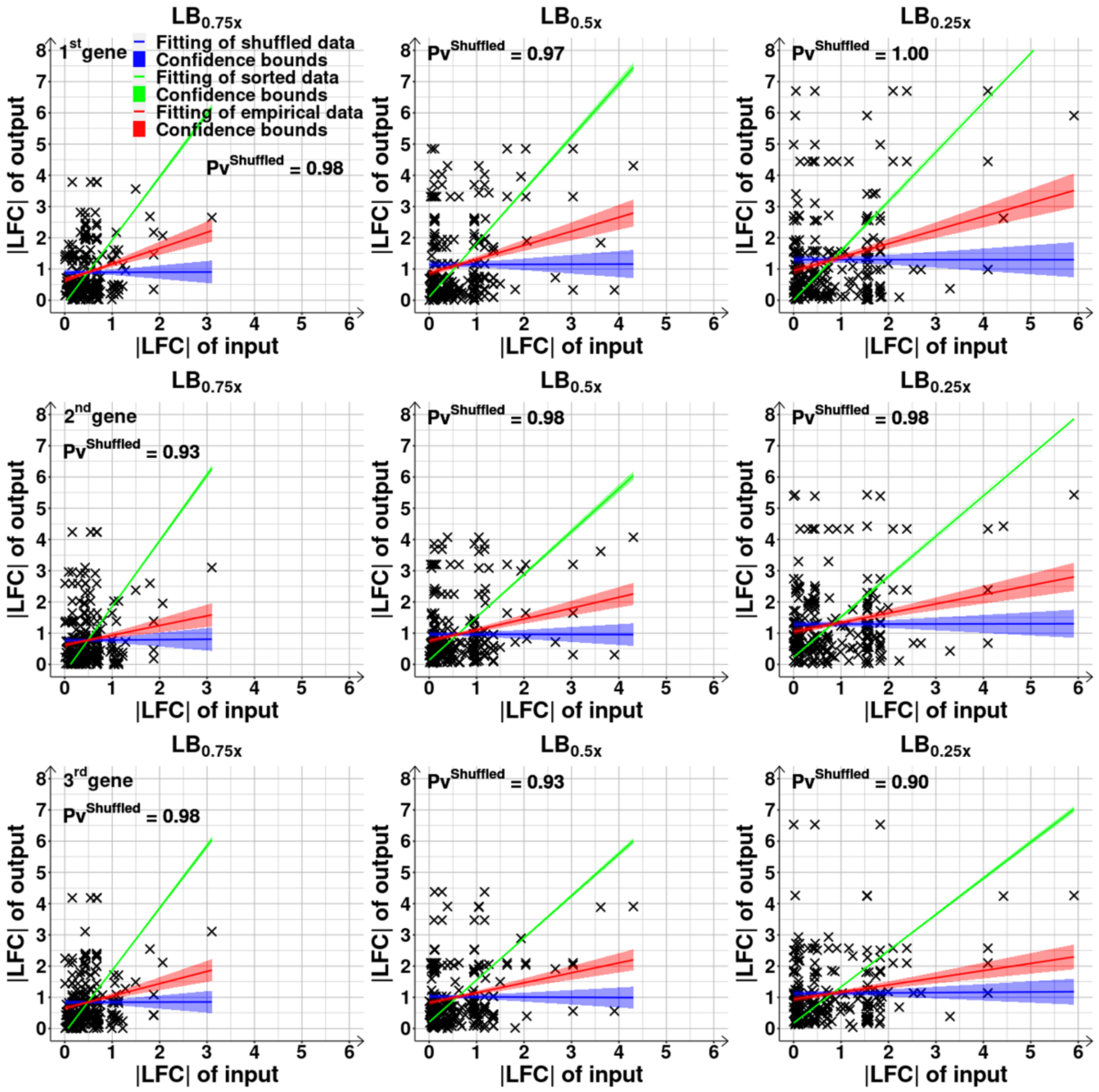

**Figure S16. (Related to Figure 4B) Relationships between the changes in RNA abundances of input and output genes as a function of the position of the output gene in the operon.** Scatter plots between the |LFC| (absolute log2 of fold change) of a gene expressing an input TF (i.e., input gene) and each of its direct output genes belonging to an operon of size 3 on the 1$^{st}$, 2$^{nd}$ and 3$^{rd}$ positions in the operon following the transcription start site. The red line is the best fitting line between the two variables. The blue line is the null-model fitting line and was obtained as described in Methods section *Statistical tests c*. The green line is the best fitting line obtained after sorting the values of the pairs input-output in ascending order to obtain the maximum correlation possible. All best fitting lines were obtained by linear least-squares regression fit using the MATLAB function *FITLM*. The colored shadow areas are the 68% confidence bounds for the best fitted lines. We also obtained the p-values of statistical significance for the red ($Pv^{Emp}$), blue

($Pv^{Shuffled}$) and green ($Pv^{Sorted}$) lines under the null hypothesis that the data is best fit by a horizontal line. We only show the p-values when the null hypothesis was not rejected at 0.1 significance level. All p-values and the coefficients of determination ($R^2$) for the red ($R^2_{Emp}$) and green ($R^2_{Sorted}$) fitted lines are shown in Supplementary Table S9 (see also Supplementary Table S10).

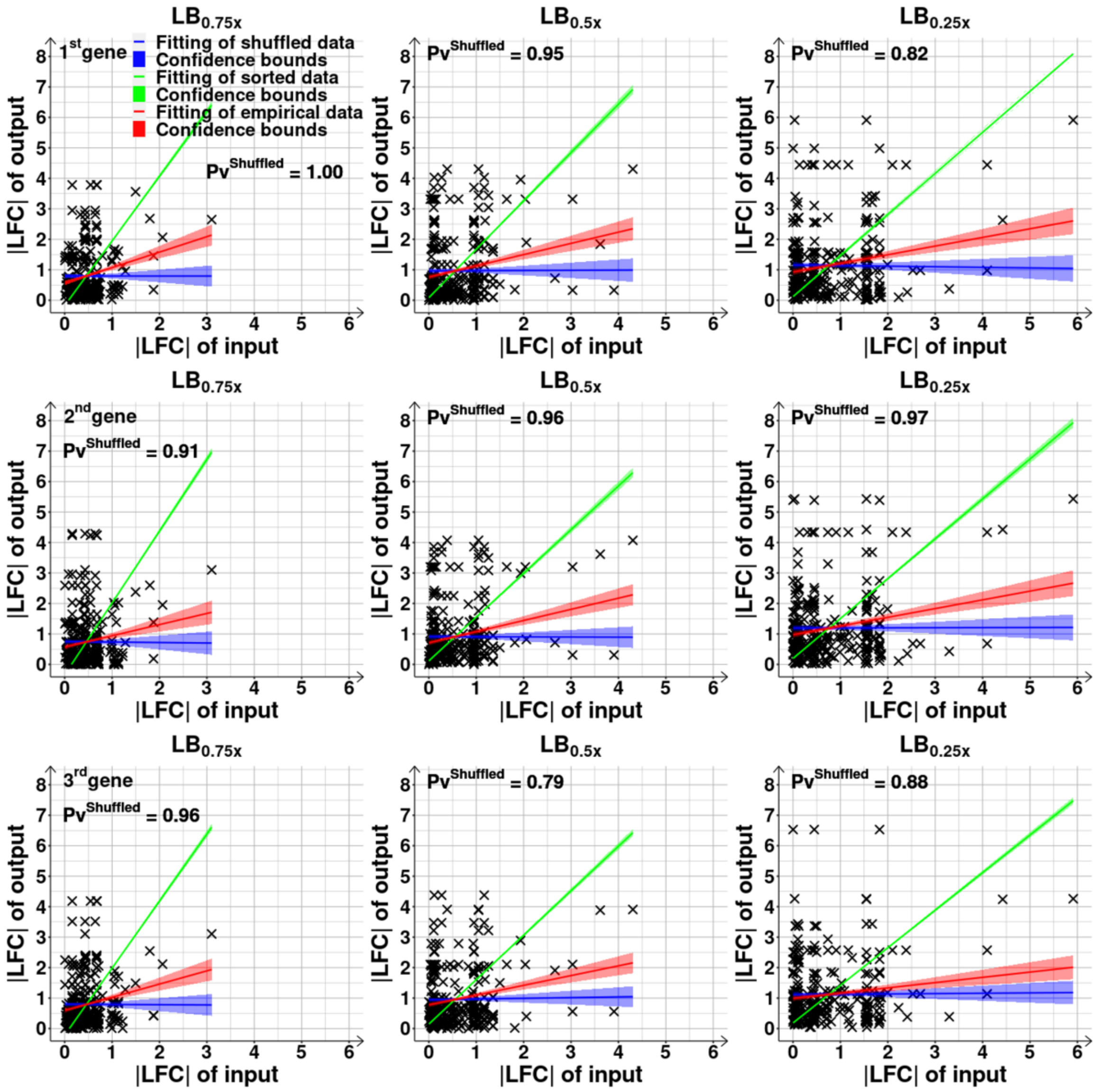

**Figure S17. (Related to Figure 4B) Relationships between the changes in RNA abundances of input and output genes as a function of the position of the output gene in the Transcription Unit (TU).** For each shift to LB$_{0.75x}$, LB$_{0.5x}$ and LB$_{0.25x}$, we plotted the absolute of log2 of fold change of each output gene (|LFC| of output) on the 1$^{st}$, 2$^{nd}$ and 3$^{rd}$ positions in a TU following the transcription start site, against the |LFC| of each gene known to express a direct input transcription factor (|LFC| of input) of these 3 TU genes. Red lines are the best fitting lines. Blue lines are the null-model fitting lines and were obtained as described in Methods section *Statistical tests c*. Green lines are the best fitting after sorting the pairs input-output in ascending order to obtain the maximum correlation possible. All best fitting lines were obtained by linear least-squares regression fit using the MATLAB function *FITLM*. The colored shadow areas represent the 68% confidence bounds for the best fitted lines. We also obtained the p-values of statistical significance for the red ($Pv^{Emp}$), blue ($Pv^{Shuffled}$) and green ($Pv^{Sorted}$) lines under the null hypothesis that the data is best fit by a horizontal line. We only show the p-values when the null hypothesis was not rejected at 0.1 significance level. Supplementary Table S11 shows all p-values and coefficients of determination ($R^2$) of the red ($R^2_{Emp}$) and green ($R^2_{Sorted}$) fitted lines (see also Supplementary Table S12).

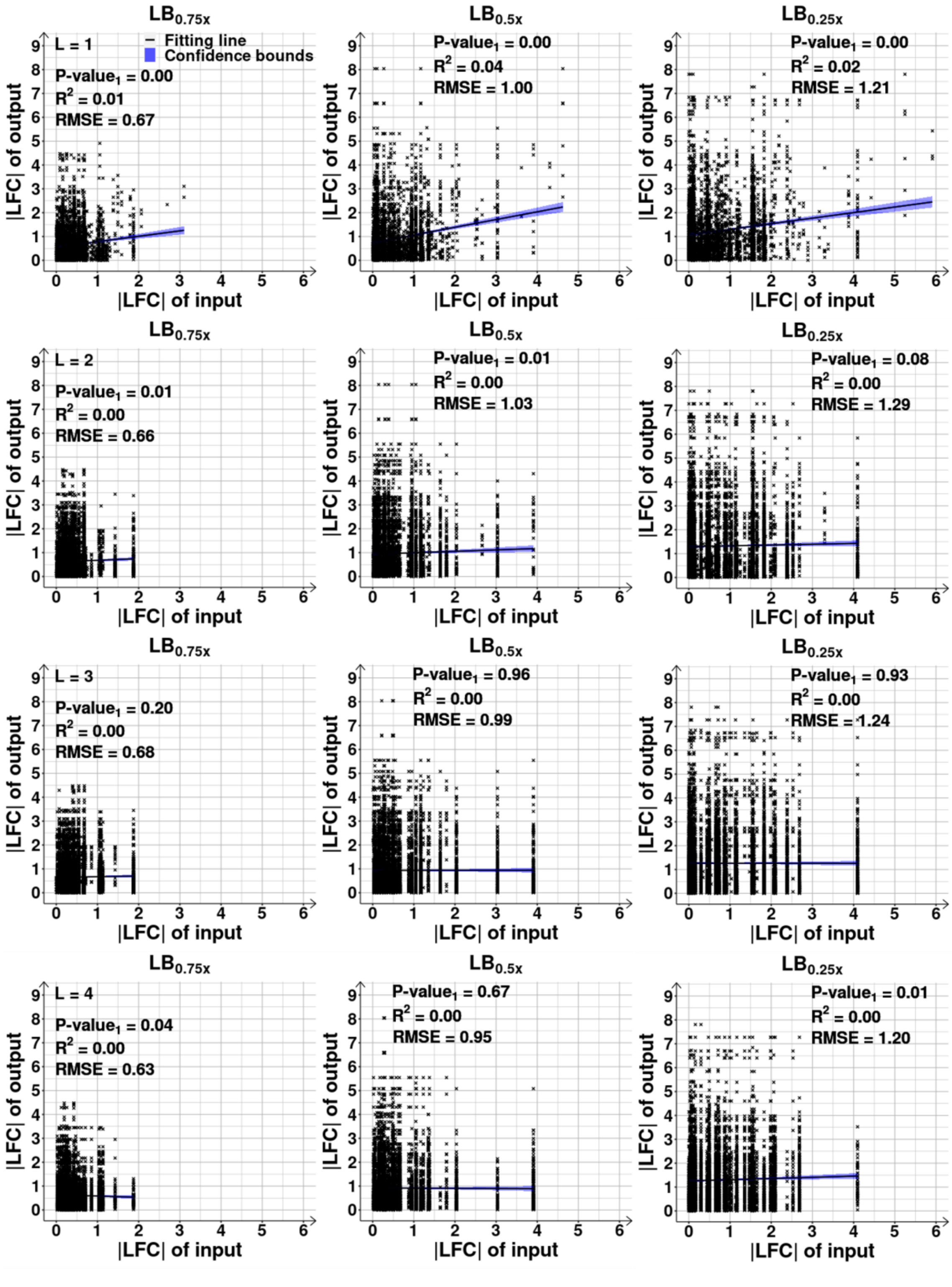
LB0.75x
L = 1
Fitting line
Confidence bounds
P-value1 = 0.00
R2 = 0.01
RMSE = 0.67
LB0.5x
P-value1 = 0.00
R2 = 0.04
RMSE = 1.00
LB0.25x
P-value1 = 0.00
R2 = 0.02
RMSE = 1.21
LB0.75x
L = 2
P-value1 = 0.01
R2 = 0.00
RMSE = 0.66
LB0.5x
P-value1 = 0.01
R2 = 0.00
RMSE = 1.03
LB0.25x
P-value1 = 0.08
R2 = 0.00
RMSE = 1.29
LB0.75x
L = 3
P-value1 = 0.20
R2 = 0.00
RMSE = 0.68
LB0.5x
P-value1 = 0.96
R2 = 0.00
RMSE = 0.99
LB0.25x
P-value1 = 0.93
R2 = 0.00
RMSE = 1.24
LB0.75x
L = 4
P-value1 = 0.04
R2 = 0.00
RMSE = 0.63
LB0.5x
P-value1 = 0.67
R2 = 0.00
RMSE = 0.95
LB0.25x
P-value1 = 0.01
R2 = 0.00
RMSE = 1.20
|LFC| of output
|LFC| of input

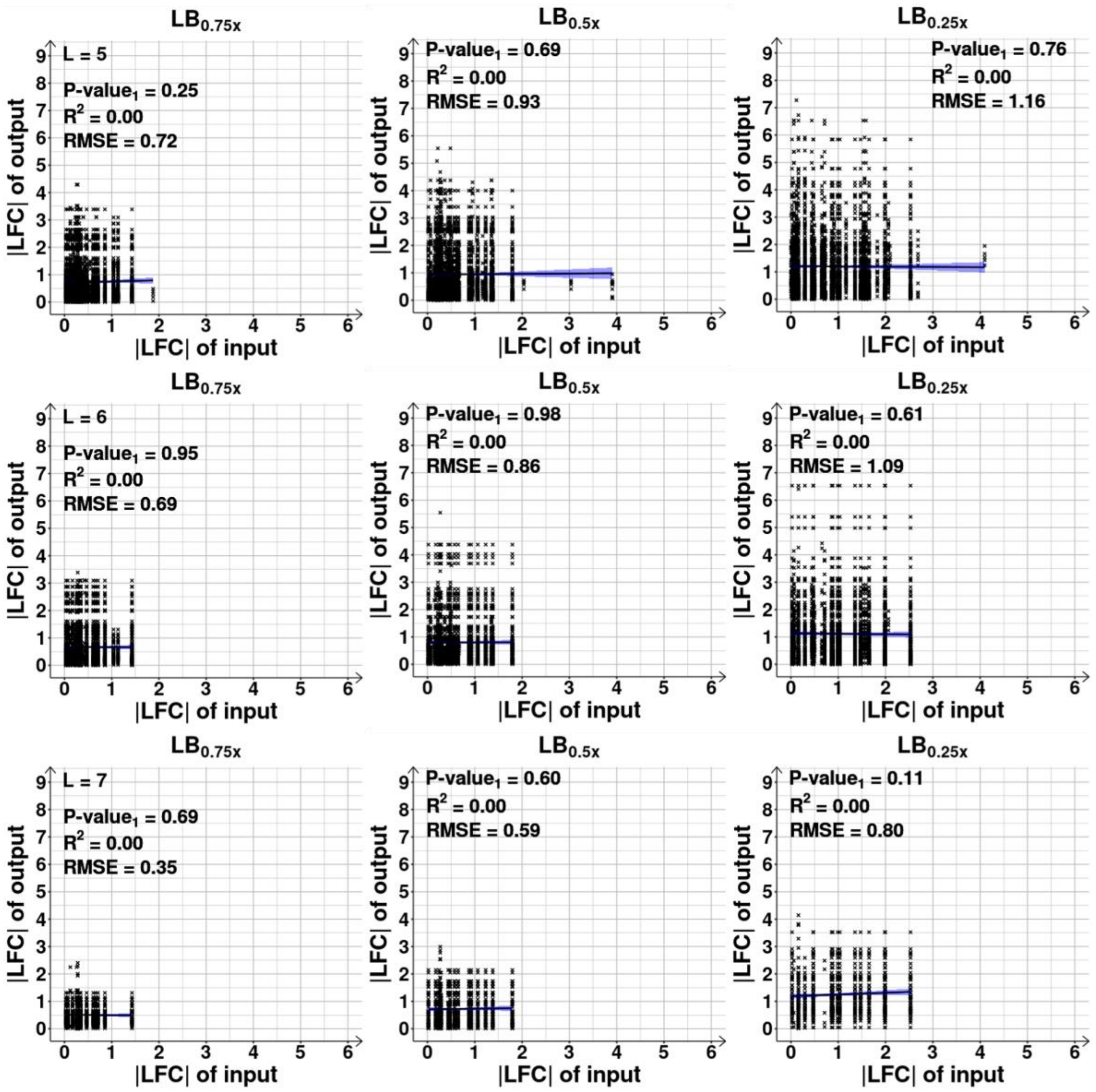
LB0.75x
L = 5
P-value1 = 0.25
R2 = 0.00
RMSE = 0.72
LB0.5x
P-value1 = 0.69
R2 = 0.00
RMSE = 0.93
LB0.25x
P-value1 = 0.76
R2 = 0.00
RMSE = 1.16
LB0.75x
L = 6
P-value1 = 0.95
R2 = 0.00
RMSE = 0.69
LB0.5x
P-value1 = 0.98
R2 = 0.00
RMSE = 0.86
LB0.25x
P-value1 = 0.61
R2 = 0.00
RMSE = 1.09
LB0.75x
L = 7
P-value1 = 0.69
R2 = 0.00
RMSE = 0.35
LB0.5x
P-value1 = 0.60
R2 = 0.00
RMSE = 0.59
LB0.25x
P-value1 = 0.11
R2 = 0.00
RMSE = 0.80
|LFC| of output
|LFC| of input

**Figure S18. (Related to Figure 4C) Relationship between the changes in RNA abundances of output and input genes as a function of their distance in the Transcription Factor (TF) Network, TFN.** Scatter plots between the |LFC| (absolute log2 of fold change) of pairs of genes as a function of their path length (L) (with L=1 to 7, L being the number of edges/input TFs in the TFN to go from one to the other), after shifting from the control condition. We included all gene pairs, regardless of being differentially expressed. The black lines are the best fitting ones (obtained by linear least-squares regression fit, MATLAB function *FITLM*) and the blue shadow areas are their 68% confidence bounds. Also shown are the coefficient of determination ($R^2$) and the root mean square error (RMSE) of the fitted lines, along with their p-values of statistical significance ($P\text{-value}_1$) (at 0.1 significance level, under the null hypothesis that the data is best fit by a horizontal line).

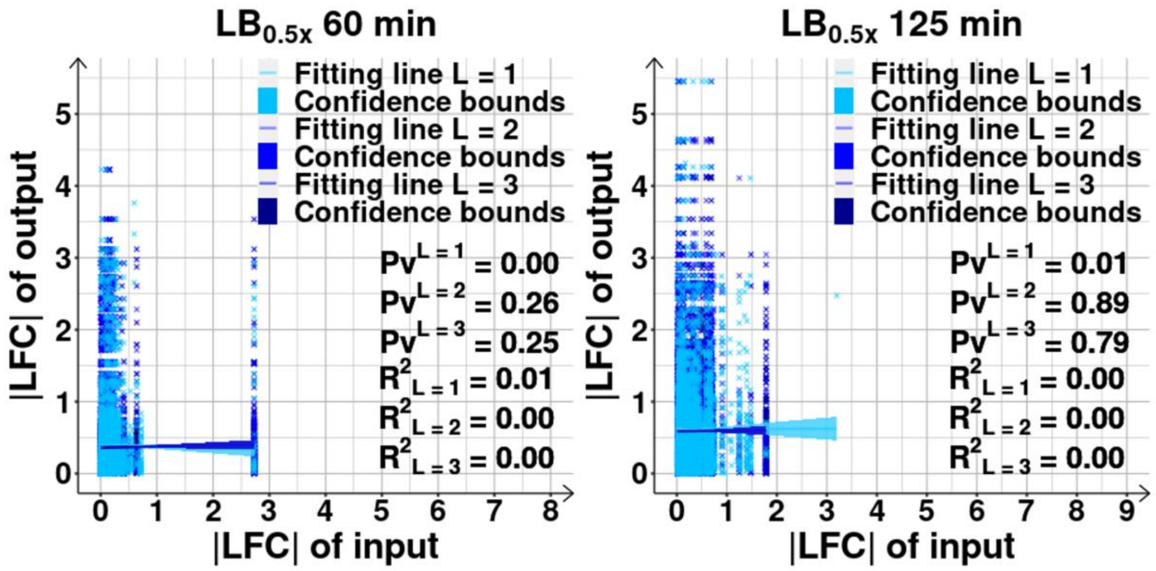

**Figure S19. (Related to Figure 4C) Relationship between the changes in RNA abundances of output and input genes as a function of their path length prior to changes in RNAP and in the short-term following it.** Scatter plots between |LFC| (absolute log2 of fold change) of output and input genes distanced by a minimum path length L of 1, 2, and 3 input TFs (edges) in the TFN, respectively (data from $LB_{0.5x}$). The data regards the short-term responses (125 min) and the responses prior to the changes in RNAP concentration (60 min). We included all gene pairs, regardless of being differentially expressed. The colored lines are the best fitting ones (obtained by linear least-squares regression fit, MATLAB function *FITLM*) and the corresponding blue shadow areas are their 68% confidence bounds. Also shown are the coefficient of determination ($R^2$) of the fitted lines, along with their p-values of statistical significance (Pv) (at 0.1 significance level, under the null hypothesis that the data is best fit by a horizontal line).

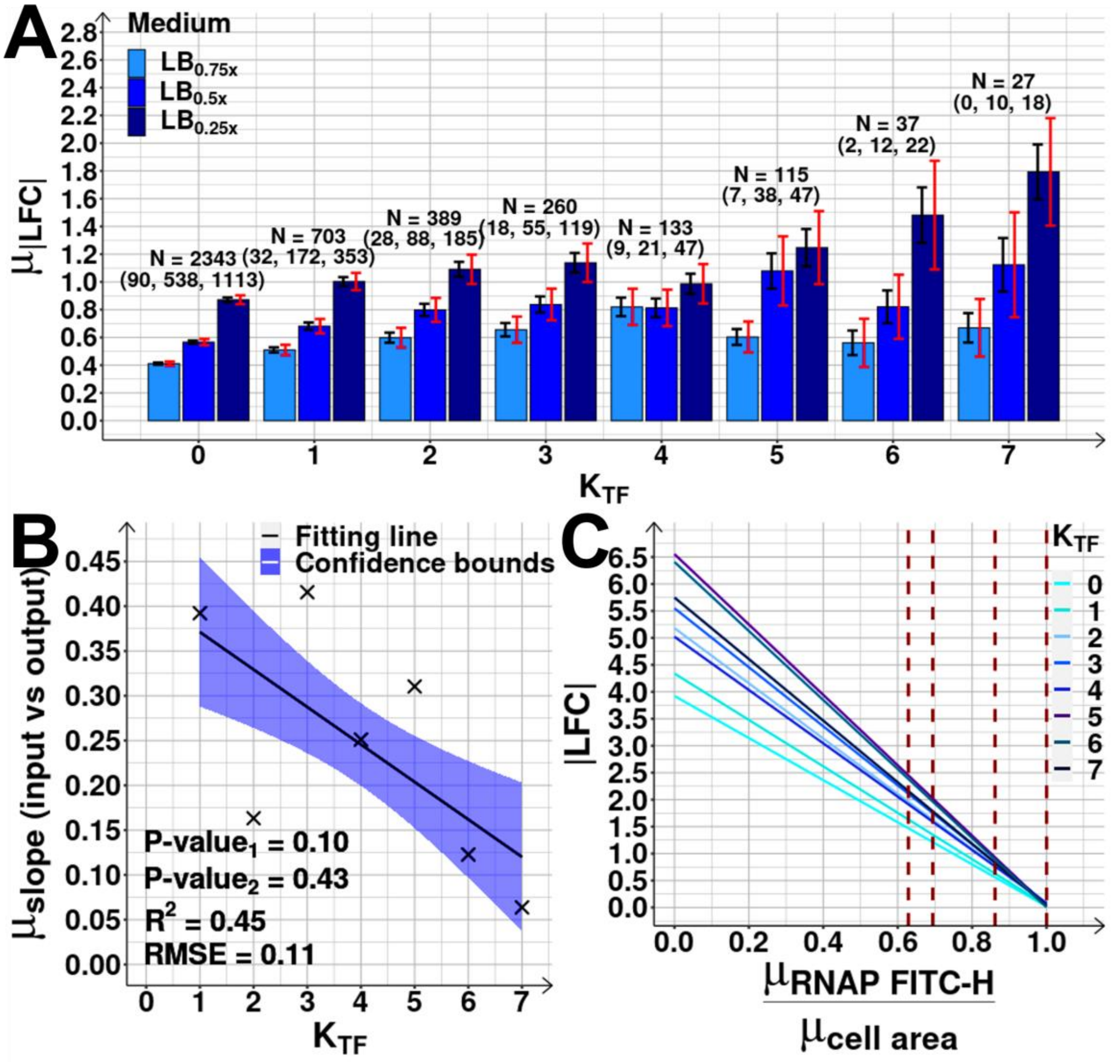

**Figure S20. (Related to Figure 4D) Strengths of the shifts in RNA abundances of cohorts of genes with a given number of input transcription factors (TFs), $K_{TF}$.**

**(A)** Mean of absolute log2 fold changes (|LFC|), $\mu_{|LFC|}$, of gene cohorts organized according to the $K_{TF}$ of the component genes after shifting the medium. Black error bars represent the standard error of the mean (SEM), while red error bars represent the 95% confidence bounds of the SEM. Also shown is the number of genes of each class (*N*), and the numbers of differentially expressed genes (DEG) in each perturbation (Methods section *RNA-seq d*, assuming a False Discovery Rate < 0.05 and a |LFC| > 0.4248 ($LB_{0.75x}$), > 0.4085 ($LB_{0.5x}$) and > 0.4138 ($LB_{0.25x}$)).

**(B)** Scatter plot between $K_{TF}$ and the average slope (over all shifts) of the fitting lines between |LFC| of the output gene and |LFC| of each of its gene expressing a direct input TF (i.e., input gene) (Supplementary Figure S14). The best fitting lines along with their 68% CI and statistics (coefficient of determination ($R^2$), root mean square error (RMSE), and P-values at 0.1

significance level) were obtained as described in Methods section *Statistical tests c*. A slope of 1 is expected if the input fully explains the output.

**(C)** Estimated rate of change of |LFC| with RNAP concentration for gene cohorts differing in $K_{TF}$ (set to 0 in the control). RNAP concentration estimated from the ratio between mean RNAP levels measured by FITC-H, $\mu_{RNAP\ FITC-H}$ (Methods section *Flow-cytometry*), and the mean cell area ($\mu_{cell\ area}$) obtained from phase-contrast images. Only DEG are included. Vertical dashed red lines mark RNAP concentration levels at which RNA-seq was performed. The correlation between the two was obtained by linear least-squares regression fit using *FITLM* of MATLAB. Shown is the best fitting line for each $K_{TF}$. We also obtained p-values of statistical significance of the fitted regression line (at 0.1 significance level, under the null hypothesis that the data is best fit by a horizontal line). For all $K_{TF}$, the p-value < 0.1.

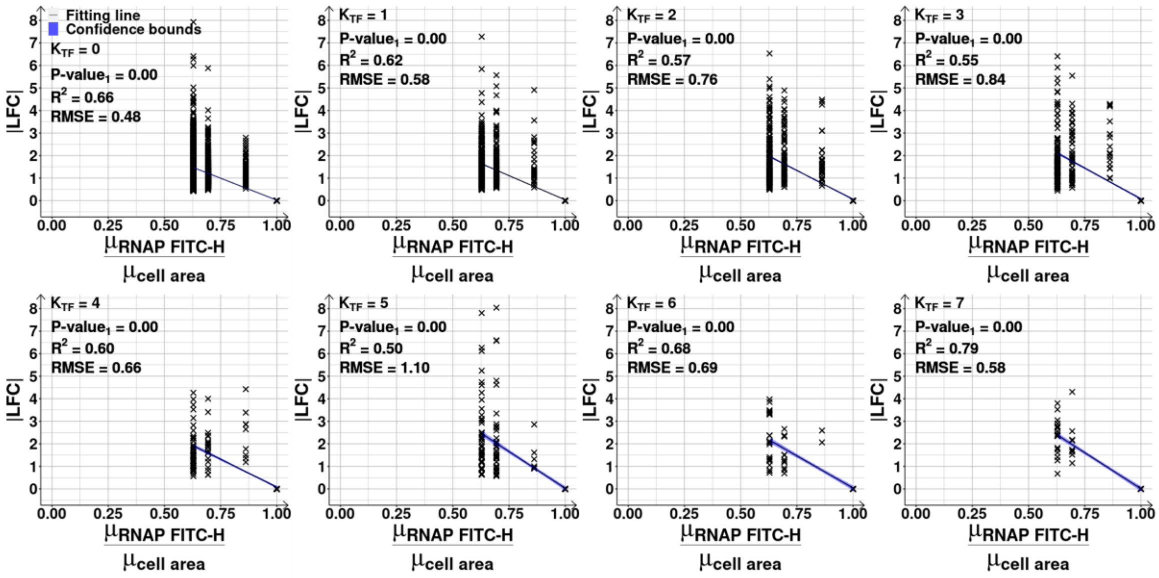

**Figure S21. (Related to Supplementary Figure S20C) RNA changes as a function of the shift in RNAP concentration**. Scatter plots between the |LFC| (absolute log2 of fold change) of each gene following each change in RNAP concentration. This concentration was estimated from the ratio between RNAP levels measured by FITC-H ( $\mu_{RNAP\ FITC-H}$ ) using RL1314 cells (Methods section *Flow-cytometry*), and the mean cell area ( $\mu_{cell\ area}$ ) obtained from phase-contrast images, relative to the control ($LB_{1.0x}$). Data for the cohorts of genes defined by $K_{TF}$ (number of input transcription factors) from 0 to 7. Only differentially expressed genes are included (Methods section *RNA-seq d*, assuming a False Discovery Rate < 0.05 and |LFC| > 0.4248 ($LB_{0.75x}$), > 0.4085 ($LB_{0.5x}$) or > 0.4138 ($LB_{0.25x}$)). Best fitting lines obtained by linear least-squares regression fit using *FITLM* of MATLAB. Blue shadow areas are the 68% confidence bounds. Shown are the coefficient of determination ($R^2$) and the root mean square error (RMSE) of the fitted regression line, along with its p-value of statistical significance (P-value$_1$) (at 0.1 significance level, under the null hypothesis that the data is best fit by a horizontal line). Supplementary Table S13 shows tests of whether the lines differ statistically.

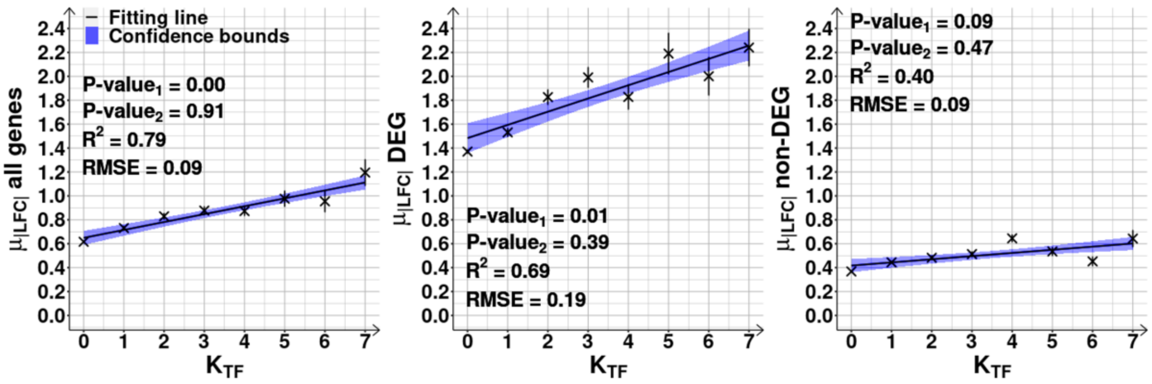

**Figure S22. (Related to Figure 4D) RNA shifts of gene cohorts as a function of the number of input transcription factors (TFs), $K_{TF}$, of the component genes**. Correlation plots between $\mu_{|LFC|}$, the mean of the absolute log2 fold changes (|LFC|), and $K_{TF}$. From left to right, results are shown for all genes, for differentially expressed genes (DEG) and, for non-differentially expressed genes 'non-DEG' (Methods section *RNA-seq d*), assuming a False Discovery Rate < 0.05 and |LFC| > 0.4248 ($LB_{0.75x}$), > 0.4085 ($LB_{0.5x}$) or > 0.4138 ($LB_{0.25x}$). $\mu_{|LFC|}$ obtained from data merged from three conditions ($LB_{0.75x}$, $LB_{0.5x}$ and $LB_{0.25x}$). The best fitting lines (solid black), their 68% confidence bounds (blue shadow area) and statistics (coefficient of determination ($R^2$), root mean square error (RMSE), and P-values at 0.1 significance level) were obtained as described in Methods section *Statistical tests c*. Both vertical and horizontal (not visible) error bars represent the standard errors of the mean.

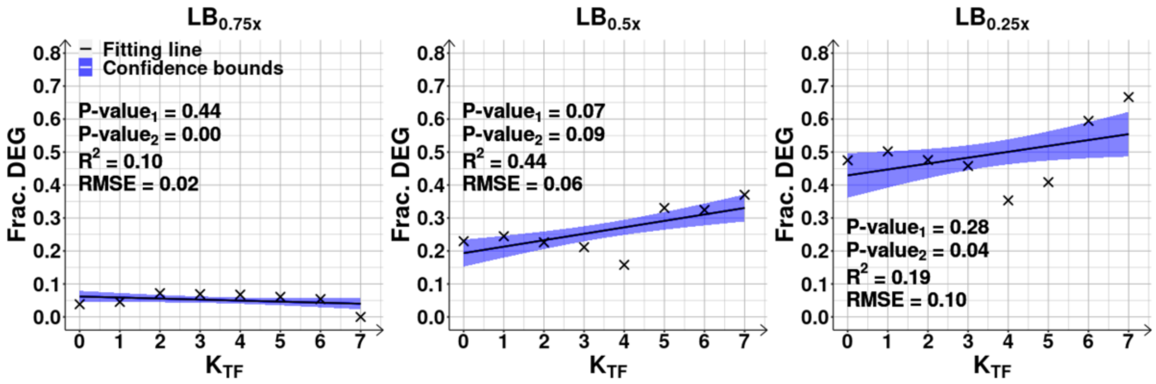

**Figure S23. (Related to Figure 4D and Supplementary Figure S22) Fraction of differentially expressed genes (DEG) as a function of $K_{TF}$, the number of input transcription factors (TFs)**. DEG assessed assuming a False Discovery Rate < 0.05 and absolute log2 fold changes > 0.4248 for $LB_{0.75x}$, > 0.4085 for $LB_{0.5x}$ and > 0.4138 for $LB_{0.25x}$ (Methods section *RNA-seq d*). Data obtained for each shift from the control ($LB_{1.0x}$). Best fitting line (solid black) obtained by linear least-squares regression fit using the MATLAB function *FITLM*. The best fitting lines (solid black) along with their 68% confidence bounds (blue shadow areas) and statistics (coefficient of determination ($R^2$), root mean square error (RMSE), and P-values at 0.1 significance level) were obtained as described in Methods section *Statistical tests c*.

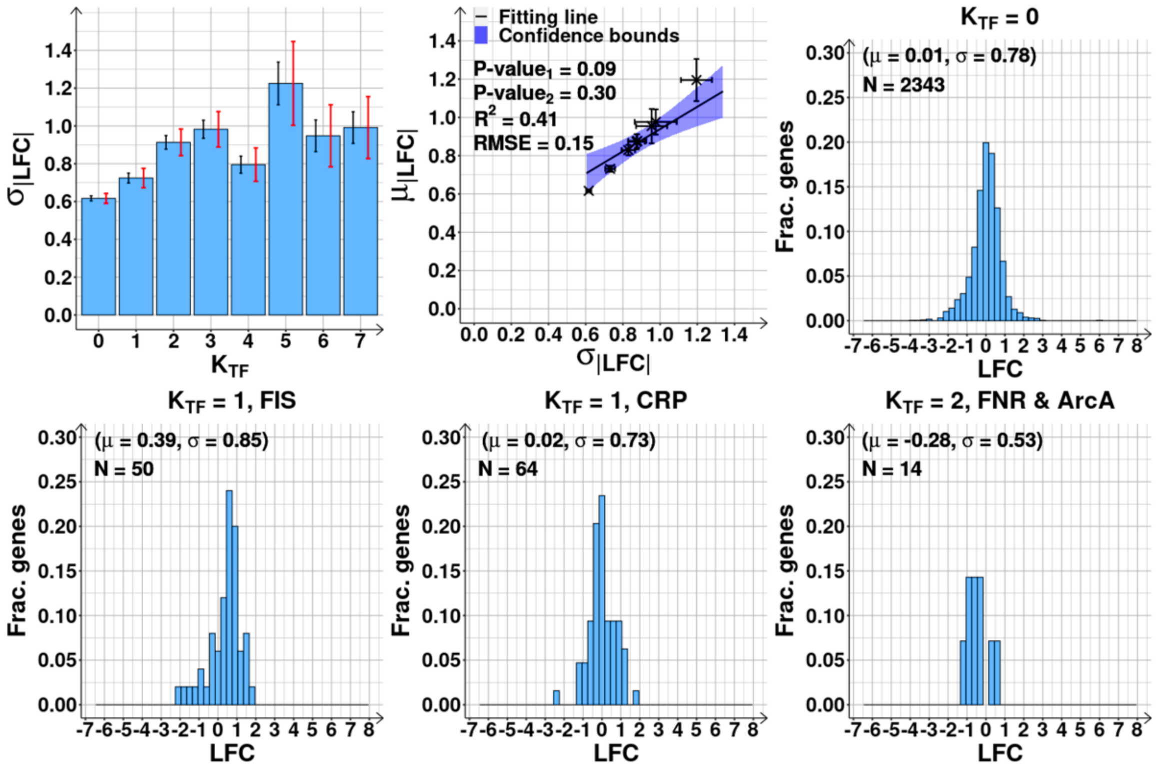

**Figure S24. (Related to Figure 4D) Variability of the strengths of the shifts in RNA abundances of genes cohorts with a given number of input transcription factors (TFs), $K_{TF}$.**

**(A)** Standard deviation of the absolute log2 of fold change (|LFC|), $\sigma_{|LFC|}$, of gene cohorts sharing the same $K_{TF}$, from 0 to 7. Results are from merged data from all shifts ($LB_{0.75x}$, $LB_{0.5x}$ and $LB_{0.25x}$). Black error bars are the standard error of the mean (SEM), while red error bars are the 95% confidence bounds (CB) of the SEM.

**(B)** Scattered plot of $\sigma_{|LFC|}$, following medium shifts, plotted against the respective mean of |LFC| ($\mu_{|LFC|}$). The best fitting line (solid black) along with its 68% CB (blue shadow area) and statistics (coefficient of determination ($R^2$), root mean square error (RMSE), and P-values at 0.1 significance level) were obtained as described in Methods section *Statistical tests c*.

Distribution of the fraction of genes with a given LFC, after shifting from the control ($LB_{1.0x}$) to $LB_{0.5x}$, for 4 gene cohorts: **(C)** Genes without input TFs; **(D)** Genes regulated only by FIS ($K_{TF}$ = 1); and (E) Genes regulated only by CRP and, thus, $K_{TF}$ = 1; **(F)** Genes regulated by FNR and ArcA ($K_{TF}$ = 2).

Shown are the number of genes (N) and the mean (μ) and standard deviation (σ) of the LFC's of each distribution. Supplementary Table S14 shows statistical tests comparing the distributions.

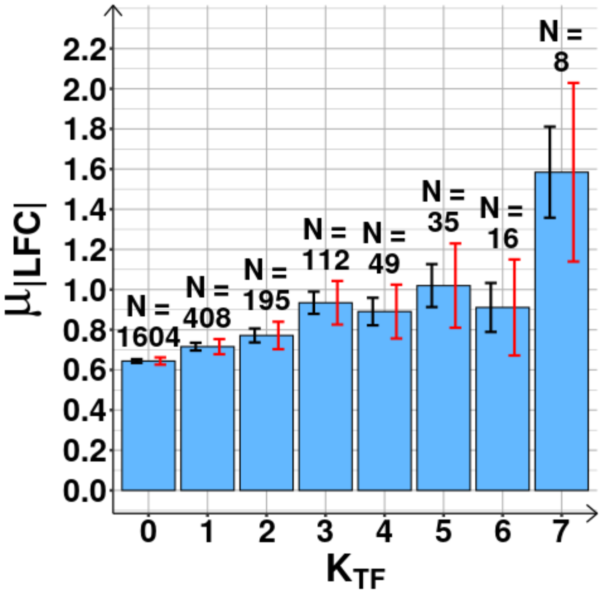

**Figure S25. (Related to Figure 4D) Average RNA fold change of the first gene of each operon (including operons with only 1 gene) as a function of $K_{TF}$, the number of input transcription factors (TFs) of the output gene.** Mean of absolute log2 fold change (|LFC|), $\mu_{|LFC|}$, of all genes in the first position of the operons, following the transcription start site. Results are from merged data from all shifts ($LB_{0.75x}$, $LB_{0.5x}$ and $LB_{0.25x}$). Black error bars are the standard error of the mean (SEM), while red error bars are the 95% confidence bounds of the SEM. *N* stands for the number of genes of each cohort.

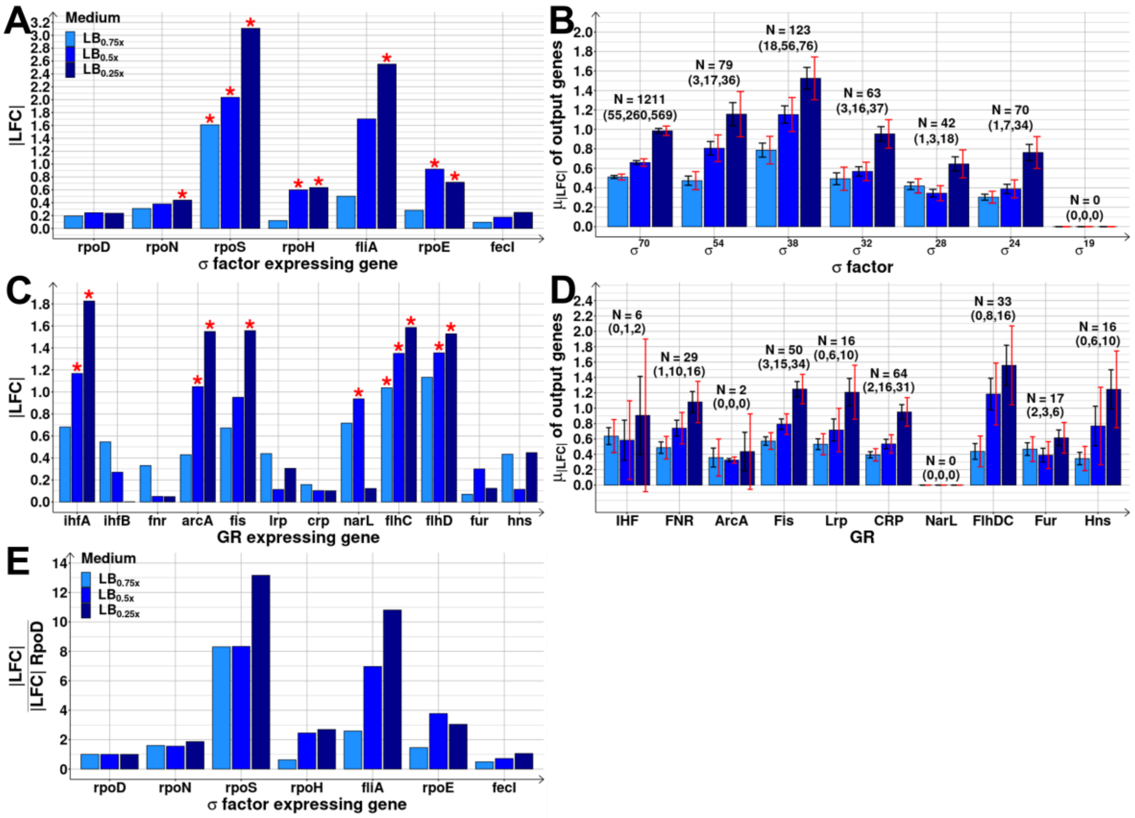

**Figure S26. Shifts in the RNA abundances of global regulators and their output genes.**

**(A)** Absolute of log2 fold change (|LFC|) of the genes expressing each σ factor (rpoD, rpoN, rpoS, rpoH, fliA, rpoE, fecI), following each dilution ($LB_{0.75x}$, $LB_{0.5x}$ and $LB_{0.25x}$).

**(B)** Mean |LFC|, $\mu_{|LFC|}$, of the *N* genes responsive only to $\sigma^{70}$, $\sigma^{54}$, $\sigma^{38}$, $\sigma^{32}$, $\sigma^{28}$, $\sigma^{24}$ and $\sigma^{19}$, respectively.

**(C)** |LFC| of the genes expressing each global regulator (GR) (ihfA, ihfB, fnr, arcA, fis, lrp, crp, narL, flhC, flhD, fur, hns) (10,11).

**(D)** $\mu_{|LFC|}$ of the *N* genes that are responsive only to IHF, FNR, ArcA, Fis, Lrp, CRP, NarL, FlhDC, Fur and Hns, respectively.

In (A) and (C), the red asterisks denote changes that are classified as differentially expressed in Figure 3C, assuming a False Discovery Rate < 0.05 and a |LFC| > 0.4248 ($LB_{0.75x}$), > 0.4085 ($LB_{0.5x}$) or > 0.4138 ($LB_{0.25x}$) (Methods section *RNA-seq d*). In (B) and (D), the black error bars are the standard error of the mean (SEM), while red error bars are the 95% confidence bounds of the SEM.

**(E)** |LFC| of the genes expressing each σ factor (rpoN, rpoS, rpoH, fliA, rpoE, fecI), following each dilution ($LB_{0.75x}$, $LB_{0.5x}$ and $LB_{0.25x}$). Values are relative to the |LFC| of rpoD in the same conditions.

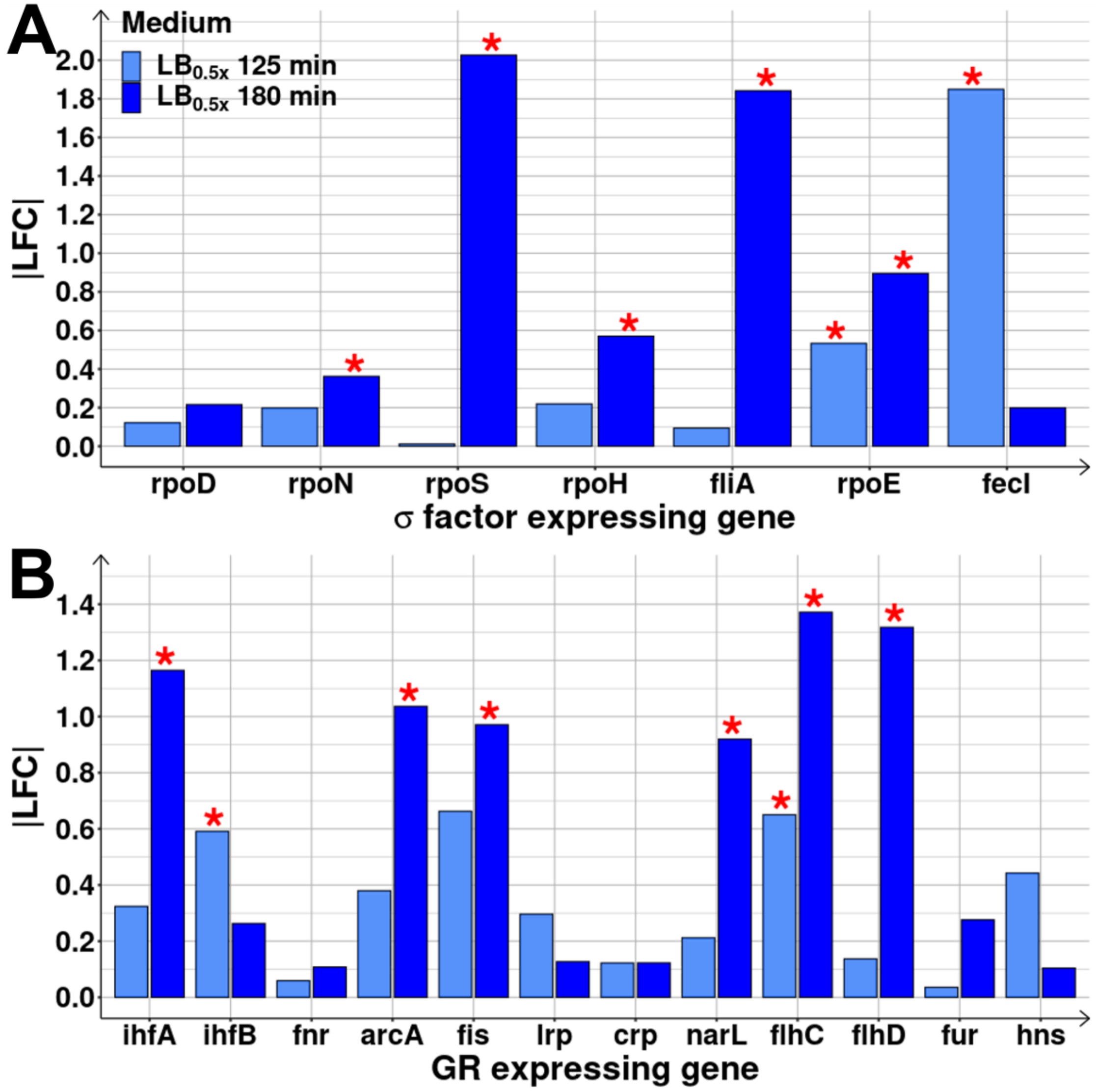

**Figure S27. Shifts in the RNA abundances of global regulators and sigma factors at short- and mid-term response.**

**(A)** Absolute of log2 fold change (|LFC|) of the genes expressing each σ factor (rpoD, rpoN, rpoS, rpoH, fliA, rpoE, fecI), when diluting the control medium ($LB_{1.0x}$) to the $LB_{0.5x}$ medium at 125 (short-term response) and 180 min (mid-term response). Here, the raw count matrices at 125 and 180 min were merged and only genes that passed the filtering were studied (Methods section *RNA-seq*).

**(B)** |LFC| of the genes expressing each global regulator (GR) (ihfA, ihfB, fnr, arcA, fis, lrp, crp, narL, flhC, flhD, fur, hns) under the same conditions as (A).

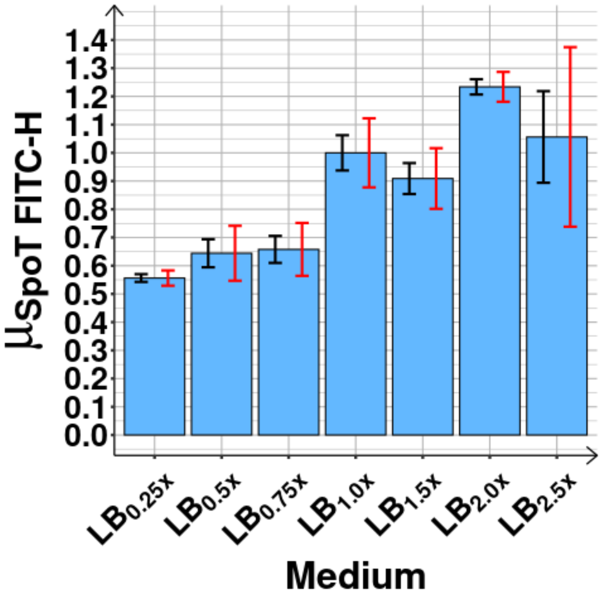

**Figure S28. Expression levels of the spoT gene following medium shifts from $LB_{1.0x}$ to $LB_{0.75x}$, $LB_{0.5x}$, $LB0_{.25x}$, $LB_{1.5x}$, $LB_{2.0x}$ and $LB_{2.5x}$.** Mean SpoT protein levels ( $\mu_{SpoT\ FITC-H}$ ) at 180 min, from 3 biological replicates in each medium condition, measured by the mean single-cell fluorescence intensities (FITC channel, Methods section *Flow-cytometry*), after subtracting mean background fluorescence(s) and scaling to the control $LB_{1.0x}$ condition.

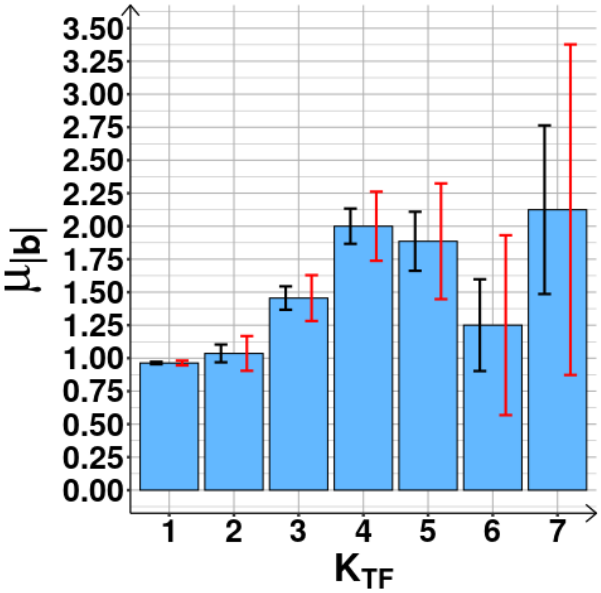

**Figure S29. (Related to Figure 5A). Mean absolute biases of the sets of input transcription factors (TFs) of the first gene of each operon, as a function of the number of input TFs, $K_{TF}$, of the output gene.** Mean of absolute of the sum of the regulatory effects '*r*' of the inputs ($\mu_{|b|}$) obtained from RegulonDB for all genes assessed by RNA-seq. We removed the genes with positions higher than 1 following the transcription start site of the operon. Black error bars are the standard error of the mean (SEM), and red error bars are the 95% confidence bounds of the SEM.

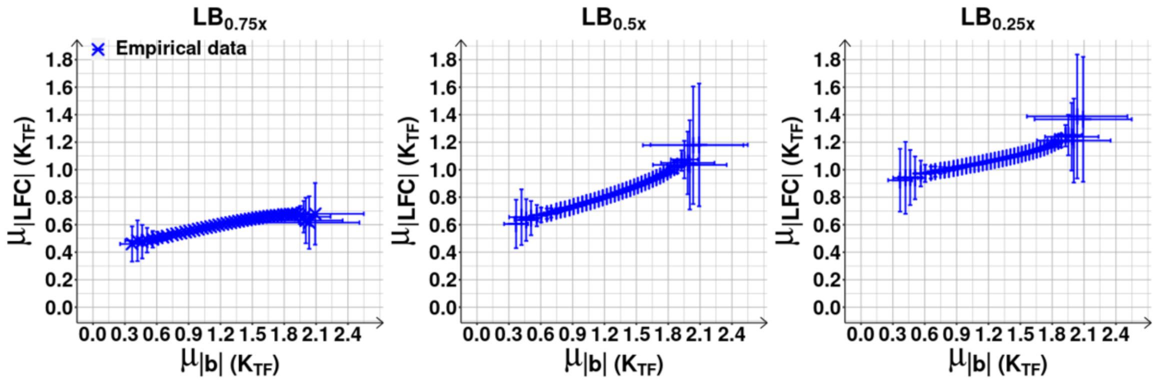

**Figure S30. (Related to Figures 5B) Mean changes in RNA abundances of genes with an average $K_{TF}$, number of input transcription factors (TFs), as a function of $\mu_{|b|}$ (mean absolute bias in the regulatory effects of the input TFs) estimated using the ensemble approach.** Mean of the absolute LFC (log2 of fold change), $\mu_{|LFC|}$, as a function of $\mu_{|b|}\left(K_{TF}\right)$ for each medium shift. Cohorts are assembled based on the $K_{TF}$ of the genes. Each blue cross is the average outcome from up to 7500 cohorts of 10 genes, Supplementary Results section *Estimation of the expected* $\mu_{K_{TF}}$ *and* $\mu_{|b|}$ *using an ensemble approach*, see also Supplementary Figure S31). Error bars (vertical and horizontal) are the standard error of the mean.

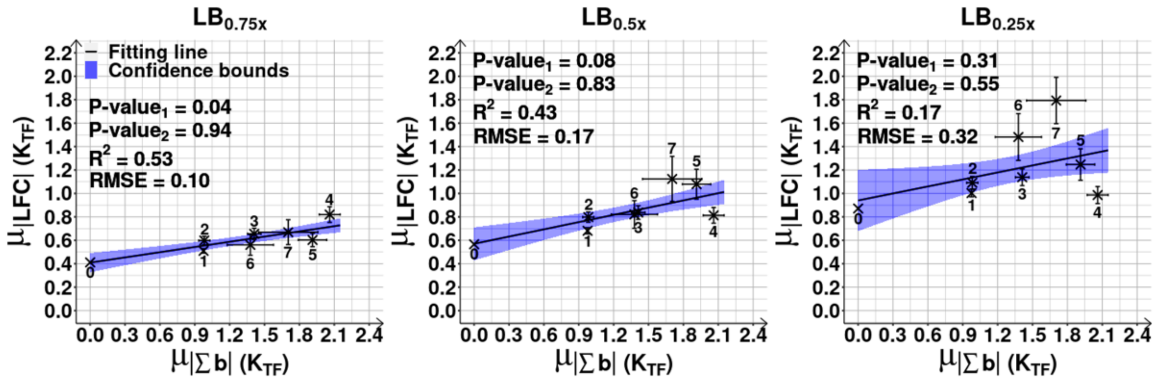

**Figure S31. (Related to Supplementary Figure S30) Mean changes in RNA abundances of all genes with a specific $K_{TF}$, number of input transcription factors (TFs), as a function of the average $|b|$ (absolute of overall regulatory effect ('$r$') of the input TFs on the output gene).** For each shift, we plot the mean of absolute LFC (log2 of fold change), $\mu_{|LFC|}$, against the mean of $|b|$, $\mu_{|b|}$, for each cohort of all genes with $K_{TF}$ = 0 to 7. The error bars represent the standard error of the mean (SEM). We included all genes, regardless of being differentially expressed. The best fitting line (solid black) along with its 68% confidence bounds (blue shadow area) and statistics (coefficient of determination ($R^2$), root mean square error (RMSE), and P-values at 0.1 significance level) were obtained as described in Methods section *Statistical tests c*. Supplementary Table S18 shows the results of the statistical tests.

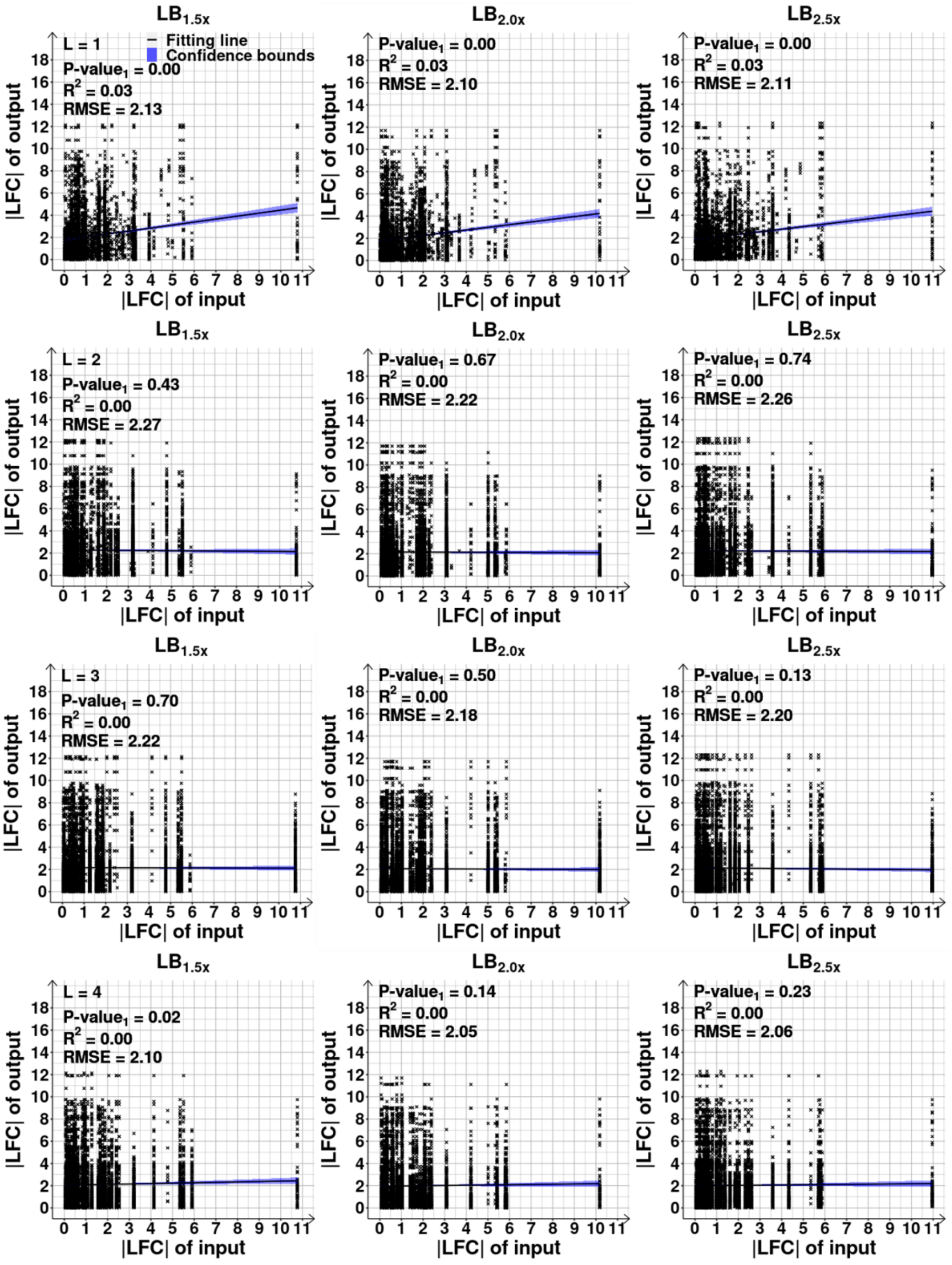
LB1.5x
L = 1
Fitting line
Confidence bounds
P-value1 = 0.00
R2 = 0.03
RMSE = 2.13
|LFC| of output
|LFC| of input
LB2.0x
P-value1 = 0.00
R2 = 0.03
RMSE = 2.10
LB2.5x
P-value1 = 0.00
R2 = 0.03
RMSE = 2.11
LB1.5x
L = 2
P-value1 = 0.43
R2 = 0.00
RMSE = 2.27
LB2.0x
P-value1 = 0.67
R2 = 0.00
RMSE = 2.22
LB2.5x
P-value1 = 0.74
R2 = 0.00
RMSE = 2.26
LB1.5x
L = 3
P-value1 = 0.70
R2 = 0.00
RMSE = 2.22
LB2.0x
P-value1 = 0.50
R2 = 0.00
RMSE = 2.18
LB2.5x
P-value1 = 0.13
R2 = 0.00
RMSE = 2.20
LB1.5x
L = 4
P-value1 = 0.02
R2 = 0.00
RMSE = 2.10
LB2.0x
P-value1 = 0.14
R2 = 0.00
RMSE = 2.05
LB2.5x
P-value1 = 0.23
R2 = 0.00
RMSE = 2.06

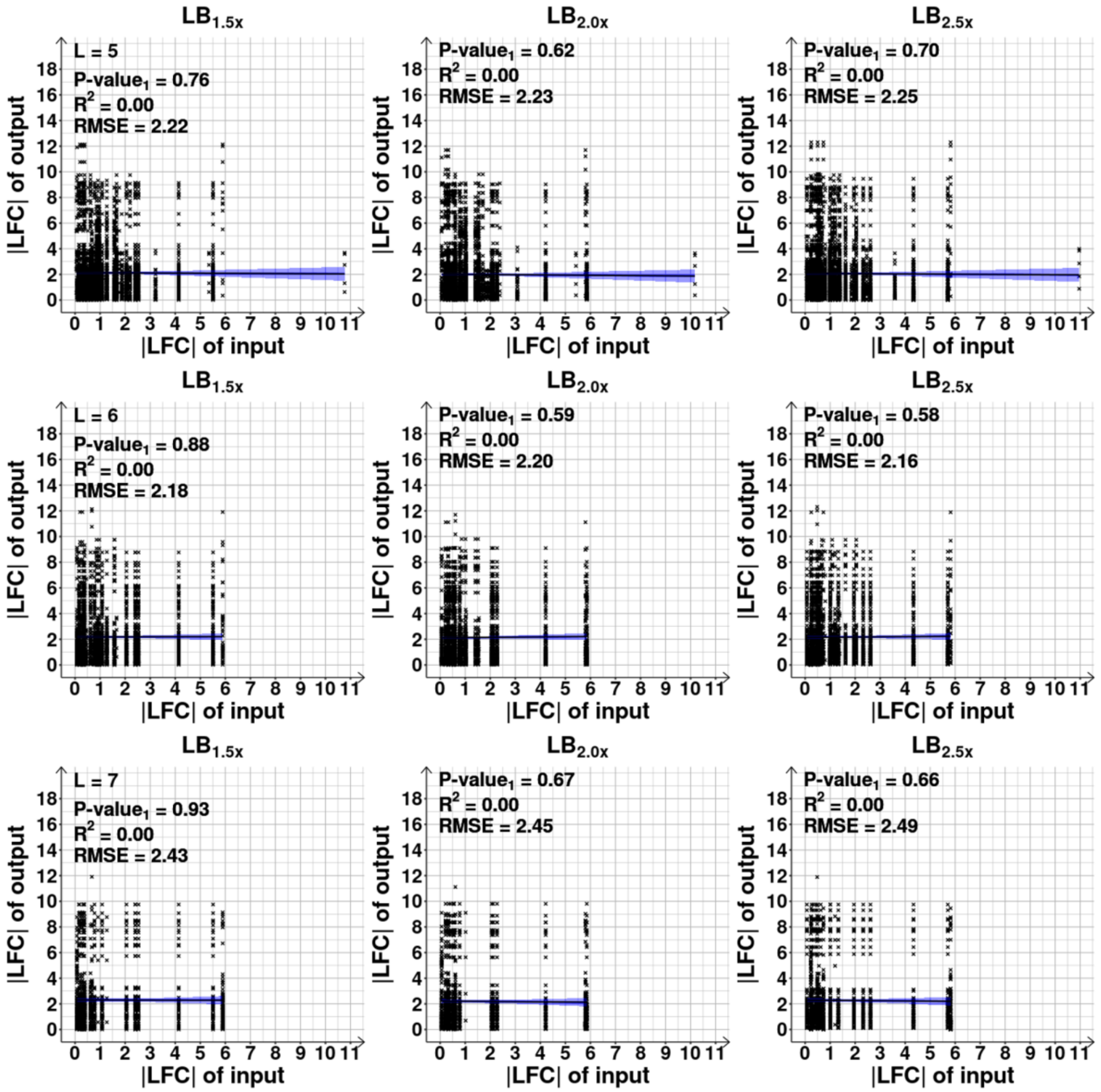

**Figure S32. (Related to Figures $6D_1$-$6D_3$) Changes in RNA abundances of output and input genes plotted as a function of their distance in the Transcription Factor (TF) Network, TFN.** Scatter plots between the |LFC| (absolute log2 of fold change) of pairs of genes as a function of their path length (L) (with L=1 to 7, and L being the number of edges/input TFs in the TFN to go from one gene to the other), after shifting from the control condition. We included all gene pairs, regardless of being differentially expressed or not. Black lines are the best fitting ones (obtained by linear least-squares regression fit, MATLAB function *FITLM*) and blue shadow areas are their 68% confidence bounds. Also shown are the coefficient of determination ($R^2$), the root mean square error (RMSE) of the fitted lines, and their p-values of statistical significance (P-value$_1$) at 0.1 significance level, under the null hypothesis that the data is best fit by a horizontal line.

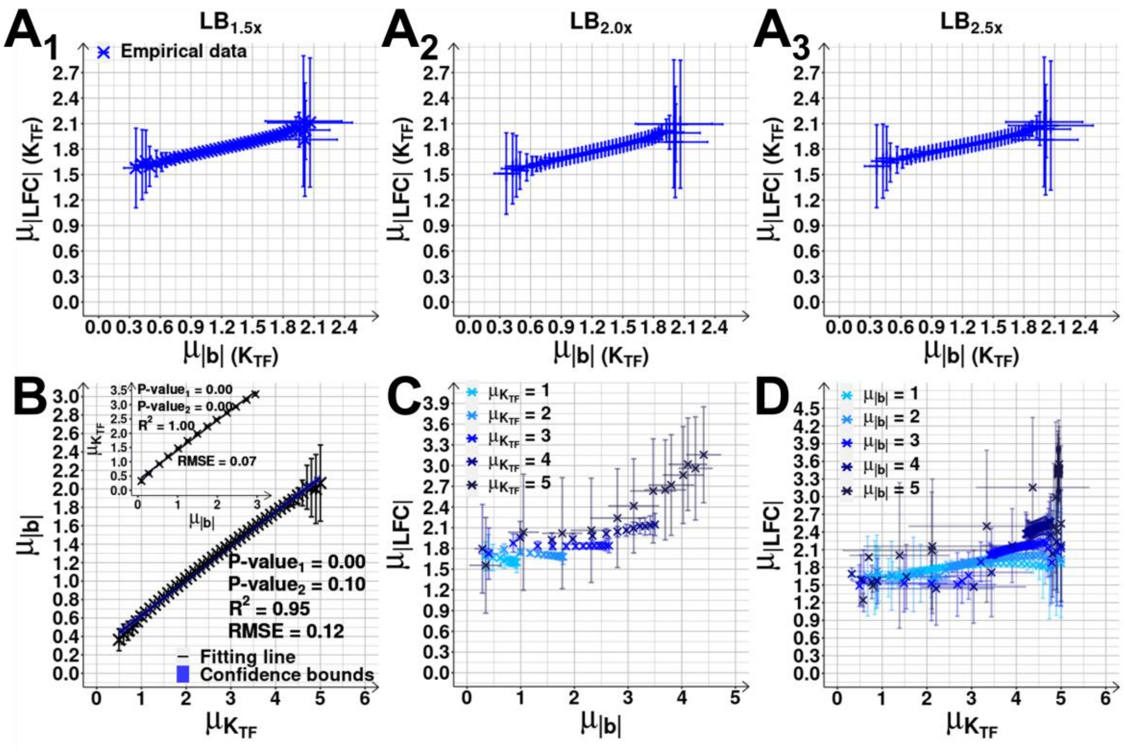

**Figure S33. (Related to Figure 7D) Mean changes in RNA abundances of cohorts with an average $K_{TF}$ or $|b|$ (mean number of input TFs and mean absolute bias, respectively).**

**($A_1$-$A_3$)** Mean |LFC|, $\mu_{|LFC|}$, as a function of $\mu_{|b|}(K_{TF})$. Data obtained using the ensemble approach (Supplementary Results section *Estimation of the expected* $\mu_{K_{TF}}$ *and* $\mu_{|b|}$ *using an ensemble approach*). Each blue cross is the average outcome from up to 7.500 cohorts of 10 genes.

**(B)** Scatter plot of $\mu_{K_{TF}}$ against the corresponding $\mu_{|b|}$ of the cohorts in ($A_1$-A3). The inset shows the inverse, scatter plot between $\mu_{|b|}$ and $\mu_{K_{TF}}$, for the cohorts of Figure 7D, assembled based on $\mu_{|b|}$ (Supplementary Results section *Estimation of the expected* $\mu_{K_{TF}}$ *and* $\mu_{|b|}$ *using an ensemble approach*). Shown are best fitting lines and their 68% confidence bound (shadow areas, barely visible), coefficient of determination ($R^2$), root mean square error (RMSE), and P-value (Methods section *Statistical tests c*).

**(C)** Cohorts with increasing $\mu_{K_{TF}}$ but constant $\mu_{|b|}$ (from 1 to 5). $\mu_{K_{TF}}$ is plotted against the corresponding $\mu_{|LFC|}$, for each $\mu_{|b|}$.

**(D)** $\mu_{|b|}$ plotted against the corresponding $\mu_{|LFC|}$ for cohorts with constant $\mu_{K_{TF}}$ (from 1 to 5) and increasing $\mu_{|b|}$ (Supplementary Results section *Estimation of the expected* $\mu_{K_{TF}}$ *and* $\mu_{|b|}$ *using an ensemble approach*).

The error bars (vertical and horizontal) are the standard error of the mean. In (C) and (D), the data from the different conditions was merged and, since they slightly differ in mean ($A_1$-$A_3$), the SEM is larger than if in each condition separately. Also, comparatively, the SEM is much larger for $\mu_{K_{TF}}$ and $\mu_{|b|}$ equal to 5, due to which we did not extend the analysis further.

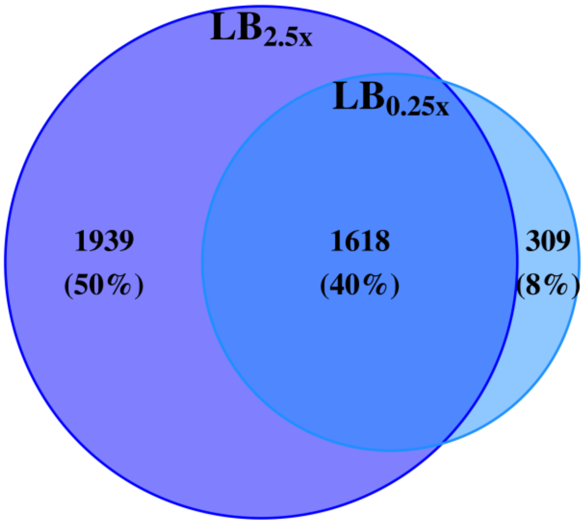

**Figure S34. Venn diagram of the number and percentage of differentially expressed genes following RNAP changes**. Data from 180 min following the shift to $LB_{0.25x}$ in the light blue circle and data from the shift to $LB_{2.5x}$ in the dark violet circle, when overlapping, become dark blue.

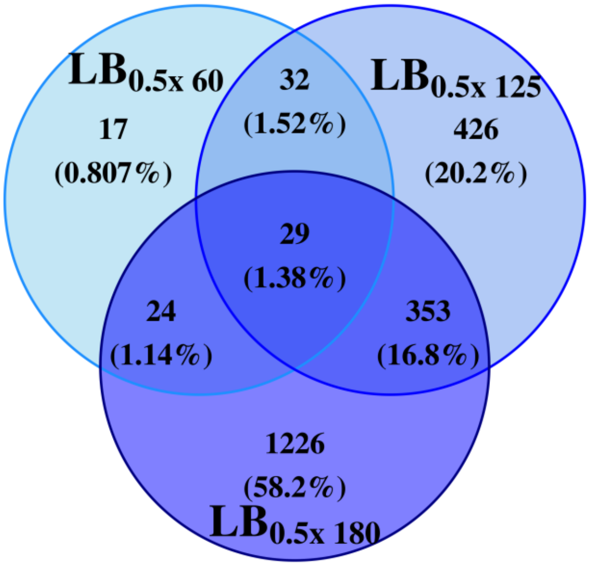

**Figure S35. Venn diagram of the number and percentage of differentially expressed genes**. Data from the shift to $LB_{0.5x}$ at 60 (prior to the changes in RNAP), 125 (short-term response) and 180 min (mid-term response).

**SUPPLEMENTARY TABLES**

**Table S1. Variables.** Short description of the main variables used.

| Variable | Description |
|---|---|
| $K_{TF}$ | Number of input transcription factors (TFs) of a gene. |
| $\mu_{K_{TF}}$ | Mean of $K_{TF}$ of a gene cohort. |
| $r$ | Regulatory effect (+1, 0, -1) |
| $b = \left\|\sum r\right\|$ | Bias. It equals the absolute of the sum of the regulatory effects, '$r$' (each equaling +1 or -1) of the input TFs of a gene. |
| $\mu_{\|b\|}$ | Mean absolute $b$ of a gene cohort |
| $\mu_{\|LFC\|}$ | Mean of \|LFC\| (absolute of log2 fold changes). |
| **TFN** | Transcription factor network |
| **GR** | Global regulator |
| **L** | Path length |
| **CC** | Closed complex formation |
| **OC** | Open complex formation |
| **SEM** | Standard error of the mean |
| **CB** | Confidence bounds |

**Table S2. (Related to Figure 2E and Supplementary Figure S2) Raw data of RNAP levels by western blotting.** Shown are all the measurement values, for the three biological replicates, as reported by the software 'ImageLab' after analyzing the images obtained by Western Blot (Supplementary Figure S2).

| Replicate 1 | | | | | | | | |
|---|---|---|---|---|---|---|---|---|
| **Lane** | **Sample** | **Channel** | **Band No.** | **Relative Front** | **Volume (Int)** | **Band %** | **Norm. Factor** | **Norm. Vol. (Int)** |
| **2** | $LB_{1.0x}$ | Chemi | 1 | 0.344164 | 21470300 | 100 | 1.000000 | 21470300 |
| **3** | $LB_{0.75x}$ | Chemi | 1 | 0.341757 | 16063000 | 100 | 0.916722 | 14725306 |
| **4** | $LB_{0.5x}$ | Chemi | 1 | 0.327316 | 9521061 | 100 | 1.132143 | 10779204 |
| **5** | $LB_{0.25x}$ | Chemi | 1 | 0.321300 | 3072728 | 100 | 1.798396 | 5525982 |
| **Replicate 2** | | | | | | | | |
| **Lane** | **Sample** | **Channel** | **Band No.** | **Relative Front** | **Volume (Int)** | **Band %** | **Norm. Factor** | **Norm. Vol. (Int)** |
| **2** | $LB_{1.0x}$ | Chemi | 1 | 0.168000 | 19907978 | 100 | 1.000000 | 19907978 |
| **3** | $LB_{0.75x}$ | Chemi | 1 | 0.162667 | 19135360 | 100 | 0.814673 | 15589056 |
| **4** | $LB_{0.5x}$ | Chemi | 1 | 0.161333 | 8066324 | 100 | 1.511187 | 12189726 |
| **5** | $LB_{0.25x}$ | Chemi | 1 | 0.152000 | 2059213 | 100 | 2.638734 | 5433715 |
| **Replicate 3** | | | | | | | | |
| **Lane** | **Sample** | **Channel** | **Band No.** | **Relative Front** | **Volume (Int)** | **Band %** | **Norm. Factor** | **Norm. Vol. (Int)** |
| **2** | $LB_{1.0x}$ | Chemi | 1 | 0.213828 | 28023120 | 100 | 1.000000 | 28023120 |
| **3** | $LB_{0.75x}$ | Chemi | 1 | 0.215109 | 22070880 | 100 | 1.006164 | 22206921 |
| **4** | $LB_{0.5x}$ | Chemi | 1 | 0.212548 | 13065840 | 100 | 1.228384 | 16049866 |
| **5** | $LB_{0.25x}$ | Chemi | 1 | 0.207426 | 4816080 | 100 | 1.782460 | 8584472 |

**Table S3. (Related to Figure 3B and Supplementary Figure S8A-8C) Statistical test between the distributions in Supplementary Figure S8A-8C.** P-values from the 2-sample T-test between the mean of the distributions (Methods section *Statistical tests a*).

| P-value | | | |
|---|---|---|---|
| | **$LB_{0.75x}$** | **$LB_{0.5x}$** | **$LB_{0.25x}$** |
| **$LB_{0.75x}$** | 1.00 | | |
| **$LB_{0.5x}$** | 0.09 | 1.00 | |
| **$LB_{0.25x}$** | 0.95 | 0.24 | 1.00 |

**Table S4. List of strains carrying an integrated YFP gene copy selected from a YFP strain library** (Methods sections *Bacterial strains, media, growth conditions and curves* and *Flow-cytometry*). Cells used to test for correlations between the absolute log2 of fold change in protein levels and RNA abundances (Supplementary Figure S9).

| | Strain [CGSC name] | Genotype | Source |
|---|---|---|---|
| **1** | cbpM [SX1494] | F-, Δ(argF-lac)169, gal-490, Δ(modF-ybhJ)803, λ[cI857 Δ(cro-bioA)], cbpM791-YFP(::cat), IN(rrnD-rrnE)1, rph-1 | Yale CGSC (CGSC # 13049) |
| **2** | tktB [SX1954] | F-, Δ(argF-lac)169, gal-490, Δ(modF-ybhJ)803, λ[cI857 Δ(cro-bioA)], tktB792-YFP(::cat), IN(rrnD-rrnE)1, rph-1 | Yale CGSC (CGSC # 13509) |
| **3** | groS [SX1398] | F-, Δ(argF-lac)169, gal-490, Δ(modF-ybhJ)803, λ[cI857 Δ(cro-bioA)], IN(rrnD-rrnE)1, rph-1, groS791-YFP(::cat) | Yale CGSC (CGSC # 12953) |
| **4** | yafD [SX1626] | F-, yafD792-YFP(::cat), Δ(argF-lac)169, gal-490, Δ(modF-ybhJ)803, λ[cI857 Δ(cro-bioA)], IN(rrnD-rrnE)1, rph-1 | Yale CGSC (CGSC # 13181) |
| **5** | bolA [SX1087] | F-, Δ(argF-lac)169, bolA791-YFP(::cat), gal-490, Δ(modF-ybhJ)803, λ[cI857 Δ(cro-bioA)], IN(rrnD-rrnE)1, rph-1 | Yale CGSC (CGSC # 12642) |
| **6** | nudI [SX1271] | F-, Δ(argF-lac)169, gal-490, Δ(modF-ybhJ)803, λ[cI857 Δ(cro-bioA)], nudI792-YFP(::cat), IN(rrnD-rrnE)1, rph-1 | Yale CGSC (CGSC # 12826) |
| **7** | cnu [SX1362] | F-, Δ(argF-lac)169, gal-490, Δ(modF-ybhJ)803, λ[cI857 Δ(cro-bioA)], cnu-791-YFP(::cat), IN(rrnD-rrnE)1, rph-1 | Yale CGSC (CGSC # 12917) |
| **8** | mobA [SX1354] | F-, Δ(argF-lac)169, gal-490, Δ(modF-ybhJ)803, λ[cI857 Δ(cro-bioA)], IN(rrnD-rrnE)1, rph-1, mobA791-YFP(::cat) | Yale CGSC (CGSC # 12909) |
| **9** | cpxR [SX1791] | F-, Δ(argF-lac)169, gal-490, Δ(modF-ybhJ)803, λ[cI857 Δ(cro-bioA)], IN(rrnD-rrnE)1, rph-1, cpxR791-YFP(::cat) | Yale CGSC (CGSC # 13346) |
| **10** | yciU [SX1384] | F-, Δ(argF-lac)169, gal-490, Δ(modF-ybhJ)803, λ[cI857 Δ(cro-bioA)], yciU796- | Yale CGSC (CGSC # 12939) |

| | | | |
|---|---|---|---|
| | | YFP(::cat), IN(rrnD-rrnE)1, rph-1 | |
| **11** | yffL [SX1283] | F-, Δ(argF-lac)169, gal-490, Δ(modF-ybhJ)803, λ[cI857 Δ(cro-bioA)], yffL791-YFP(::cat), IN(rrnD-rrnE)1, rph-1 | Yale CGSC (CGSC # 12838) |
| **12** | recN [SX1220] | F-, Δ(argF-lac)169, gal-490, Δ(modF-ybhJ)803, λ[cI857 Δ(cro-bioA)], recN796-YFP(::cat), IN(rrnD-rrnE)1, rph-1 | Yale CGSC (CGSC # 12775) |
| **13** | yceD [SX1638] | F-, Δ(argF-lac)169, gal-490, Δ(modF-ybhJ)803, λ[cI857 Δ(cro-bioA)], yceD792-YFP(::cat), IN(rrnD-rrnE)1, rph-1 | Yale CGSC (CGSC # 13193) |
| **14** | rpsE [SX1340] | F-, Δ(argF-lac)169, gal-490, Δ(modF-ybhJ)803, λ[cI857 Δ(cro-bioA)], IN(rrnD-rrnE)1, rpsE791-YFP(::cat), rph-1 | Yale CGSC (CGSC # 12895) |
| **15** | mrcA [SX1938] | F-, Δ(argF-lac)169, gal-490, Δ(modF-ybhJ)803, λ[cI857 Δ(cro-bioA)], IN(rrnD-rrnE)1, mrcA791-YFP(::cat), rph-1 | Yale CGSC (CGSC # 13493) |
| **16** | napD [SX1339] | F-, Δ(argF-lac)169, gal-490, Δ(modF-ybhJ)803, λ[cI857 Δ(cro-bioA)], napD791-YFP(::cat), IN(rrnD-rrnE)1, rph-1 | Yale CGSC (CGSC # 12894) |
| **17** | hyuA [SX1664] | F-, Δ(argF-lac)169, gal-490, Δ(modF-ybhJ)803, λ[cI857 Δ(cro-bioA)], hyuA791-YFP(::cat), IN(rrnD-rrnE)1, rph-1 | Yale CGSC (CGSC # 13219) |
| **18** | rbsB [SX1190] | F-, Δ(argF-lac)169, gal-490, Δ(modF-ybhJ)803, λ[cI857 Δ(cro-bioA)], IN(rrnD-rrnE)1, rph-1, rbsB791-YFP(::cat) | Yale CGSC (CGSC # 12745) |
| **19** | speC [SX1952] | F-, Δ(argF-lac)169, gal-490, Δ(modF-ybhJ)803, λ[cI857 Δ(cro-bioA)], speC791-YFP(::cat), IN(rrnD-rrnE)1, rph-1 | Yale CGSC (CGSC # 13507) |
| **20** | yjhP [SX1874] | F-, Δ(argF-lac)169, gal-490, Δ(modF-ybhJ)803, λ[cI857 Δ(cro-bioA)], IN(rrnD-rrnE)1, rph-1, yjhP794-YFP(::cat) | Yale CGSC (CGSC # 13429) |

**Table S5. (Related to Supplementary Figure S12) Features of the network topology.** Network global topology parameters of the known TFN of *E. coli* and of (1000) randomly generated networks with the same number of nodes and edges but with 'Erdös Random' and with 'Scale-free' topology (using a Power-law exponent of -1.19) (12). Data from RegulonDB (13), from the genes in the RNA-seq data (Figure 3). For a description of the parameters, see Methods section *Transcription Factor Network of Escherichia coli*.

| **Parameters** | ***E. coli*** | **Erdös random** | **Scale free** |
|---|---|---|---|
| No. nodes | 4053 | 4053 | 4053 |
| No. edges | 4471 | 4471 | 4471 |
| Clustering coefficient | 0.06 | $(1.79 \pm 1.51) \times 10^{-4}$ | $(3.60 \pm 1.49) \times 10^{-4}$ |
| No. connected components | 2324 | 521.46 ± 16.98 | 563 |
| Avg. path length (L) | 3.28 | 24.06 ± 4.69 | 3.32 |
| No. isolated nodes | 2301 | 446.14 ± 17.25 | 485 |
| No. self-loops | 133 | 1.10 ± 1.02 | 0 |

**Table S6. (Related to Figure 4A) Associations between having input transcription factors (TFs) and being a DEG (differentially expressed gene)**. P-values obtained by a Fisher test (Methods section *Statistical tests b*) to determine if there is an association between having one or more input TFs and being a DEG (Methods section *RNA-seq d*, assuming a False Discovery Rate < 0.05 and absolute log2 fold change (|LFC|) > 0.4248 for $LB_{0.75x}$, > 0.4085 for $LB_{0.5x}$ and > 0.4138 for $LB_{0.25x}$). For each condition, we use the specific values of $\mu_{|LFC|}$ (mean of |LFC|) and number of DEG. The null hypothesis is that there is random association between the two variables. The test rejects the null hypothesis at 0.1 significance level.

| Medium | P-value |
|---|---|
| **$LB_{0.75x}$** | 0.00 |
| **$LB_{0.5x}$** | 0.29 |
| **$LB_{0.25x}$** | 0.85 |

**Table S7. (Related to Figure 4B, and Supplementary Figures S14 and S20B) Statistics of the linear fits in Supplementary Figure S14.** $R^2_{Emp}$, $R^2_{Sorted}$, $Pv^{Emp}$, $Pv^{Shuffled}$ and $Pv^{Sorted}$ values for gene cohorts differing in $K_{TF}$ (number of known input transcription factors (TFs)) (from 1 to 7) and medium ($LB_{0.75x}$, $LB_{0.5x}$ and $LB_{0.25x}$), when testing in Supplementary Figure S14 the correlation between absolute of log2 fold change (|LFC|) of outputs genes and each gene known to express their input TFs.

| | **$LB_{0.75x}$** | | | | |
|---|---|---|---|---|---|
| | **$Pv^{Shuffled}$** | **$Pv^{Emp}$** | **$Pv^{Sorted}$** | **$R^2_{Emp}$** | **$R^2_{Sorted}$** |
| **$K_{TF}$ = 1** | 0.83 | 0.00 | 0.00 | 0.08 | 0.87 |
| **$K_{TF}$ = 2** | 0.74 | 0.03 | 0.00 | 0.01 | 0.95 |
| **$K_{TF}$ = 3** | 0.97 | 0.00 | 0.00 | 0.04 | 0.87 |
| **$K_{TF}$ = 4** | 0.98 | 0.00 | 0.00 | 0.04 | 0.93 |
| **$K_{TF}$ = 5** | 0.81 | 0.00 | 0.00 | 0.03 | 0.91 |
| **$K_{TF}$ = 6** | 1.00 | 0.72 | 0.00 | 0.00 | 0.88 |
| **$K_{TF}$ = 7** | 1.00 | 1.00 | 0.00 | 0.00 | 0.93 |
| | **$LB_{0.5x}$** | | | | |
| | **$Pv^{Shuffled}$** | **$Pv^{Emp}$** | **$Pv^{Sorted}$** | **$R^2_{Emp}$** | **$R^2_{Sorted}$** |
| **$K_{TF}$ = 1** | 0.96 | 0.00 | 0.00 | 0.11 | 0.88 |
| **$K_{TF}$ = 2** | 0.82 | 0.00 | 0.00 | 0.02 | 0.92 |
| **$K_{TF}$ = 3** | 0.63 | 0.00 | 0.00 | 0.06 | 0.88 |
| **$K_{TF}$ = 4** | 0.86 | 0.01 | 0.00 | 0.01 | 0.96 |
| **$K_{TF}$ = 5** | 0.91 | 0.00 | 0.00 | 0.04 | 0.85 |
| **$K_{TF}$ = 6** | 0.97 | 0.00 | 0.00 | 0.03 | 0.88 |
| **$K_{TF}$ = 7** | 0.96 | 0.10 | 0.00 | 0.01 | 0.93 |
| | **$LB_{0.25x}$** | | | | |
| | **$Pv^{Shuffled}$** | **$Pv^{Emp}$** | **$Pv^{Sorted}$** | **$R^2_{Emp}$** | **$R^2_{Sorted}$** |
| **$K_{TF}$ = 1** | 0.85 | 0.00 | 0.00 | 0.06 | 0.93 |
| **$K_{TF}$ = 2** | 0.97 | 0.01 | 0.00 | 0.01 | 0.96 |
| **$K_{TF}$ = 3** | 0.87 | 0.00 | 0.00 | 0.07 | 0.93 |
| **$K_{TF}$ = 4** | 0.80 | 0.68 | 0.00 | 0.00 | 0.89 |
| **$K_{TF}$ = 5** | 0.89 | 0.00 | 0.00 | 0.02 | 0.88 |
| **$K_{TF}$ = 6** | 0.94 | 0.09 | 0.00 | 0.01 | 0.88 |
| **$K_{TF}$ = 7** | 0.87 | 0.93 | 0.00 | 0.00 | 0.88 |

**Table S8. (Related to Figure 4B, and Supplementary Figures S15) Statistics of the linear fits in Supplementary Figure S15.** $R^2_{Emp}$, $R^2_{Sorted}$, $Pv^{Emp}$, $Pv^{Shuffled}$ and $Pv^{Sorted}$ values for gene cohorts differing in $K_{TF}$ (number of known input transcription factors (TFs)) (from 1 to 7) in $LB_{0.5x}$ medium (60 and 125 min), when testing in Supplementary Figure S15 the correlation between absolute of log2 fold change (|LFC|) of outputs genes and each gene known to express their input TFs.

| | **$LB_{0.5x}$ 60 mim** | | | | | **$LB_{0.5x}$ 125 min** | | | | |
|---|---|---|---|---|---|---|---|---|---|---|
| | **$Pv^{Shuffled}$** | **$Pv^{Emp}$** | **$Pv^{Sorted}$** | **$R^2_{Emp}$** | **$R^2_{Sorted}$** | **$Pv^{Shuffled}$** | **$Pv^{Emp}$** | **$Pv^{Sorted}$** | **$R^2_{Emp}$** | **$R^2_{Sorted}$** |
| **$K_{TF}$ = 1** | 0.88 | 0.00 | 0.00 | 0.01 | 0.84 | 0.98 | 0.25 | 0.00 | 0.00 | 0.86 |
| **$K_{TF}$ = 2** | 0.89 | 0.18 | 0.00 | 0.00 | 0.78 | 0.81 | 0.00 | 0.00 | 0.03 | 0.91 |
| **$K_{TF}$ = 3** | 0.95 | 0.25 | 0.00 | 0.00 | 0.64 | 0.72 | 0.91 | 0.00 | 0.00 | 0.96 |
| **$K_{TF}$ = 4** | 0.98 | 0.47 | 0.00 | 0.00 | 0.76 | 0.82 | 0.00 | 0.00 | 0.02 | 0.95 |
| **$K_{TF}$ = 5** | 0.99 | 0.01 | 0.00 | 0.01 | 0.56 | 0.93 | 0.09 | 0.00 | 0.00 | 0.90 |
| **$K_{TF}$ = 6** | 0.97 | 0.01 | 0.00 | 0.03 | 0.38 | 0.99 | 0.98 | 0.00 | 0.00 | 0.88 |
| **$K_{TF}$ = 7** | 0.95 | 1.00 | 0.00 | 0.00 | 0.32 | 0.90 | 0.42 | 0.00 | 0.00 | 0.90 |

**Table S9. (Related to Figure 4B and Supplementary Figure S16) Statistics of the linear fits in Supplementary Figure S16.** $R^2_{Emp}$, $R^2_{Sorted}$, $Pv^{Emp}$, $Pv^{Shuffled}$ and $Pv^{Sorted}$ for results in Supplementary Figure S16, in various media. Data for the 1st, 2nd and 3rd genes (following the transcription start site) of operons of size 3, when confronting the correlation between absolute of LFC, log2 fold change of each of these output gene of the operon and each gene known to express an input transcription factor (TF) common to all 3 genes.

| | **$LB_{0.75x}$** | | | | |
|---|---|---|---|---|---|
| | **$Pv^{Shuffled}$** | **$Pv^{Emp}$** | **$Pv^{Sorted}$** | **$R^2_{Emp}$** | **$R^2_{Sorted}$** |
| **1st gene** | 0.98 | 0.00 | 0.00 | 0.06 | 0.90 |
| **2nd gene** | 0.93 | 0.03 | 0.00 | 0.02 | 0.92 |
| **3rd gene** | 0.98 | 0.00 | 0.00 | 0.04 | 0.93 |
| | **$LB_{0.5x}$** | | | | |
| | **$Pv^{Shuffled}$** | **$Pv^{Emp}$** | **$Pv^{Sorted}$** | **$R^2_{Emp}$** | **$R^2_{Sorted}$** |
| **1st gene** | 0.97 | 0.00 | 0.00 | 0.06 | 0.88 |
| **2nd gene** | 0.98 | 0.00 | 0.00 | 0.05 | 0.85 |
| **3rd gene** | 0.93 | 0.00 | 0.00 | 0.05 | 0.92 |
| | **$LB_{0.25x}$** | | | | |
| | **$Pv^{Shuffled}$** | **$Pv^{Emp}$** | **$Pv^{Sorted}$** | **$R^2_{Emp}$** | **$R^2_{Sorted}$** |
| **1st gene** | 1.00 | 0.00 | 0.00 | 0.07 | 0.88 |
| **2nd gene** | 0.98 | 0.00 | 0.00 | 0.05 | 0.89 |
| **3rd gene** | 0.90 | 0.00 | 0.00 | 0.04 | 0.93 |

**Table S10. (Related to Figure 4B and Supplementary Figure S16) Analysis of Covariance of results in Supplementary Figure S16.** We performed an analysis of covariances, ANCOVA (Methods section *Statistical tests d*) test, to assess if the red fitting lines in Supplementary Figure S16 have the same intercept (I) and/or slope (S). The fitting lines were calculated for the shifts from the control, to assess the correlation between the absolute of log2 fold change of each output gene belonging to an operon of size 3 (i.e., 1$^{st}$, 2$^{nd}$ and 3$^{rd}$ gene in the operon following the transcription start site), and each gene known to express an input TF, common to the 3 genes in the operon. We evaluated if the fitting lines differ between the 1$^{st}$, 2$^{nd}$, and 3$^{rd}$ gene, for each shift. Also, we evaluated if the fitting lines differ between the different shifts, for the 1$^{st}$, 2$^{nd}$, and 3$^{rd}$ gene, respectively. We consider 2 lines statistically different in I and/or S at 0.1 significance level.

| **P-value** | | | | | | |
|---|---|---|---|---|---|---|
| | **$LB_{0.75x}$** | | **$LB_{0.5x}$** | | **$LB_{0.25x}$** | |
| | **I** | **S** | **I** | **S** | **I** | **S** |
| **1$^{st}$ gene vs 2$^{nd}$ gene** | 0.85 | 0.31 | 0.42 | 0.51 | 0.53 | 0.31 |
| **1$^{st}$ gene vs 3$^{rd}$ gene** | 0.96 | 0.54 | 0.77 | 0.38 | 1.00 | 0.12 |
| **2$^{nd}$ gene vs 3$^{rd}$ gene** | 0.81 | 0.67 | 0.54 | 0.82 | 0.43 | 0.56 |
| | **1$^{st}$ gene** | | **2$^{nd}$ gene** | | **3$^{rd}$ gene** | |
| | **I** | **S** | **I** | **S** | **I** | **S** |
| **$LB_{0.75x}$ vs $LB_{0.5x}$** | 0.12 | 0.75 | 0.30 | 0.83 | 0.14 | 0.66 |
| **$LB_{0.75x}$ vs $LB_{0.25x}$** | 0.06 | 0.72 | 0.00 | 0.94 | 0.03 | 0.33 |
| **$LB_{0.5x}$ vs $LB_{0.25x}$** | 0.63 | 0.94 | 0.03 | 0.69 | 0.37 | 0.45 |

**Table S11. (Related to Figure 4B and Supplementary Figure S17) Statistics of the linear fits in Supplementary Figure S17.** $R^2_{Emp}$, $R^2_{Sorted}$, $Pv^{Emp}$, $Pv^{Shuffled}$ and $Pv^{Sorted}$ values for results in Supplementary Figure S17, in various shifts. Data for the 1st, 2nd and 3rd gene (following the transcription start site) belonging to a Transcription Unit (TU) of size 3, when confronting the correlation between the absolute LFC of each output gene of the TU and each gene expressing an input TF common to all 3 genes.

| | $LB_{0.75x}$ | | | | |
|---|---|---|---|---|---|
| | $Pv^{Shuffled}$ | $Pv^{Emp}$ | $Pv^{Sorted}$ | $R^2_{Emp}$ | $R^2_{Sorted}$ |
| **1st gene** | 1.00 | 0.00 | 0.00 | 0.05 | 0.89 |
| **2nd gene** | 0.91 | 0.01 | 0.00 | 0.02 | 0.91 |
| **3rd gene** | 0.96 | 0.00 | 0.00 | 0.03 | 0.90 |
| | $LB_{0.5x}$ | | | | |
| | $Pv^{Shuffled}$ | $Pv^{Emp}$ | $Pv^{Sorted}$ | $R^2_{Emp}$ | $R^2_{Sorted}$ |
| **1st gene** | 0.95 | 0.00 | 0.00 | 0.05 | 0.84 |
| **2nd gene** | 0.96 | 0.00 | 0.00 | 0.05 | 0.81 |
| **3rd gene** | 0.79 | 0.00 | 0.00 | 0.04 | 0.88 |
| | $LB_{0.25x}$ | | | | |
| | $Pv^{Shuffled}$ | $Pv^{Emp}$ | $Pv^{Sorted}$ | $R^2_{Emp}$ | $R^2_{Sorted}$ |
| **1st gene** | 0.82 | 0.00 | 0.00 | 0.04 | 0.88 |
| **2nd gene** | 0.97 | 0.00 | 0.00 | 0.04 | 0.86 |
| **3rd gene** | 0.88 | 0.02 | 0.00 | 0.02 | 0.92 |

**Table S12. (Related to Figure 4B and Supplementary Figure S17). Analysis of Covariance of results in Supplementary Figure S17.** Analysis of covariances, ANCOVA (Methods section *Statistical tests d*) to test if the red fitting lines in Supplementary Figure S17 have the same intercept (I) and/or slope (S). Fitting lines calculated for the shifts from the control, to assess the correlation between the absolute of LFC, log2 fold change of each output gene belonging to a Transcription Unit (TU) of size 3 (i.e., $1^{st}$, $2^{nd}$ and $3^{rd}$ gene in the TU following the transcription start site), and the |LFC| of each gene known to express an input TF common to the 3 genes in the TU. We evaluate if the fitting lines differ between the $1^{st}$, $2^{nd}$, and $3^{rd}$ gene, for each shift. Also, we evaluate if the fitting lines differ between the different shifts, for the $1^{st}$, $2^{nd}$, and $3^{rd}$ gene, respectively. We consider 2 lines to be statistically different in I and/or S at 0.1 significance level.

| **P-value** | | | | | | |
|---|---|---|---|---|---|---|
| | **$LB_{0.75x}$** | | **$LB_{0.5x}$** | | **$LB_{0.25x}$** | |
| | **I** | **S** | **I** | **S** | **I** | **S** |
| **$1^{st}$ gene vs $2^{nd}$ gene** | 0.94 | 0.46 | 0.64 | 1.00 | 0.72 | 0.98 |
| **$1^{st}$ gene vs $3^{rd}$ gene** | 0.79 | 0.69 | 0.88 | 0.72 | 0.61 | 0.32 |
| **$2^{nd}$ gene vs $3^{rd}$ gene** | 0.86 | 0.73 | 0.52 | 0.72 | 0.89 | 0.30 |
| | **$1^{st}$ gene** | | **$2^{nd}$ gene** | | **$3^{rd}$ gene** | |
| | **I** | **S** | **I** | **S** | **I** | **S** |
| **$LB_{0.75x}$ vs $LB_{0.5x}$** | 0.09 | 0.42 | 0.23 | 1.00 | 0.09 | 0.50 |
| **$LB_{0.75x}$ vs $LB_{0.25x}$** | 0.00 | 0.20 | 0.00 | 0.66 | 0.00 | 0.12 |
| **$LB_{0.5x}$ vs $LB_{0.25x}$** | 0.18 | 0.52 | 0.03 | 0.52 | 0.06 | 0.21 |

**Table S13. (Related to Supplementary Figures S20C and S21) Analysis of Covariance of results in Supplementary Figure S21.** Analysis of covariances (ANCOVA, Methods section *Statistical tests d*) to assess if the fitting lines in Supplementary Figure S21 have the same intercept (I) and/or slope (S). We evaluate, as a function of $K_{TF}$ (number of input transcription factors), the chance of correlation between |LFC| and the shift in mean RNA Polymerase (RNAP) concentration. We consider 2 lines to be statistically different in I and/or S if P-value < 0.1.

| **P-value** | | | | | | | | | | | | | | | | |
|---|---|---|---|---|---|---|---|---|---|---|---|---|---|---|---|---|
| | **$K_{TF}$ = 0** | | **$K_{TF}$ = 1** | | **$K_{TF}$ = 2** | | **$K_{TF}$ = 3** | | **$K_{TF}$ = 4** | | **$K_{TF}$ = 5** | | **$K_{TF}$ = 6** | | **$K_{TF}$ = 7** | |
| | **I** | **S** | **I** | **S** | **I** | **S** | **I** | **S** | **I** | **S** | **I** | **S** | **I** | **S** | **I** | **S** |
| **$K_{TF}$ = 0** | 1.00 | 1.00 | | | | | | | | | | | | | | |
| **$K_{TF}$ = 1** | 0.00 | 0.00 | 1.00 | 1.00 | | | | | | | | | | | | |
| **$K_{TF}$ = 2** | 0.00 | 0.00 | 0.00 | 0.00 | 1.00 | 1.00 | | | | | | | | | | |
| **$K_{TF}$ = 3** | 0.00 | 0.00 | 0.00 | 0.00 | 0.12 | 0.19 | 1.00 | 1.00 | | | | | | | | |
| **$K_{TF}$ = 4** | 0.00 | 0.00 | 0.00 | 0.02 | 0.65 | 0.64 | 0.14 | 0.18 | 1.00 | 1.00 | | | | | | |
| **$K_{TF}$ = 5** | 0.00 | 0.00 | 0.00 | 0.00 | 0.00 | 0.00 | 0.01 | 0.02 | 0.00 | 0.00 | 1.00 | 1.00 | | | | |
| **$K_{TF}$ = 6** | 0.00 | 0.00 | 0.00 | 0.00 | 0.20 | 0.26 | 0.70 | 0.69 | 0.11 | 0.15 | 0.24 | 0.31 | 1.00 | 1.00 | | |
| **$K_{TF}$ = 7** | 0.00 | 0.00 | 0.00 | 0.00 | 0.01 | 0.03 | 0.13 | 0.17 | 0.00 | 0.01 | 0.85 | 0.89 | 0.24 | 0.30 | 1.00 | 1.00 |

**Table S14. (Related to Figure 4D and Supplementary Figures S24C-S24F) Statistical tests between the distributions in Supplementary Figures S24C-S24F.** P-values from the 2-sample T-test and 2-sample KS-test confronting the distributions and the Z-test (Methods section *Statistical tests a*). For the z-test, the mean and standard deviation are from the distribution of genes without input transcription factors (TFs), $K_{TF}$ = 0. The data is from the shift to $LB_{0.25x}$.

**P-value, 2-sample T-test**

| | **$K_{TF}$ = 0** | **$K_{TF}$ = 1, CRP** | **$K_{TF}$ = 1, FIS** | **$K_{TF}$ = 2, FNR & ArcA** |
|---|---|---|---|---|
| **$K_{TF}$ = 0** | 1.00 | | | |
| **$K_{TF}$ = 1, CRP** | 0.88 | 1.00 | | |
| **$K_{TF}$ = 1, FIS** | 0.00 | 0.02 | 1.00 | |
| **$K_{TF}$ = 2, FNR & ArcA** | 0.06 | 0.08 | 0.00 | 1.00 |

**P-value, 2-sample KS-test**

| | **$K_{TF}$ = 0** | **$K_{TF}$ = 1, CRP** | **$K_{TF}$ = 1, FIS** | **$K_{TF}$ = 2, FNR & ArcA** |
|---|---|---|---|---|
| **$K_{TF}$ = 0** | 1.00 | | | |
| **$K_{TF}$ = 1, CRP** | 0.64 | 1.00 | | |
| **$K_{TF}$ = 1, FIS** | 0.00 | 0.00 | 1.00 | |
| **$K_{TF}$ = 2, FNR & ArcA** | 0.10 | 0.50 | 0.00 | 1.00 |

| **P-value, Z-test** | |
|---|---|
| **$K_{TF}$ = 0** | 1.00 |
| **$K_{TF}$ = 1, CRP** | 0.89 |
| **$K_{TF}$ = 1, FIS** | 0.00 |
| **$K_{TF}$ = 2, FNR & ArcA** | 0.16 |

**Table S15. (Related to Supplementary Figures S26A and S26C) Analysis of the potential role of each global regulatory in the genome-wide shifts in RNA abundances.**

| | |
|---|---|
| rpoD ($\sigma^{70}$) | Likely not influential due to lack of changes in its concentration. |
| rpoN ($\sigma^{54}$) | Likely not influential due to lack of changes in its concentration in 2 of 3 perturbations. |
| rpoS ($\sigma^{38}$) | Likely not influential since it follows the RNAP changes (Supplementary Figure S5). |
| rpoH ($\sigma^{34}$) | Likely not influential due to lack of changes in its concentration in the first perturbation and the inconsistency with changes in the LFCs of its outputs (Supplementary Figure S26B). |
| fliA ($\sigma^{28}$) | Likely not influential due to lack of changes in its concentration in 2 of 3 perturbations. |
| rpoE ($\sigma^{24}$) | Likely not influential since, while it responded to 2 of 3 perturbations, its response strength was inconsistent with the perturbation strengths. Also, its outputs responded inconsistently. |
| fecI ($\sigma^{19}$) | Likely not influential since it did not respond to any perturbation. |
| ihfA | Likely not influential since its outputs did not respond consistently. |
| ihfB | Likely not influential due to lack of change in its concentration with the perturbations. |
| fnr | Likely not influential due to lack of change in its concentration with the perturbations. |
| arcA | Likely not influential since its outputs did not respond consistently to either arcA or RNAP. |
| fis | Likely not influential due to lack of change in its concentration in 2 of 3 perturbations. |
| lrp | Likely not influential due to lack of change in its concentration with the perturbations. |
| crp | Likely not influential due to lack of change in its concentration with the perturbations. |
| narL | Likely not influential due to lack of change in its concentration in 2 of 3 perturbations. |
| flhC | Likely not influential as it followed the RNAP changes and only controls 32 genes. |
| flhD | Likely not influential as it followed the RNAP changes and only controls 32 genes. |
| fur | Likely not influential due to lack of change in its concentration with the perturbations. |
| hns | Likely not influential due to lack of change in its concentration with the perturbations. |

**Table S16. (Related to Figures 5A and 5B and Supplementary Figure S12) Numbers of genes with input transcription factors (TFs) with a given overall bias.** Number of genes with a given absolute of the sum of the regulatory effects, ' $r$ ' of inputs ($|b|$) from 0 to 5. Calculated from data from RegulonDB.

| $\lvert \mathbf{b} \rvert$ | **0** | **1** | **2** | **3** | **4** | **5** |
|---|---|---|---|---|---|---|
| **No. genes** | 2598 | 996 | 305 | 107 | 27 | 8 |

**Table S17. (Related to Figure 5A) Number of genes with all input transcription factors (TFs) having the same regulatory effect (‘*r*’).** Number of genes whose absolute sum of the $r$ of inputs ($|b|$) equals their number of input TFs, $K_{TF}$ (for $K_{TF}$ = 0 to 7). As $K_{TF}$ increases, the fraction of genes with $|b|$ = $K_{TF}$ decreases.

| $K_{TF}$ | No. genes | No. genes with $|b|$ = $K_{TF}$ |
|---|---|---|
| 0 | 2343 | 2343 |
| 1 | 703 | 686 |
| 2 | 389 | 169 |
| 3 | 260 | 61 |
| 4 | 133 | 19 |
| 5 | 115 | 8 |
| 6 | 37 | 0 |
| 7 | 27 | 0 |

**Table S18. (Related to Supplementary Figure S31) Analysis of Covariance of results in Supplementary Figure S31.** We performed an analysis of covariances, ANCOVA (Methods section *Statistical tests d*) assessing if the fitting lines in Supplementary Figure S31 have the same intercept (I) and/or slope (S). The fitting lines were calculated for the shifts from the control ($LB_{1.0x}$) to other media, to assess the correlation between the mean of |LFC| (absolute of LFC, log2 fold change), $\mu_{|LFC|}$, and the mean of $|b|$ (absolute of the sum of the regulatory effects (' $r$ ') of inputs), for each cohort of genes with a specific $K_{TF}$ (number of input transcription factors (TFs), from 0 to 5). We consider that 2 lines have statistically different I and/or S at a 0.1 significance level.

| P-value | | | | | | |
|---|---|---|---|---|---|---|
| | **$LB_{0.75x}$** | | **$LB_{0.5x}$** | | **$LB_{0.25x}$** | |
| | **I** | **S** | **I** | **S** | **I** | **S** |
| **$LB_{0.75x}$** | 1.00 | 1.00 | | | | |
| **$LB_{0.5x}$** | 0.32 | 0.56 | 1.00 | 1.00 | | |
| **$LB_{0.25x}$** | 0.08 | 0.71 | 0.23 | 0.76 | 1.00 | 1.00 |

**Table S19. Statistics of various features of the TFN topology of *E. coli*.** Shown are the coefficients of determination ($R^2$s) and corresponding p-values of statistical significance of a linear fit between the |LFC| (absolute log2 fold change) of each gene and its corresponding topological feature (defined in (14), Methods section *Transcription Factor Network of Escherichia coli*). We reject the null hypothesis that the data is best fit by a horizontal line at 0.1 significance level.

| | $LB_{0.75x}$ | | $LB_{0.5x}$ | | $LB_{0.25x}$ | |
|---|---|---|---|---|---|---|
| **Topological feature** | **P-value** | $R^2$ | **P-value** | $R^2$ | **P-value** | $R^2$ |
| **Avg. short path length** | 0.72 | 0.00 | 0.42 | 0.00 | 0.50 | 0.00 |
| **Betweenness centrality** | 0.48 | 0.00 | 0.14 | 0.00 | 0.50 | 0.00 |
| **Stress centrality** | 0.41 | 0.00 | 0.13 | 0.00 | 0.15 | 0.00 |
| **Clustering coefficient** | 0.00 | 0.01 | 0.00 | 0.01 | 0.00 | 0.01 |
| **Eccentricity** | 0.54 | 0.00 | 0.65 | 0.00 | 0.79 | 0.00 |
| **Edge-count** | 0.26 | 0.00 | 0.27 | 0.00 | 0.73 | 0.00 |
| **Out-degree** | 0.55 | 0.00 | 0.39 | 0.00 | 0.19 | 0.00 |
| **Neighborhood connectivity** | 0.87 | 0.00 | 0.91 | 0.00 | 0.34 | 0.00 |

**Table S20. Statistics of the mean behavior of the genes responsive to (p)ppGpp (15).** Shown are the p-values from the 2-sample T-test (Methods section *Statistical tests a*) between the mean behavior of log2 fold change (LFC) of genes responsive to (p)ppGpp (15) and the LFC of all genes analyzed by RNA-seq.

| | $LB_{0.25X}$ | $LB_{0.5X}$ 125 min | $LB_{0.5X}$ 180 min | $LB_{0.75X}$ | $LB_{1.5X}$ | $LB_{2.0X}$ | $LB_{2.5X}$ |
|---|---|---|---|---|---|---|---|
| **After 5 min** | 0.18 | 0.00 | 0.87 | 0.29 | 0.00 | 0.00 | 0.00 |
| **After 10 min** | 0.53 | 0.00 | 0.01 | 0.02 | 0.39 | 0.11 | 0.41 |

**Table S21. Statistics of the behavior of sRNA.** Shown are the p-values from a 2-sample T-test (Methods section *Statistical tests a*) between the mean log2 fold change (LFC) of sRNAs and the LFC of all genes analyzed by RNA-seq.

| P-value, 2-sample T-test | | | | | | | |
|---|---|---|---|---|---|---|---|
| | **$LB_{0.25X}$** | **$LB_{0.5X}$ 125 min** | **$LB_{0.5X}$ 180 min** | **$LB_{0.75X}$** | **$LB_{1.5X}$** | **$LB_{2.0X}$** | **$LB_{2.5X}$** |
| **sRNA (93 genes)** | 0.39 | 0.1 | 0.32 | 0.13 | 0.23 | 0.24 | 0.40 |
| **Number of DEG rRNAs** | | | | | | | |
| **sRNA** | 18 DEG | 4 DEG | 7 DEG | 0 | 33 DEG | 33 DEG | 30 DEG |

**Table S22. Statistics of the behavior of rRNAs.** Shown, for each shift, are the p-values from the 2-sample T-test (Methods section *Statistical tests a*) between the mean log2 fold change (LFC) of rRNAs and the LFC of all genes analyzed by RNA-seq. Also shown is the number of DEG genes.

| P-value, 2-sample T-test | | | | | | | |
|---|---|---|---|---|---|---|---|
| | $LB_{0.25X}$ | $LB_{0.5X}$ 125 min | $LB_{0.5X}$ 180 min | $LB_{0.75X}$ | $LB_{1.5X}$ | $LB_{2.0X}$ | $LB_{2.5X}$ |
| **rRNA (22 genes)** | 0.00 | 1.00 | 0.00 | 0.00 | 0.00 | 0.00 | 0.00 |
| **Number of DEG rRNAs** | | | | | | | |
| **rRNA** | 0 | 0 | 0 | 0 | 15 DEG | 0 | 2 DEG |
